# High-throughput mapping of spontaneous mitotic crossover and genome instability events with sci-L3-Strand-seq

**DOI:** 10.1101/2025.05.19.654945

**Authors:** Peter Chovanec, Trevor Ridgley, Yi Yin

**Affiliations:** Department of Human Genetics, David Geffen School of Medicine, UCLA, Los Angeles, CA, 90095, USA

## Abstract

Despite the many advances in single cell genomics, detecting structural rearrangements in single cells, particularly error-free sister-chromatid exchanges, remains challenging. Here we describe sci-L3-Strand-seq, a combinatorial indexing method with linear amplification for DNA template strand sequencing that cost-effectively scales to millions of single cells, as a platform for mapping mitotic crossover and resulting genome instability events. We provide a computational framework to fully leverage the throughput, as well as the relatively sparse but multifaceted genotype information within each cell that includes strandedness, digital counting of copy numbers, and haplotype-aware chromosome segmentation, to systematically distinguish seven possible types of mitotic crossover outcomes. We showcase the power of sci-L3-Strand-seq by quantifying the rates of error-free and mutational crossovers in thousands of cells, enabling us to explore enrichment patterns of genomic and epigenomic features. The throughput of sci-L3-Strand-seq also gave us the ability to measure subtle phenotypes, opening the door for future large mutational screens. Furthermore, mapping clonal lineages provided insights into the temporal order of certain genome instability events, showcasing the potential to dissect cancer evolution. Altogether, we show the wide applicability of sci-L3-Strand-seq to the study of DNA repair and structural variations.

## Introduction

DNA double stranded breaks (DSB) pose a major risk to genome integrity. During the S and G2 phase of the cell cycle, the use of homologous DNA from sister chromatids or homologs facilitates DSB repair through homologous recombination (HR). After resection of the DSB ends, HR can initiate repair through single-ended invasion via synthesis-dependent strand annealing (SDSA) or break-induced replication (BIR), or in combination with second-end capture produce a Holliday junction (HJ) intermediate that can undergo dissolution or resolution. Dissolution of a double HJ leads to a non-crossover (NCO), while resolution can produce both NCO and crossover (CO) outcomes (1, 2). Despite the error-free nature of HR, COs can lead to large-scale genome rearrangements that pose a risk to the stability of the genome. In mammalian cells, repair by end joining, including non-homologous end joining (NHEJ) and microhomology-mediated end joining (MMEJ), can also lead to genome rearrangements. NHEJ can occur throughout the cell cycle, while MMEJ favors S/G2 in competition with HR. However, the resulting mutations by end joining are often local to the DSBs, with small insertions or deletions asserting its physiological impact in exons or splicing sites. Large scale rearrangements caused by end joining often require two DSBs, which are not as frequent spontaneously without exogenous damage-inducing agents. Mapping of germline structure variations (SV) in the human population implicates HR as the major contributing source (3), while a majority of large-scale DNA rearrangements in cancer genomes do not display signatures of homology-dependent recombination (4). However, spontaneous SVs during normal development have not been intensively studied and require high-throughput, unbiased mapping methods given their rare occurrence. In particular, pre-meiotic mitotic divisions could also contribute to germline SVs observed in the population.

We have previously categorized genomic rearrangements produced from mitotic COs into seven classes that can be broadly divided into events arising from allelic HR and non-allelic HR (NAHR). Meanwhile, unresolved HR intermediates could also affect proper chromosome segregation (5). Briefly, allelic HR can generate error-free sister chromatid exchanges (SCE, class 1) (Fig 1A and FigS1B), or large-scale copy-neutral loss-of-heterozygosity (cnLOH, class2) (Fig1B) if parental homolog was used as repair template. With the proximity of the newly replicated sister, the bias of using the sister chromatid is over 100-fold compared to using the parental homolog harboring single-nucleotide variations (SNVs) (6). The high prevalence of repeats in mammalian genomes provides an alternative source of homology that can result in rearrangements between non-allelic sequences. NAHR between intra-chromatid repeats can give rise to copy number variations (CNV, class 3) (Fig1C and FigS1C) and inversions (class 4) (Fig1D); while those between inter-chromosomal repeats can result in translocations (class 5) (Fig1E). Repair by end joining with two initiating DSBs are also likely the source for CNV, inversions and translocations (FigS1E). Lastly, unresolved HR intermediates could give rise to aneuploidy (class 6) (Fig1F and FigS1D) and uniparental disomy (UPD, class 7) (Fig1G). The exact relationship between crossover and whole-chromosome alterations has not been fully-established. Chua and Jinks-Robertson (7) found increased chromosome loss following a mitotic crossover event in yeast, possibly due to increased nondisjunction. UPD may be formed either through an aneuploid intermediate resulting from nondisjunction, or through meiosis-like homolog separation (8, 9) in a single division. Altogether, the ability to detect a wide variety of repair outcomes genome-wide without bias is key to understanding the full complexity of mitotic crossover.

**Figure 1:**
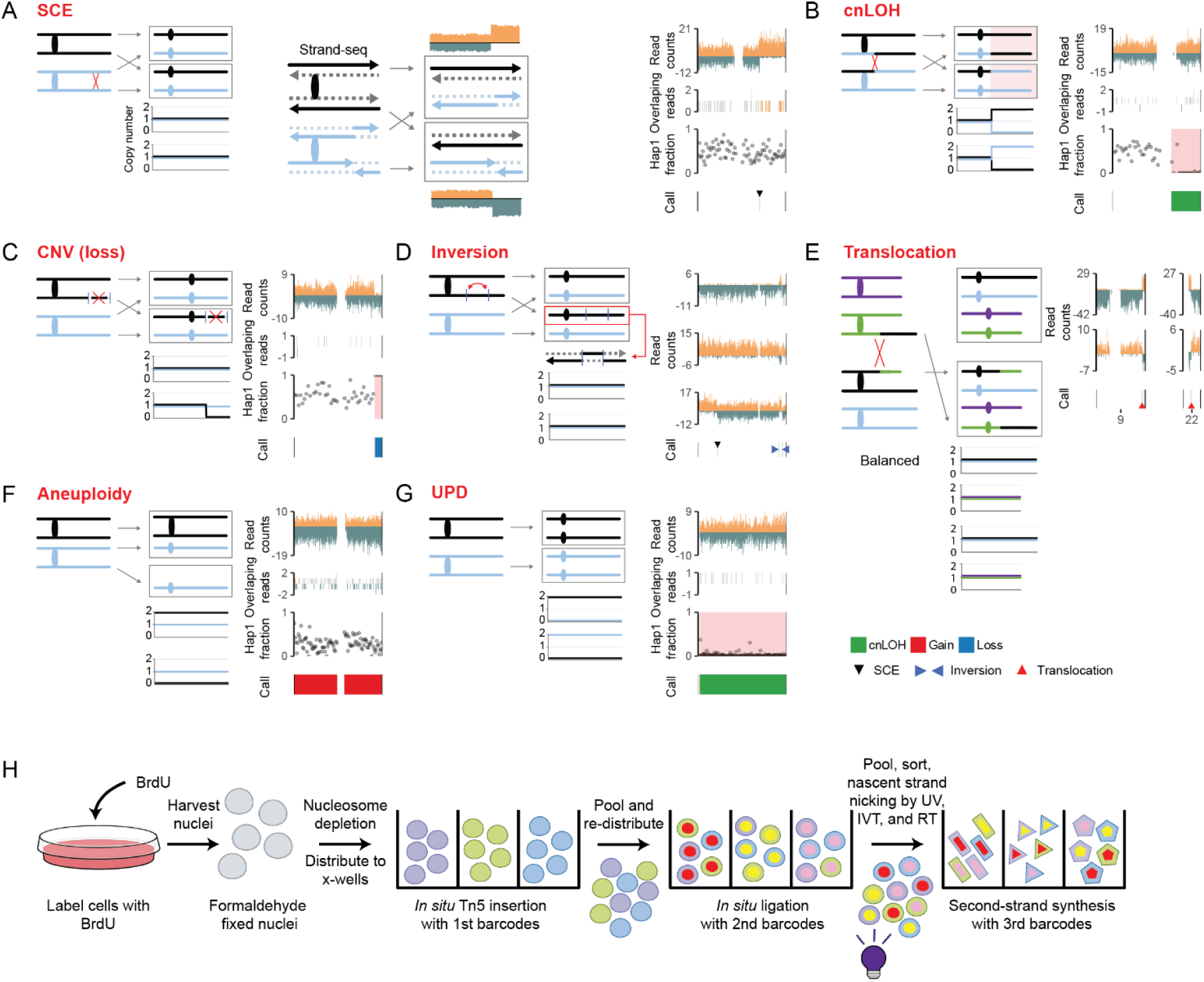
sci-L3-Strand-seq enables the detection of all seven mitotic crossover outcomes. (A-G) Schematics of the seven types of mitotic crossover outcomes with example sci-L3-Strand-seq data. Homologs with SNVs are shown in black and blue (green and purple for translocation), each containing two identical sister chromatids. Oval indicates centromere. Read counts in the example data show Watson (W) reads colored dark yellow and Crick (C) reads colored dark green. This color code for strand directionality is used throughout this paper unless noted otherwise. Overlapping reads in the example data show the locations of overlaps, with nearby locations offset on the y-axis. Each location belongs to one of three categories based on the type of overlap: dark green represents overlapping C strand reads, dark yellow represents overlapping W strand reads, and gray represents overlapping reads of opposite strands. Each point for the haplotype1 (hap1) fraction is the phased allelic fraction of approximately 50 binned SNVs for Patski and 15 for BJ-5ta. LOH calls are shaded in pink. **(A) Sister chromatid exchanges (SCE)**. Both the chromosome heterozygosity and copy number remain unchanged after SCE. Only a strand-switch on the replication template strand (solid arrows) can be detected by Strand-seq. Without an SCE, the template strands map continuously to either Watson or Crick (black homolog). With an SCE, the template strands are exchanged. In the example, SCEs are marked by a black triangle in the “Call” track. **(B) Copy neutral loss of heterozygosity (cnLOH).** Inter-homolog (IH) crossover leads to reciprocal cnLOH (shaded in pink) in the daughter cells. Note that with reciprocal events, if both daughter cells are sequenced together with bulk WGS, every position will appear heterozygous given the 1:1 ratio of the two alleles. **(C) Copy number variation (CNV).** Insertion or deletions between intra-chromatid sequences with high similarity as an example, resulting from non-allelic homologous recombination (NAHR). **(D) Inversions.** Intra-chromatid reversal of sequences end-to-end, for example between repeats at the two vertical bar, results in no change in chromosome heterozygosity or copy number. Just like with SCE, a double strand-switch on the replication template strand (two inward facing blue triangles) can be detected by Strand-seq. However, confident identification of inversions requires the detection of a shared strand state across multiple cells that negatively correlate with the stand state of the adjoining regions. **(E) Translocations.** Exchange of sequence between non-homologous chromosomes. Translocations can be unbalanced, where recombined chromatids segregate apart and result in the loss or gain of chromosomal segments in the daughter cells (not shown); or balanced, where the co-segregation of recombined chromatids results in an equal exchange of chromosomal segments with no change in copy number of heterozygosity. Similar to inversions, the identification of balanced translocations requires the detection of a shared strand state across multiple cells that positively correlates (as opposed to negatively correlating in inversions) between the translocated chromosomes. In black and blue, green and purple are the homologs of two different chromosomes. **(F) Aneuploidy.** Aberrant crossovers can cause nondisjunction, resulting in monosomy and trisomy in the reciprocal daughter cells. Example data (trisomy) is shown for only one of the daughter cells. **(G) Uniparental disomy (UPD).** Reciprocal UPD is shown with example data for only one of the daughter cells. Other routes leading to UPD include the duplication of a monosomy or the loss of the non-allelic chromosome from a trisomy. **(H) sci-L3-Strand-seq workflow overview.** IVT: in-vitro transcription; RT: reverse transcription. See FigS2.

To enable the study of these genomic rearrangements, we recently developed a set of cost-effective methodologies that combine single-cell combinatorial indexing (sci) with reduced amplification bias using linear amplification (L), and 3-level barcoding for increased throughput called sci-L3 (10).The principal method, sci-L3, enables single-cell whole genome sequencing (WGS) for thousands of cells and addresses several limitations of methods widely used for the study of mitotic CO outcomes. The few studies examining mitotic CO outcomes genome-wide rely on bulk whole genome sequencing (WGS) of subcloned individual cells (11–14). The need to manually clone daughter cells is labor intensive, intractable for scaling to large sets of mutants, and unfeasible for *in vivo* studies. Sci-L3-WGS can directly map reciprocal cnLOH and UPD events in single cells, including *in vivo* that otherwise requires clonal expansion (15), and has the potential to scale to large sets of mutants. Despite the advances gained with sci-L3-WGS, the 99% of mitotic COs that constitute error-free SCEs still cannot be detected at all. Moreover, the detection of inversions and balanced translocation is still reliant on junction-spanning reads that require a high sequencing depth (Fig S1A).

The semi-conservative nature of DNA replication facilitates the differential incorporation of the thymidine analog bromodeoxyuridine (BrdU) between sister chromatids. After one cell division with BrdU, the labeling in daughter cells enables the detection of strand exchanges between recombined sister chromatids or homologs (16, 17). These concepts have been applied in a sequencing-based approach called Strand-seq, which allows the genome-wide detection of SCEs at high resolution (Fig1A) (18).

Strand-seq has seen many additional applications such as the improved detection and genotyping of inversions and some of the other SVs (Fig1C-F) (19–21), and the construction of chromosome level haplotype-phased genome assemblies (22, 23). However, Strand-seq is constrained by: 1) a very low throughput that leads to 2) a high cost per cell, 3) and the reliance on bias-prone PCR amplification (18, 24). The development of OP-Strand-seq improved throughput but requires specialized equipment not commonly available (25). The chemistry in the original and the OP-Strand-seq uses MNase, which prevents chromosome fragmentation pattern-based “digital” information for counting chromosome copy numbers, while transposon-based chromosome fragmentation uniquely enables digital counting (26).

Such digital copy number counts can be further used in a strand-specific and haplotype-aware fashion, which substantially improves analyses of various types of SVs with low coverage.

### Design

With the sci-L3-Strand-seq methodology we provide both experimental and computational advances that together establish a unifying framework for mapping all the seven types of mitotic crossover outcomes at scale. Experimentally, we extended the sci-L3 barcoding and linear amplification scheme to Strand-seq, which requires: 1) successful ablation of the nascent strand by employing a novel enzymatic step (27); and 2) an amplification chemistry that fully preserves strand information. Computationally, we tackled the trade-off between throughput and coverage per cell. Our computational framework 1) at a single cell level fully integrates relatively sparse but multifaceted genotype information by using strandedness, haplotype, digital counting of copy numbers, and their combinations for both chromosome segmentation and systematically annotate different types of mitotic crossover outcomes; and 2) at a population level fully leverages the throughput to dissect subclonal complex rearrangement involving inversions, translocations in duplicated regions. Conceptually, we show the possibility of interpreting the underlying mechanisms for some of the genome instability events given the observed subclonal structure.

## Results

### Expansion of sci-L3 to Strand-seq

Recombination between identical sister chromatids does not change the DNA sequence and is thus missed in short- and long-read WGS. Strand-seq only sequences the template strands of replication in the next G1 post cell division, enabling mapping of error-free SCE (18, 24). To integrate Strand-seq with the sci-L3 scheme (Fig1H) requires ablation of BrdU labeled nascent strands while maintaining strand directionality in the whole-genome amplification process. We summarize key deviations from the original Strand-seq (24): 1) nuclei are formaldehyde fixed to enable the split-and-pool barcoding strategy; 2) Tn5 transposome is used instead of MNase for genome fragmentation and first-round barcoding. The single-stranded strand transfer activity of Tn5 transposase preserves template strand directionality, and uniquely enables digital counting of DNA copy number (26) with low per cell coverage; 3) sci-L3 employs sticky-end ligation of the second-round barcode with the T7 promoter in a hairpin loop to improve ligation efficiency and to minimize amplification of non-genomic DNA if the T7 promoter sequence were supplied in duplex. These features further ensures subsequent unidirectional linear amplification with *in-vitro* transcription (IVT) (FigS2); 4) notably, we found that an additional enzymatic step with Uracil DNA Glycosylase (UDG) and endonuclease VIII (EndoVIII) is required to ablate the incorporated BrdU (27).

These novel additions to both the sci-L3 and Strand-seq not only produced Strand-seq libraries at scale, but pose opportunities for computational method development. To tackle the throughput vs. coverage trade-off, we typically build iterative processes that go through cycles of: 1) leveraging population information for a large number of cells (e.g., phasing: assigning heterozygous SNVs to their parental homologs by strandedness of the reads); 2) integrating multifaceted genotype information (strandedness, newly phased haplotype, digital counting of copy numbers, and *<u>their combinations</u>*) at a single cell level, 3) identifying subclonal complex SVs, again at a population level; and 4) revisiting event type annotation at a single cell level given the subclonal information. For example, the limited throughput of Strand-seq typically renders phasing a stand-alone effort (23, 28) from calling SVs. Here, we provide examples of an iterative process involving phasing, calling SV, and refining phasing. Additionally, mutational SVs (like deletions and inversions) could also involve strand switches, and are sometimes indistinguishable from SCE. With our increased number of cells analyzed, we identified “twin hotspots” in SCE profiles actually reflecting underlying subclonal inversions. We present our computational toolkit in the order of the seven types of mitotic crossover outcomes below, but highlight where these analyses crosstalk.

We piloted our sci-L3-Strand-seq analysis framework in three cell lines. HAP1 is a human haploid cell line that is extensively used for genome engineering and is an ideal model for future mutational screens (29, 30). We have a clear-cut expectation of observing only Watson (W; positive strand) or Crick (C; negative strand) reads within any given region if originating only from the template strand, and we can accurately assess background levels and hence library quality as a result (see Supplemental Text and Methods).

Patski is a mouse female interspecific F1 (C57BL/6J x *M. spretus*) cell line with high quality SNVs published (31, 32) and ground truth for benchmarking phasing with large cell number but sparse coverage, as well as for exploring adjustment to phasing methods with pre-existing subclonal LOH events. Lastly, we examined BJ-5ta, a stable human male line derived from neonatal tissue without any karyotypic abnormalities (10, 26), but unphased. Commercial and patient-derived human cell lines are typically unphased, yet phasing is required for identifying cnLOH. Collectively, these cell lines are ideal for assessing sci-L3-Strand-seq library quality, benchmarking the ability to phase haplotypes and identify SVs, and using sci-L3-Strand-seq as a unifying framework for crossover outcome analyses.

### Sci-L3-Strand-seq captures the replication template strands and identifies SCEs

In HAP1 cells, aside from the known disomic region on chr15 (33), sci-L3-Strand-seq generates clear strand specific profiles and strand switches with a median background estimate of 4.9% (meaning 4.9 C reads for every 100 W reads in a W chromosome, FigS3A-B). Importantly, our ability to sequence DNA template strands showed all the enzymatic steps in sci-L3 preserve strand information (Fig2A).

**Figure 2:**
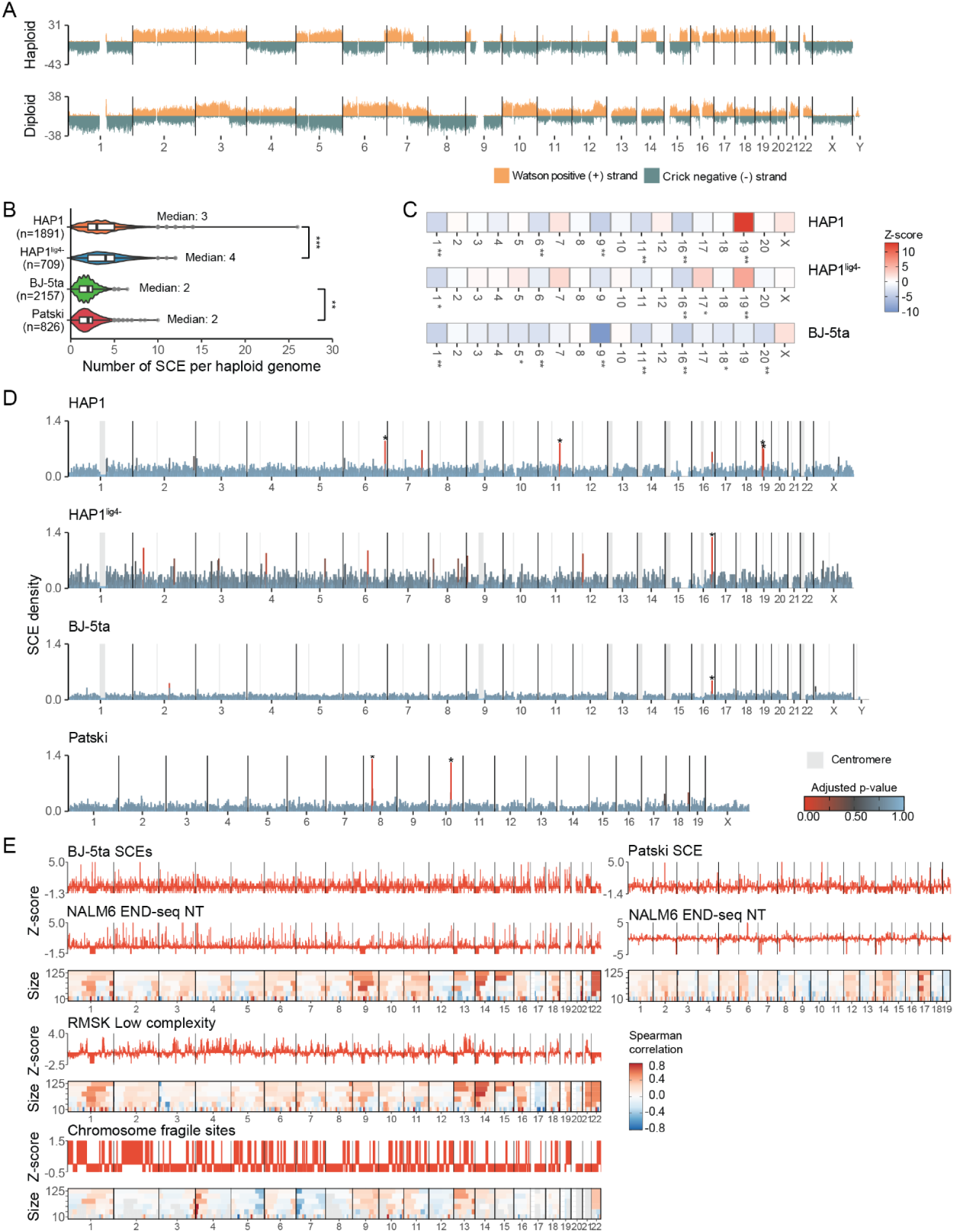
Sister chromatid exchange rate and genomic hotspots.6. **(A) sci-L3-Strand-seq example data.** Top panel shows a haploid human cell (HAP1; median reads per Mb: 97, about 2.1% coverage), while the bottom panel shows a diploid human (BJ-5ta; median reads per Mb: 99) cell. Reads were counted across 200 Kb bins. **(B) Number of SCE per haploid genome for each cell line**. Significant difference level (*** < 0.001, ** <0.01, * <0.05) calculated using Mann-Whitney U, HAP1 vs HAP1^lig4-^ (BH adjusted p-value: 0.00038; MAD for HAP1 1.48, HAP1^lig4-^ 1.48) and BJ-5ta vs Patski (BH adjusted p-value: 0.003; MAD for BJ-5ta 0.74, Patski 1.48). Cells with a high number of loss or gain annotations were excluded. **(C) SCE centromere depletion and enrichment.** The permutation test z-scores highlight the centromeres with an SCE enrichment (red) or depletion (blue). Before testing, the centromeres were resized to the T2T-CHM13 width to more accurately determine the expected SCE frequency within the centromeres. In addition, the centromere of chr1 was extended upstream to encompass a proximal inversion that masked centromeric breakpoints. For each non-acrocentric chromosome, the number of SCEs overlapping centromeres was counted after 1000 random permutations of SCE coordinates. The significance levels (*** < 0.0005, ** <0.005, * <0.05 p-value) and z-scores of enrichment or depletion were obtained using the permutation test (108). Human acrocentric and mouse chromosomes have poor p-arm mappability, preventing examination of SCE across the centromere. **(D) Genome-wide SCE density.** The distribution of SCEs are plotted at 1Mb resolution and normalized by the number of bins each SCE overlaps with, the total number of cells, and ploidy of each cell line. Bins with significant enrichment of SCEs are highlighted by an asterisk (BH adjusted p-value < 0.05). The genome-wide SCE enrichment p-values from 1Mb bins are provided in TabS3. Coordinates of SCE breakpoints are provided in TabS4. Centromeres are highlighted in gray. Raw breakpoint count distribution is shown in FigS7. The elevated rate of SCEs in HAP1^lig4-^ cells was accompanied by a higher variance in the genome-wide distribution (variance HAP1: 0.008; HAP1^lig4-^: 0.018), rendering the hot regions less likely to be statistically significant. **(E) Genome-wide distribution of scaled features and SCE correlation domainograms.** Domainograms show Spearman correlation between increasing number of 1 Mb bins at windows of 10, 25, 50, 75, 100, 125 Mb. BJ-5ta cell feature tracks are plotted at 1 Mb resolution, while Patski are plotted at 3 Mb.

Diploidization, however, is commonly observed with HAP1 cells. We devised a cell ploidy filter based on strand-neutral percentage and background estimate (FigS4). Subsequent analyses on HAP1 SCEs are for filtered haploid cells only. For the two diploid cell lines, we also found clear strand states and strand switches, and observed a median background of 3.3% and 4.4% for Patski and BJ-5ta cells respectively (FigS3A-C). Selecting cells with consistent profiles, low background and the expected distributions of strand states for each ploidy eliminated cells with low quality strand information and gave us a total of 1968 HAP1, 2176 BJ-5ta, and 931 Patski single cells (see Supplemental Text, Methods, FigS4A-C, FigS5A-C, TabS1).

Several methods have been developed for identifying strand switches (breakpoints) produced by SCEs and other SVs in Strand-seq datasets (34–36). However, the sequencing effort between Strand-seq and sci-L3-Strand-seq is substantially different. In Strand-seq, sequencing 700-1, 400k read pairs provides 3-5% coverage, while we typically sequence 30-200k read pairs given the substantially increased throughput with sci-L3 of typically 10-100k to potentially 1M single cells. For haploid cells the projected coverage is 1% at 100k read pairs per cell. For diploid cells, sequencing over 300k can achieve 2% coverage (FigS6A). For cost-effectiveness, we therefore performed sub-saturation sequencing that produced a 0.27-0.55% median coverage per cell (FigS6B). Despite this low depth, we were able to reliably identify breakpoints by developing a framework of filters targeting false positive calls (36) (FigS6C, see Methods). In brief, we filter strand switches as a result of misassembly, centromere artifacts, pre-existing SVs including inversions, translocations, and duplications (FigS7, “region filter”), and/or subclonal mutational CO outcomes including CNVs and inversions (TabS2, TabS4). Details for mapping such events and separating SCEs from these other types of breakpoints are discussed in respective sections. On top of manual validation, we examined cells across a range of less than 0.3% to over 1.8% sequencing coverage and found a steady median number of breakpoints throughout with no significant differences (FigS6D-E). We found a high correlation between SCE frequency and chromosome size (R^2^ of 0.66-0.94, FigS8A-B) that matched previous observations and further validated our calls (18, 24, 37). Overall, we were able to consistently identify true breakpoints and obtain high confidence sets of 21, 761 SCEs with median resolutions ranging from 128 to 225 Kb for both haploid and diploid cells (FigS6F).

The ability of sci-L3-Strand-seq to examine thousands of single cells not only enables the mapping of rare spontaneous mitotic COs, but also provides statistical power for the detection of subtle mutant phenotypes. As a proof-of-concept, we used the HAP1 cell line with a CRISPR-Cas9 engineered knockout (KO) of DNA ligase 4 (*LIG4*) (referred to as HAP1^lig4-^) and applied sci-L3-Strand-seq to measure the rate of SCE (38). With alternative end-joining and non-crossover HR still active, the expectation was to observe only a moderate increase in the number of crossovers. We analyzed 751 HAP1^lig4-^ single cells (TabS1) and found a significant increase in the number of SCEs per cell in the *LIG4* KO (p-value: 0.000376; Mann-Whitney U) with a median increase from 3 to 4 SCEs per cell (Fig2B). Interestingly, HAP1 cells have almost double the SCEs per haploid genome of diploid BJ-5ta and Patski cells, suggesting cell-type difference in the level of SCE. Overall, the ability to observe subtle alteration in the rate of SCEs demonstrates the scalability of sci-L3-Strand-seq and opens the doors for large scale mutational screens.

### Human centromeric regions have reduced levels of SCEs

Sci-L3-Strand-seq offers an advantage in detecting CO events by 1) providing strand information for every read, unlike SNVs, and 2) propagating this information across genomic distances. As a result, CO events can be detected even in poorly mappable regions, such as centromeres, by observing opposing strand directionality in flanking unique sequences. Crossover recombination during meiosis is suppressed within centromeres to prevent segregation errors (39). However, during mitotic proliferation centromeres are prone to ss- and ds-DNA breaks (40). These breaks could presumably be repaired by crossover, or other repeat-mediated annealing not involving crossovers. Meanwhile, studies of mosaic chromosomal alterations in healthy individuals and in various tumor sequencing revealed large-scale cnLOH as one of the most abundant type of SVs, frequently involving the entire chromosome arm, suggesting increased level of centromere-initiated inter-homolog crossovers (41, 42). We therefore examined whether mitotic centromeres on human non-acrocentric chromosomes have elevated or reduced levels of SCEs. After normalizing by centromere sizes from the T2T-CHM13 genome assembly (FigS9A) (43), we found that in HAP1 cells 5 out of 18 centromeres have significantly reduced rates of SCEs compared to the rest of the chromosome (p-value < 0.001; permutation test) (Fig2C). BJ-5ta cells also displayed significantly lower SCE levels for 8 centromeres (p-value < 0.01; permutation test). Interestingly, HAP1^lig4-^ cells had only 2 centromeres showing reduced level of SCEs (p-value <0.01; permutation test) with a general increase in centromeric SCE compared to HAP1. This includes a significant elevation of SCE level on chr17 and a shared elevation with HAP1 cells on chr19 (p-value < 0.01; permutation test). The exact position of the insertional translocation of chr15 into chr19 (44) was not mapped, but it could have disrupted the centromere, triggering instability and an enrichment of SCEs. Altogether, our data suggest centromeres generally suppress mitotic HR crossovers.

### Late replicating domains and several repeat classes show increased rate of SCEs

Focusing on the genome-wide distribution of SCEs we found HAP1, BJ-5ta, and Patski cells had a generally uniform distribution (Fig2D). Nonetheless, we found 4 out 6 hot regions (in red) to be significantly enriched for SCEs in HAP1, 1 in BJ-5ta, and 2 in Patski cells (Fig2D, TabS3 and TabS4). Although not all statistically significant, HAP1^lig4-^ cells have 11 hot regions of enrichment (Fig2B and D), suggesting HR has preferential sites of repair (45). We thus explored various genome correlates with SCE hotness (Fig2E, FigS10).

HR being the favored repair pathway within heterochromatin led us to examine the correlation of SCEs with replication timing domains and AB compartments (46, 47). Late replicating domains and B compartments are highly correlated with each other and composed primarily of heterochromatin with low levels of DNA methylation (48–51). As expected, we found SCEs were more associated with late replicating domains (Repli-seq) and B compartments (Hi-C) in human cells.

We also wondered if we could determine the type of break that initiated our SCEs based on correlation with END-seq and GLOE-seq datasets (52, 53). Interestingly, END-seq on etoposide (ETO) treated cells showed no correlation with SCEs in either mouse or human (FigS10 and TabS5). Instead, the no treatment controls were significantly correlated with SCEs, suggesting the observed SCEs are the result of spontaneous DSBs rather than Topoisomerase II (TOP2) induced DSBs (p < 6×10^-08^, Spearman correlation test). Moreover, we found no association of SCEs with single stranded nicks (SSN) from GLOE-seq, suggesting spontaneous SSNs are unlikely to be the lesions generating our SCEs (FigS10A and TabS5).

To test whether any other genomic or epigenomic features could be tied to SCE enrichment, we compiled a set of additional 22 human and 39 mouse features that included GC content, sequence divergence, repeat annotations, chromosome domains, and transcription factor binding sites (FigS10). In agreement with our permutation analysis, we found centromeric regions and satellite repeats were negatively correlated with SCEs across both human and mouse cell lines (Fig2C, FigS10A-B and TabS5). In contrast, we found several other repeat classes to be positively correlated with SCEs such as long interspersed nuclear elements (LINEs) and low complexity (LC) regions (FigS10A and TabS5).

Interestingly, the strongest positively correlated feature in mouse Patski cells were regions of sequence divergence between the C57BL/6J and *M.Spretus* genomes (p < 3.4×10^-07^; Spearman correlation test; FigS10B). The divergent sequences may prevent inter-homolog (IH) recombination (6), possibly due to the anti-recombination function of mismatch repair (54). Notably, chromosome fragile sites (CFS) were also positively correlated with SCEs in HAP1 and BJ-5ta cells (55). Overall, regions with higher rate of initiating lesions, structure forming repeats, and sequence divergence likely further influence the distribution of SCEs (56, 57).

### Sci-L3-Strand-seq enables accurate phasing of heterozygous variants and detection of LOH

In low-coverage single-cell sequencing, identifying copy neutral events, such as cnLOH and UPD, requires the separation or ‘phasing’ of heterozygous variants by their maternal or paternal origin. This is because for heterozygous sites, the occurrence of reads that represent both alleles are rare. Phasing is thus required to determine whether long contiguous stretches of variants are homozygous for one haplotype, or alternating between two haplotypes by randomly sampling at heterozygous sites. In regions where the orientation of the inherited DNA template strands differ, Strand-seq captures one parental homolog with Watson and the other with Crick strand-oriented reads (“WC coverage”). This one-to-one correspondence of strand state with haplotype can be used to obtain chromosome level phasing (22, 28).

As proof-of-concept that sci-L3-Strand-seq can accurately phase variants, we used Patski cells with known haplotypes (C57BL/6J x *M.spretus*) (32, 58) to assess phasing accuracy, and BJ-5ta cells, which have no phasing information available. Reads from WC state regions covered 85.7% (24.5M heterozygous SNVs out of 28.7M) and 92.3% (2.4M heterozygous SNVs out of 2.6M excluding chromosomes X and Y) of the total identified heterozygous sites within Patski and BJ-5ta cells respectively. Out of these, 91.6% (22.5M with WC coverage) were successfully phased for Patski and 90.7% (2.1M SNVs with WC coverage) for BJ-5ta cells (TabS6 and TabS7). Altogether, after applying corrections and filtering (summarized below, and in Methods), we obtained a final average phasing accuracy of 98.6% for Patski cells (Fig3A and FigS12). With human cell lines that are typically unphased, only successful phasing enables LOH calling; we therefore validated phasing by successful calling of LOH events.

**Figure 3.**
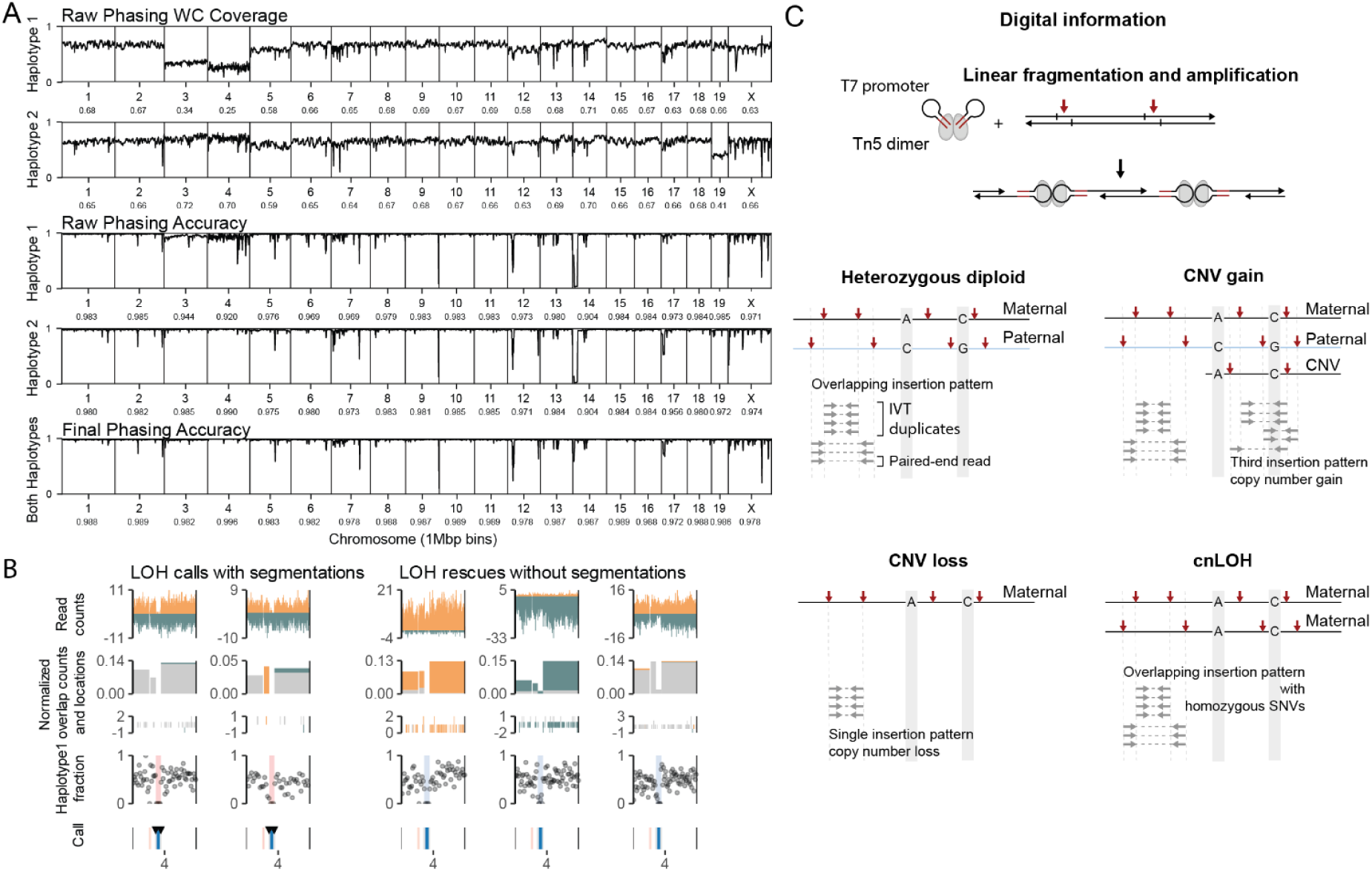
Phasing of SNVs and digital information for calling LOH and CNV. **(A) Phasing accuracy of Patski cells across the mm10 genome.** The phasing accuracy was plotted at a 1Mb bin resolution. In general only WC read (around half) will be informative for phasing. The SNV coverage from reads in the WC regions is shown in the top two panels, with the mean chromosome coverage shown under each chromosome label. Phasing by sci-L3-Strand-seq was compared to the existing references to evaluate accuracy. Raw phasing accuracies are slightly different between the two haplotypes (3rd and 4th panel). Bottom panel shows the final phasing accuracy after inference of the opposite haplotype, conflict removal, phasing with chr4 LOH, and inversion correction (FigS12, TabS6-7). Sharp decreases in accuracy can be explained by inversion (FigS13). **(B) LOH calls and rescues.** LOH calls (pink) on chr4 in two BJ-5ta cells are shown (left panels) with strand switch-based segmentation (two black triangles within call track). Three additional BJ-5ta cells (right panels) are shown without breakpoint calls for which the LOH calls were rescued (blue) and segmentation based on shared LOH with other cells. Read counts and overlapping reads are plotted as in Fig1A. Red vertical lines within the call track represent centromere bounds. Breakdown of LOH calling and rescue is provided in TabS8-10. **(C) Diagram of digital information within overlapping reads used for chromosome copy number counts.** Random genome fragmentation, barcode insertion and T7 promoter ligation (red arrows) is mediated by the Tn5 transposase dimer. IVT amplicon structure is depicted post ligation. The insertion leaves behind a 9 nt single-stranded overhang that is shared between adjacent fragments and ligated to the end of the Tn5 mosaic end duplex sequence (highlighted in red). Due to this shared 9nt sequence, to identify overlapping reads, an overlap is required to be greater than 12 bp (9 nt gap + 3 bp offset for deduplication). With IVT, each unique genomic fragment is linearly amplified that creates blocks of IVT duplicates (gray horizontal dashed lines with inward arrows). A disomic region (top left panel) will contain at most one overlap of two unique insertion patterns (ignoring IVT duplicates), each of which must originate from one of the parental homologs if heterozygous. In the event of a CNV gain, an overlap with three unique insertion patterns will be observed (top right panel). With a CNV loss, there will be no overlapping insertion patterns observed and only homozygous SNVs will be present (bottom left panel). The presence of overlapping insertion patterns and homozygous SNVs distinguishes cnLOH from CNV loss (bottom right panel).

Phasing of the WC covered sites with StrandPhaseR initially produced disjoint sets of either one or both SNVs at each site being phased; while the second haplotype can be easily inferred (FigS12A-C) (59). Unexpectedly, we also found an average 1.3% and 6% of heterozygous sites with the same variant assigned to both haplotypes in Patski and BJ-5ta cells respectively, which we labeled as conflicting sites and excluded from downstream analysis. For Patski, this improved phasing accuracy from 97.4% to 98.1%. With over 900 cells used for phasing, we could leverage the throughput to investigate how subclonal SVs affect phasing accuracy, and in some cases, phase with a different strategy. For example, when pre-existing, subclonal LOH events are present in over 80% of the cells (chr4), we found that manual phasing using the LOH provides proficient and accurate answers. Chromosome 4 has a mixture of both monosomy and UPD (Fig4A) and the UPD events could have WC coverage, potentially interfering with phasing. Another subclonal LOH on chr12 only affects about 30% of the cells and are exclusively deletions. Deletions do not contribute WC coverage and thus phasing without considering these SVs does not affect phasing performance. Lastly, we addressed inaccurate phasing due to subclonal inversions and centromere-proximal regions, both improving our overall phasing accuracy (see Methods).

**Figure 4.**
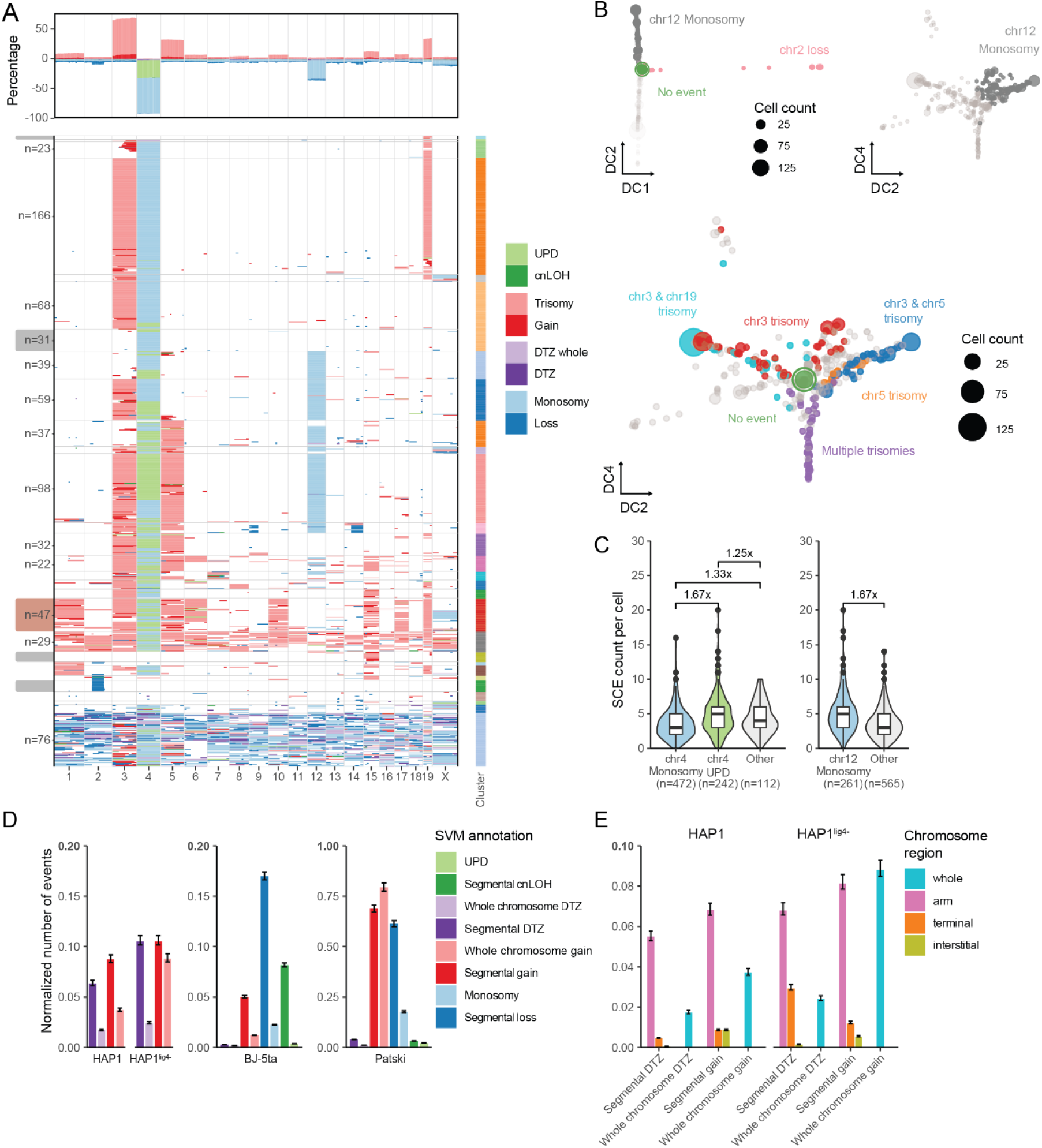
Annotation and quantification of mutational mitotic crossover outcomes. **(A) Heatmap of Patski SV annotation**. The heatmap shows only cells with annotated events. Cumulative percentages of cells with an annotation are shown in the top panel. Each row represents a single cell. Cells were ordered and grouped using hierarchical clustering. The chosen 29 clusters (not plotted with the 177 optimal number of clusters here) are shown on the right. Only cell counts (n) of clusters with more than 20 cells are shown. Clusters containing a single annotated event representing founder populations are indicated with a gray bar on the left. The cluster with multiple trisomies is highlighted in dark red. Heatmaps of mutational SVs in BJ-5ta, HAP1 and HAP1^lig4-^ cells are shown in FigS20A-C. **(B) Diffusion maps of mutational SVs in Patski cells**. Diffusion components 1 (DC1) and 2 (DC2) are shown in the top panel colored by segmental loss on chr2 (separated out by DC1, x-axis) and monosomy on chromosome 12 (separated out by DC2, y-axis). The bottom panel shows DC2 and DC4 colored by different combinations of trisomy. DC3 separated out two specific clusters, examined in FigS23B. Each point represents a unique set of annotated SVs, while the size of each point represents the number of cells that share the same set of annotated outcomes. High gain and high loss cells were excluded. Multiple trisomy represents cells with trisomy on chromosome 1, 3, 5, 6, 10, 15, 17 and/or 19 (dark red highlight in A). **(C) SCE rates within subsets of Patski SV outcomes.** In the right panel, cells were divided into groups based on the presence of chr4 monosomy or chr4 UPD or the absence of both (other category), while the left panel was divided based on the presence or absence of monosomy on chr12. The fold change of the median SCE counts per cell are shown for pairs of annotated events above the relevant plots. Significant differences for the left and right panels were calculated with the Mann-Whitney U test, comparing chr4 Monosomy vs chr4 UPD (p-value: 1.54×10^-11^); chr4 Monosomy vs Other (p-value: 2×10^-03^); chr4 UPD vs Other (p-value: 4.6×10^-02^); chr12 Monosomy vs Other (p-value: 1.94×10^-13^). Significant differences with additional summary statistics such as median, MAD, and fold change are provided in TabS13. **(D-E)** Pre-existing events (see FigS25A) and chromosomes without any event annotation are not plotted. Plotted events were counted per chromosome and normalized by the total number of cells and the ploidy of respective cell line. Sex chromosomes were excluded from BJ-5ta cells. Error bars show the 95% confidence interval. **(D) Normalized counts of mutational SVs.** Significant differences were calculated with chi-squared for HAP1^lig4-^ vs HAP1 with segmental DTZ (p-value 5.18×10^-04^; 1.7x), whole chromosome gain (p-value: 5.32×10^-12^; 2.5x). Contingency tables of additional event combinations with normalized counts, fold change and chi-squared p-values are provided in TabS14. **(E) Normalized counts of mutational SVs in HAP1 and HAP1^lig4-^ cells split by chromosome location.** Significant differences were calculated with chi-squared within HAP1 whole vs terminal DTZ (p-value: 1.20×10^-06^); HAP1^lig4-^ whole vs terminal DTZ (p-value: 1). Whole categories from (D) are included for reference. Contingency tables of additional event combinations with normalized counts, fold change and chi-squared p-values are provided in TabS14.

Without a ground truth, a reasonable validation of phasing in BJ-5ta cells is the ability to detect long contiguous regions of an extreme haplotype fraction that would otherwise be masked due to random haplotype assignment. We defined extreme as the proportion above 0.9 (primarily haplotype 1) or below 0.1 (primarily haplotype 2) of the total phased haplotypes for regions derived from breakpoint calls and chromosomal arms and considered these as potential regions of LOH (FigS14A and B). For Patski cells we identified a total of 2306 regions with extreme haplotype fractions (TabS8). The segmental deletion type of LOH has a 50% chance of containing identifiable strand switches (FigS18A). Nonetheless, the heritable and irreversible nature of LOH that leads to its clonal expansion enabled us to rescue the other 50% by querying all the cells for the same region to find LOH (Fig3B, see Methods). This led to the rescue of over 400 additional segmental LOH (0.508 versus the expected 0.5; TabS10). Taking the same approach with BJ-5ta cells, we identified a total of 228 regions with extreme haplotype fractions (TabS9), despite BJ-5ta cells having 1/12 as many phased SNVs compared to Patski. Overall, we identified multiple potential regions of LOH that reaffirmed our ability to accurately phase variants.

### Digital counting of overlapping reads better informs copy number

As with LOH, the low coverage and amplification noise of single-cell WGS methods makes it challenging to identify CNVs. Promisingly, a digital fragment counting approach implemented for LIANTI was shown to additionally reduce noise over using read depths alone (26). In essence, collapsing reads into insertion patterns (fragments) based on read ends and Tn5 overhangs gives an approximation that is closer to the absolute copy number. We took this concept further by first devising a rigorous deduplication scheme that accounts for possible IVT artifacts and additionally makes use of the sci-L3-Strand-seq strand information (FigS11, see Methods). Subsequently, we counted the number of overlaps between insertional patterns that are indicative of copy number gain or loss (Fig3C). These digital copy number counts are powerful when combined with additional layers of information including haplotypes and strand directionality. For example, a WC chromosome or region could have W-specific overlapping reads suggesting at least two W copies, but only one C copy, which immediately suggests a copy number gain event (2 W and 1 C). Similarly, haplotype-specific overlapping reads also suggest copy number gain or UPD. We thus build machine learning tools to integrate such information for SV calling.

### Strand switch- and clonal structure-based segmentation with machine learning classifiers facilitate accurate annotation of SVs

Chromosome segmentation for SV analysis in sparse single-cell sequencing is challenging. Conceptually, one can use a binned chromosome window (35), a binned number of SNV sites (10), or Hidden Markov Models (15) for segmentation. Mechanistically, mutational SVs that change copy number and/or strand directionality can couple with strand switches (20). Therefore, not all strand switches reflect an SCE event. For example, a deletion on a WC chromosome (50% in theory) retains only one of the two strands.

We thus use strand switches to segment the genome for calling other types of mutational SVs. As mentioned above, subclonal SVs that do not couple with strand switches (e.g., a deletion on a WW chromosome) can be rescued. We use strand switches-defined segments for initial LOH scan, and perform clonal rescues from the LOH scan, and then use both breakpoint-defined and rescued segments as candidate regions for SV annotation.

Using digital counts together with phased haplotype fractions, strand information, and various other metrics capturing read depth and background levels (FigS15A-C) we trained a set of support vector machine (SVM) classifiers for the annotation of cnLOH (segmental and UPD), CNV (gain or loss, segmental and aneuploidy), and “drop-to-zero” regions (DTZ; see Methods) (FigS16 and FigS17). The training consisted of manually annotated datasets with 1) 106 HAP1 and 62 HAP1^lig4-^ cells for haploid gain events, 2) 24 HAP1 and 13 HAP1^lig4-^ cells for haploid DTZ events, and 3) 75 BJ-5ta and 64 Patski cells for all four mutational event classes in diploid cells (see Supplemental Text and Methods). Using a 70:30 training to test data split we found all 12 SVM classifiers achieved high accuracies ranging from 0.971 to 1 (FigS16 and FigS17, TabS12). Examining the importance of each feature revealed the biggest drivers of accurate loss and cnLOH classification were the phased haplotype fraction and strand state features, while for gain it was the digital counts of overlapping reads (FigS16B, FigS17B and D). We found that stratifying the overlap counts with strand information facilitated the identification of unbalanced CNV gains and importantly helped us distinguish CNV loss from cnLOH. Incorporating haplotype information into the overlap counting also helped resolve additional CNV gains, losses, and complex events that we discuss below (haplotype-aware segmentation). Overall, our digital counting of overlaps substantially improved our ability to identify copy number changes with sub-saturation sequencing and together with other metrics enabled us to train accurate machine learning models for the annotation of various SVs.

### Haplotype-aware segmentation addresses lack of strand switch in cnLOH, enabling their identification, and improves other SV calls

Our rescue scheme, leveraging the high-throughput nature of sci-L3-Strand-seq, can map clonally expanded deletions lacking strand-based segmentation (expected for 50% of deletions), but also highlight the limitations of using strand switches for SV annotation (FigS18A and TabS10). We thus reviewed what type of SVs can be systematically missed with strand-switch-based and rescued segmentation. Most notably, cnLOH are generational and typically do not contain strand switches. They are copy-neutral in nature (unlike deletions) and thus lack loss of one strand that would enable initial discovery for a candidate region to scan for rescue (FigS18B). Manual examination of cnLOH events that were rescued revealed them to be complex events such as cnLOH embedded within a whole chromosome gain and therefore a deletion in nature. Motivated by the lack of strand switches with cnLOH, we devised a haplotype-aware approach for segmentation (FigS18A-C; see supplemental text).

Copy-neutral chromosomes without an LOH strand switch can contain 1) segmental cnLOH, 2) a linkage switch without a cnLOH produced by inter-homolog crossovers where the two recombined chromatids co-segregated, or 3) just normal heterozygous chromosomes without an inter-homolog recombination (FigS18C). In the first two scenarios, reads that contain phased variants can be split into Haplotype1 and Haplotype2 to reveal strand switches within the individual haplotypes. Firstly, given a WC region with a segmental cnLOH, the separation of the reads by haplotype forms C to WC and W to DTZ regions that creates identifiable point of segmentation (FigS18C top panel). For WW or CC regions with a cnLOH, the separation produces regions with a W or C to DTZ and a W to WW or a C to CC transition that are both hard to detect due to our low sequencing coverage (FigS18C middle panel). Nevertheless, if the segmentation was successful for clonal cnLOH in WC chromosomes, the cnLOH within same-strand chromosomes can be rescued. Secondly, inter-homolog crossovers that produce a linkage switch (uncoupled with BrdU labeling) without a cnLOH also produce identifiable W to C and C to W transitions with the separation of reads by haplotype, enabling segmentation (FigS18C bottom panel). While our analysis is focused on the detection of cnLOH, linkage switches without cnLOH are diagnostically important as they can have phenotypic consequences in cases where haplotype-specific enhancers are brought to and hijacked by oncogenes (60). It is also of interest to understand whether recombined chromatids in an inter-homolog crossover event have any bias in subsequent chromosome segregation (7). Notably, these events elude most bulk and single-cell WGS analysis. We manually examined newly annotated events and found the haplotype-aware segmentation enabled the identification of new cnLOH events, which contain the expected segmental DTZ in one haplotype and matching W and C state reads in the other haplotype for an otherwise fully WC strand-state chromosome (FigS19A and TabS11).

Using the haplotype-aware approach on Patski and BJ-5ta cells enabled not only the identification of cnLOH, but also de-novo segmental loss, and copy number gain events that originally lacked segmentation with strand-switch calling (FigS19). With low sequencing coverage the drop in read count for loss events can be difficult to identify (2 copies vs 1 copy). However, splitting reads in these regions by haplotype with the contrast in read counts between each independent haplotype makes differentiating the drop much easier (1 copy vs 0 copies). This is because regions termed ‘drop to zero’ (DTZ – 0 copies) containing only background reads can be thresholded on the background levels established from W or C-only regions (see Methods), essentially converting the problem into DTZ identification. This enabled us to annotate additional clonal loss events that were just outside of our haplotype fraction thresholds for rescue (FigS19B right panel).

For most of the new gain annotations that we manually validated, we found them to be trisomies with missed SCE calls (5 out of 7 manually examined events in Patski cells) (FigS19C first panel). In these scenarios, a missed strand-switch results in the two regions with opposing information to average out and create the appearance of a balanced copy number (e.g. 2-1 and 1-2 into 3-3). The haplotype-aware approach finds WC regions for a single haplotype, meaning at least two copies of one haplotype are present, signifying a copy number gain (FigS19C second panel). For non-WC regions, haplotype-specific overlapping reads indicate gains (FigS19C third panel). Notably, WC strand states present within both haplotypes can identify balanced gains (4-copy) and ultimately help disentangle complex events (alternate haplotype loss in a tetrasomy, FigS19C fourth panel) and high copy-number events (FigS19D). Beyond event annotation, the haplotype-aware segmentation approach also highlights inversions in the mouse reference genome that appear as reciprocal “SCE” breakpoints between the two haplotypes for WC chromosomes (FigS19E). These could also be actual homozygous inversions. Altogether, we show the haplotype-aware segmentation framework as described here facilitates the identification and interpretation of various mitotic crossover outcomes from low coverage single cells.

### Annotation of SVs reveals clonal population structures

After further stratifying our initial mutational SV annotations (SCE in the last cycle of BrdU labeling are not clonal) into whole chromosome or segmental events, we performed hierarchical clustering with the optimal number of clusters to get an overview of cells with at least one event (Fig4A and FigS20A-C, see Methods). In Patski cells we found 16 clonal clusters out of 177 total clusters with more than 10 cells that accounted for 58% (519/894) of all annotated cells (Fig4A). Since almost all the cells have a pre-existing chr4 LOH, 96.03% of Patski cells contain at least one annotated SV event compared to only 13.51% of BJ-5ta cells (FigS20E). Interestingly, we found a single annotated SV difference between most clonal populations in Patski, suggesting a stepwise addition of events where cells with only a single SV would represent the founder population. This led us to identify the following early founding events: monosomy on chr4, UPD on chr4, trisomy on chr15, trisomy on chr19, and a segmental loss on chr2 (Fig4A – highlighted on the side); which we later use to reconstruct a hierarchy of events that reveals a bias in SV occurrence. With the wealth of events in Patski cells we repeated the features correlation analysis and found the underlying (epi)genomic context did not allow us to distinguish between error-free (SCE) and mutational repair (the borders of segmental SVs; termed “adjacent breakpoints”) (FigS10B). In BJ-5ta cells, despite the sparsity of SVs, we observed 3 clonal populations with more than 10 cells (3/58) accounting for 28% (83/294) of all annotated cells (FigS21A). The most prominent clone contained an interstitial loss on chr4 (44 in 294 cells). Manual examination of coincident breakpoint hotspots (double breakpoints) without an initial SV annotation revealed a further 9% (4/44) of BJ-5ta cells with the same SV (Fig3B and FigS21). Our ability to identify most of these clonal events gave us confidence in our SV annotations.

Despite the smaller clonal populations in haploid cells, we found 14 out of 60 HAP1 clonal clusters and 7 out of 48 HAP1^lig4-^ with more than 5 cells that accounted for 41% (105/255) and 31% (48/157) of all annotated cells respectively (FigS20B and C). We found gains to be almost double that of DTZ events in HAP1 (78 gain to 46 DTZ) and almost three times in HAP1^lig4-^ cells (41 gain to 15 DTZ). The sci-L3 method does not require live cells in general, and could profile cells in the G1 cycle post BrdU labeling with fresh but lethal loss (like DTZ in haploid cells). The “clonal” DTZ present in a few haploid cells likely reflect independent loss occasionally for a whole chromosome. Overall, we observed substantially more cells with at least one annotated SV in HAP1^lig4-^ cells (20.91%) compared to HAP1 (7.98%), excluding the known disomy on chr15 (FigS20E). The substantial increase in mutational SVs, particularly whole chromosome gains, compared to a relatively minor increase in SCE rate in the absence of classical NHEJ is unexpected. One possible explanation could be that a fraction of the observed SCEs are a result of NHEJ repair of isochromatid breaks instead of mitotic crossover.

### Detection of Patski cells with a near-haploid genotype

Less restrictive clustering of Patski cells (29 total clusters) combined a set of 76 cells containing an almost ubiquitous level of segmental and chromosomal loss (Fig4A). Cells with similar characteristics were not found in any of the other examined cell lines. Certain cancers have been shown to aberrantly express meiotic genes that could lead to widespread chromosomal loss (61, 62). However, with only two haplotype switches per cell at most (FigS22A), meiosis-like division cannot explain the observed near-haploid genotype. Karyotype alterations in a single punctuated burst have also been observed in yeast (63) and human cancers (64). A study in colorectal cancer organoids showed multipolar spindle defects as the causal mechanism (65). These defects can give rise to cells with reciprocal karyotypes. Interestingly, a second cluster consisting of 29 Patski cells displays similar levels of segmental and chromosomal gain events (FigS22B), again suggesting a single punctuated burst event as the cause.

### Clonal lineage suggests monosomy as necessary intermediate for UPD

Having observed chr4 LOH in Patski cell with sci-L3-WGS (10), we now confidently resolve the events into monosomy and UPD with sci-L3-Strand-seq (Fig4A) with the implementation of digital copy number counts (applicable to sci-L3-WGS as well), and most conclusively by observing both W and C reads for a single haplotype. UPD can arise from two sequential nondisjunction events with a trisomy followed by the loss of the non-duplicate chromosome or with a monosomy followed by its duplication. An alternative process leading to UPD in a single cell division was observed in yeast, termed reciprocal UPD (RUD), where chromosome homologs instead of sister chromatids segregate into reciprocal daughter cells in a meiosis I manner (8). The presence of aneuploidy intermediates makes RUD an unlikely source of UPD in our diploid cells. Most of the clonal clusters consist of a mixture of cells with both chr4 monosomy and UPD that create mirror populations centered around the status of chr4 (Fig4A and FigS22B). Collapsing cells with identical events revealed chr4 monosomy had a higher diversity of events compared to chr4 UPD, suggesting monosomy preceded the appearance of UPD (FigS22C). We also observed two cells with a trisomy of chr4, in which the gained copy matches the lost haplotype in the monosomy, suggesting a reciprocal chromosomal nondisjunction produced the trisomy and the monosomy, and monosomy was the primary intermediate of UPD (the trisomy had the “wrong” duplicated haplotype). A cluster of cells that retain both parental copies of chr4 has a unique interstitial loss on chr2, and lack the highly prevalent chr3 and/or chr5 trisomy events, suggesting that they are more likely early founders in the population.

In BJ-5ta, both aneuploidy and UPD are particularly rare (Fig4D, FigS20A, TabS4). Previous analysis by single-cell WGS in BJ cells treated with UV radiation revealed almost no large-scale genome rearrangement (26), while UV of the same dose increases mitotic crossovers by 8, 000-fold in yeast (66). This suggests BJ cells could have an extremely stable genome that agrees with our observation.

Alternatively, cells with aneuploidy could have a strong selective disadvantage in BJ-5ta cells. As our single-cell assay does not require cells to be viable, we would expect high levels of single non-clonal events if selection were the main reason for the observed low levels of aneuploidy. Nevertheless, the higher fitness of UPD compared to monosomy, due to UPD being copy-neutral, makes negative selection less likely, particularly if the UPD is formed in a reciprocal, meiosis-like manner of chromosome segregation. Despite their low rate, we observed UPD only when a monosomy for the same chromosome was also present in the population, again suggesting UPD needs to go through a monosomy intermediate.

### Clonal lineage suggests recurrent SVs

Having identified several potential founder populations in Patski cells, we wanted to characterize the lineage structure of the various clonal populations using diffusion maps, a non-linear dimensionality reduction technique (FigS23A) (67, 68). We found four main branch points that can be broadly summarize by: 1) a segmental loss on chr2 (DC1), 2) chr4 LOH with combinations of chr12 monosomy, and chr3 and/or chr5 trisomy (DC2), 3) chr4 LOH with a combination of chr3 and/or chr19 trisomy (DC2), and 4) multiple trisomies encompassing up to 8 chromosomes (DC3 and 4) (Fig4B and FigS23B; see supplemental text). As mentioned above, the unique features of the chr2 loss cells, such as chr4 heterozygosity, suggest an early branching point from other prevailing clonal populations. The opposite haplotype of chr1 and chr15 trisomy in these cells, contrasting other clonal populations, together with a chr15 founder population, points to the same events arising multiple times independently (Fig4A, FigS22D). Similarly, the presence of both monosomy and UPD on chr4 with respect to other SVs, forming mirror populations, suggests several independent duplications of monosomy resulting in UPD occurred in the course of accumulating other SVs (FigS22B and D). The presence of chr4 LOH with only one of the additional trisomies on chr3, chr5, and chr19, or chr12 monosomy suggests these represent the earliest branching events of this lineage (Fig4B). Within these early branches the same set of events seem to have recurred, creating clonal populations with multiple combinations of events on both chr3 and chr12, or chr5 and chr12, or chr3 and chr19, or chr3, chr5 and chr12 (Fig4B and FigS22D). These, together with the multiple trisomy cells and the founder event clones reaffirm a striking predisposition of Patski cells for acquiring SVs within a preferred subset of chromosomes, a bias previously observed in yeast with perturbed checkpoint (69), and often seen in cancers (64, 70, 71).

### SVs alter the rate of SCEs

An increased rate of SCEs has been observed in numerous cancers and can be a sensitive indicator of genomic stress and instability (72–77). The significantly higher rate of SCEs in Patski cells compared to BJ-5ta (p=3.76×10^-04^; Mann-Whitney U) (Fig2B), along with the 96.03% of cells having at least one annotated event (FigS20E), raises the question of whether specific SVs coincide with increased rates of SCEs, or whether cells with unstable genome have higher instances of both. Focusing on dominant whole chromosomal events, we found cells with a chr4 UPD or a chr12 monosomy had a significantly higher number of SCEs (Fig4C). Since monosomy on chr12 is only present within cells that also have a chr4 monosomy or UPD, we broke down cells into five categories to disentangle the contribution of the different events to the rate of SCE (FigS24A, see Supplemental Text). We found chr4 monosomy decreases SCEs, while chr4 UPD and chr12 monosomy increase SCEs in an additive manner, suggesting the way the two events increase SCE rates are mechanistically independent (FigS24A, see supplemental text). Interestingly, neither the chr2 segmental loss nor trisomies on chr5, 15 and 19 had a significant impact on SCE rates, with only chr3 trisomy cells having a lower SCE rate compared to control cells (p=2.18×10^-04^; Mann-Whitney U; FigS24B and TabS13).

### Measuring the rates of SVs

As with SCE, the ability to measure the rate of various SVs arising in cells is invaluable to understanding genome instability in disease and in cellular perturbation experiments. The challenge is in disentangling the many branches of clonal, newly acquired, and recurring events to obtain the true rate of occurrence. Unlike with Patski cells, overcounting in haploid and BJ-5ta cells is less problematic due to the simpler clonal structure and smaller populations. As a compromise between preserving recurrent and removing propagated events we chose to remove the most frequent outlier events (FigS25A). Altogether, we excluded the known disomic region in haploid cells, five events across Patski cells, along with any near-haploid and high proportion of gain cells (FigS25A and B, see methods).

Comparing HAP1 and HAP1^lig4-^ cells, we found the biggest difference came from HAP1^lig4-^ cells having a significantly higher rate of whole chromosome gains compared to HAP1 cells (p-value: 5.32×10^-12^, 2.5x; chi-squared) (Fig4A, FigS25B and TabS14). A more frequent formation of unresolved crossover intermediates in HAP1^lig4-^ cells that leads to nondisjunction events could explain this higher rate. The reciprocal event of whole chromosome gain in haploid cells is whole chromosome DTZ. However, we did not observe a higher rate of whole chromosome DTZ in the HAP1^lig4-^ cells, likely due to selection or lack of viability. The counterpart in diploid cells would be a trisomy accompanied by a reciprocal monosomy. For BJ-5ta cells we observed a similar number of both event types, supporting a reciprocal origin (Fig4A, Fig1F and TabS14). In contrast, Patski cells contain a significantly higher number of whole-chromosome gains that suggest additional selection (p-value 4.95×10^-75^; chi-squared) (Fig4D, Fig4A, FigS25C and TabS14). The second most significant difference between HAP1 and HAP1^lig4-^ cells was in segmental DTZs, with HAP1^lig4-^ once again having the higher rate of occurrence (p=5.18×10^-4^, 1.7x; chi-squared) (Fig4A and FigS25B). Possible explanations for segmental DTZ could be break induced replication (BIR). Since sci-L3-Strand-seq only sequences the template strands, the conservative replication of BIR (78, 79), where both strands are nascent, would lead to complete degradation of the DNA and manifest as a DTZ event in haploid cells and a segmental loss in diploid cells. Micronuclei fractionated from bulk nuclei could contribute to both whole-chromosome or segmental DTZ (see Supplemental Text).

### Site of a HAP1^lig4-^ translocation shows an increased occurrence of SVs

After categorizing SVs into arm, terminal, interstitial or whole chromosome, we found that in both haploid cell lines, whole arm events were the most frequent (Fig4B, FigS25C-E). The relative proportions of the four categories were maintained between HAP1 and HAP1^lig4-^ cells, with the notable exception of terminal DTZ that does not span the whole arm (Fig4B, FigS25C-E). Genome-wide distribution revealed HAP1^lig4-^ cells have a substantial number of terminal DTZ events on the short arm of chrX (chrXp), which are absent from HAP1 cells (FigS20B and C). The region on ChrXp corresponds to a “SCE” hotspot that we later found to be translocated with the acrocentric chr13 (FigS7B and F, see **Strand-state correlation identifies known and novel translocations**). Interestingly, we found this region to be susceptible to three patterns of loss. In Class 1, only the translocated portion of chrX is lost, leaving chr13 intact (8 cells). In Class2, the entire chr13 is lost along with the translocated portion of chrX (2 cells). We manually inspected these two cells and found segments containing reads on the centromere proximal portion of the acrocentric chr13 that explain how the translocated chrX fragment was retained (FigS25F). In Class 3, a break on chr13 leads to the loss of the majority of chr13 to the right of the translocation, thus leaving the translocated chrXp intact (2 cells) (FigS20C and FigS25F). The third pattern involving almost the entire right arm of chr13 suggests the chrXp translocation is very close to the chr13 centromere, or in the sub-telomeric region of the acrocentric left arm (FigS25F). We also observed similar patterns with gain events that overall points to the chrXp-chr13 translocation forming a fragile region prone to further genome instability (FigS20C).

### Opposite strand reads identify reference or population inversions

Inversions are challenging to identify and, as a result, represent an understudied class of copy-neutral SV. The advent of Strand-seq is helping to address this gap (18, 19, 21), but even with Strand-seq, an inversion must be observed across multiple cells to be confidently identified, as it cannot be distinguished from a double SCE in a single cell. The large numbers of cells captured with sci-L3-Strand-seq makes identification easier, as the likelihood of observing subclonal inversions becomes greater. We identify inversions from the opposite strand of otherwise fully WW or CC chromosomes where: 1) a heterozygous inversion would show up as a region of WC, or 2) for homozygous inversions and haploid cells as WW in an otherwise CC chromosome, and vice versa (Fig5A and FigS26A, see supplemental text and methods). Overall, we identified 19 global (shared by HAP1 and BJ-5ta) inversions in human cells, along with 9 inversions unique to BJ-5ta cells, 2 unique to HAP1^lig4-^ and 5 shared by both HAP1 and HAP1^lig4-^ cells (Fig5B), with an overall median size of 300Kb (FigS26B). We found 65.7% of our inversion calls overlapped with previously described inversions (19), including 18 of our global inversions, 4 BJ-5ta unique inversions and 1 HAP1 and HAP1^lig4-^ shared inversions (Fig5B). For Patski cells, we identified a total of 38 inversions with a median size of 200Kb (Fig5B and FigS26B), with 1 overlapping with a limited study of 9 previously mapped inversions in C2 mouse embryonic stem cells. We also found the misoriented contig on chr14 in mm9 was still present in mm10 (FigS7D) (18).

**Figure 5.**
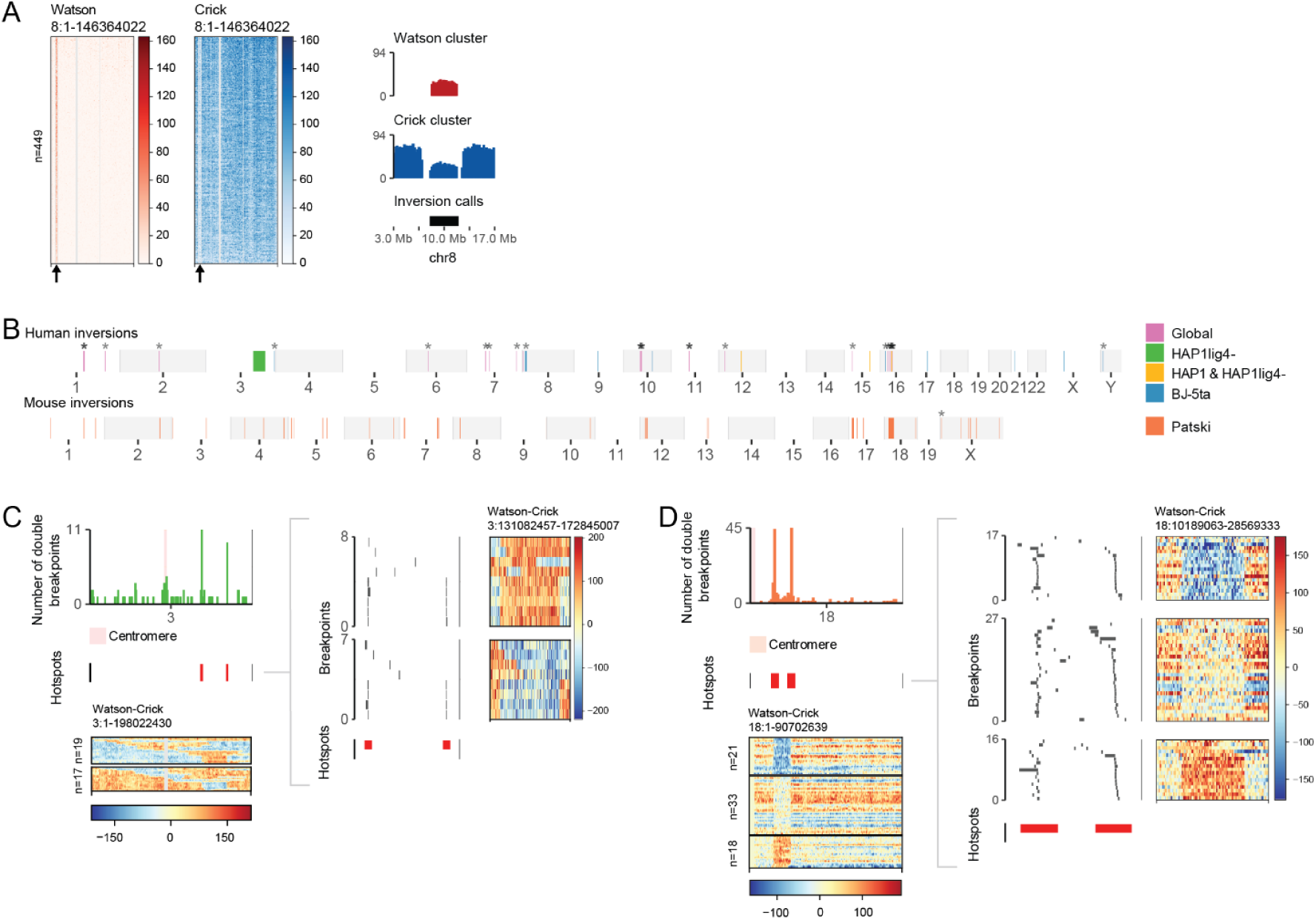
Detecting inversions with sci-L3-Strand-seq. **(A) A universal inversion on chr8 in BJ-5ta.** The left panel shows chr8 heatmaps from a subset of 449 single cells with both chromosomes in the C strand orientation. Each row represents a single cell plotted at an approximate resolution of 200Kb. W reads are plotted in red (representing background reads except for the inversion), and C reads are in blue. Masked regions are in gray. The same bin size and color code are used for heatmaps in Fig5 to Fig7 unless noted otherwise. Arrows highlight an inversion at the start of chr8. The right panel shows the pileup tracks of the heatmaps zoomed-in on the inversion, showing the median value for each bin across all plotted cells. The pileup for the W read heatmap of CC chromosomes is shown on the top track, and the C read heatmap of CC chromosomes is shown on the middle track. The bottom track shows our inversion call. **(B) Genome-wide locations of inversion calls.** Inversions identified across the three human cell lines are shown in the top panel. Patski inversions are shown on the bottom panel. The size of inversions have been scaled to a minimum of 1Mb to allow visualization. Inversions found in at least two cell lines were classed as global. There are no inversions private to HAP1 cells (all shared with HAP1^lig4-^). Even chromosomes are shaded gray for visual clarity. Calls overlapping with previously published inversions are highlighted with an asterisk (18, 19). **(C) Inversion detected from double strand switch hotspots in HAP1^lig4-^ cells.** The sub-clonal inversion measuring 27.98-33.8Mb was observed as a hotspot (red) of coincident strand switches shown in the top left bar plot. Matching whole chromosome heatmaps are shown underneath, where each row represents a single cell with at least two strand switches on chr3. The heatmaps show the inversion as a block of strand state change in a subpopulation of cells. Each roughly 200 kb bin shows the subtraction of W from C read counts. A zoom-in on the breakpoints within the hotspots is shown in the right panel, revealing 9 cells that contain paired strand switches in this region. The list of subclonal inversions with exact coordinates is provided in TabS2. **(D) Inversion detected from double strand switch hotspots in Patski cells.** The subclonal inversion measuring 4.9-14.5Mb was observed as a hotspot (red) of coincident breakpoints shown in the top left bar plot. Whole chromosome heatmaps are shown underneath, same as in (C). The zoom-in on breakpoints within the hotspots is shown in the right panel, revealing 55 cells that contain paired strand switches in this region (TabS2).

### Identifying large and sub-clonal inversions from breakpoint hotspots

Identifying inversions from the opposite strand of otherwise fully WW or CC chromosomes has three caveats: 1) opposite strand reads are either background or genuine inversions. Small inversions or inversions in a small percentage of cells regardless of their size may not meet our threshold to eliminate background reads, and could be missed; 2) using only WW or CC chromosomes is limiting; and 3) operationally we only use chromosomes without any breakpoints. We filter immediately adjacent strand switches (2 Mb in haploid and 10 Mb in diploid). Dual strand switches as a result of inversions smaller than the filter size are first removed, subjecting the chromosome to inversion analysis. However, larger inversions are omitted due to the presence of mapped SCEs. We therefore also called inversions by examining adjacent double SCE hotspots (Fig5C-D). SCEs are not clonal in nature and should not co-occur in the same cell even at hotspots. Out of the four candidate regions identified across all four cell lines, we found two bona fide inversions. The first inversion is located on chr3 in only 9 HAP1^lig4-^ cells (1.2% of the population) measuring approximately 30.9Mb (Fig5C). We did not find an equivalent inversion in the parental HAP1 cells, suggesting it was newly formed within the knockout cells. The second inversion located on chr18 was found in 39 Patski cells (4% of the population) measuring approximately 9.7Mb (Fig5D). Examining the strand states of the region shows the inversion is heterozygous. Our results show that sci-L3-Strand-seq can identify both global and sub-clonal inversions.

### Strand-state correlation identifies known and novel translocations

HAP1 cells have been previously shown to contain an insertional translocation of chr15 into chr19 and a balanced translocation between chr9 and chr22 (Philadelphia chromosome) with a BCR-ABL fusion (33, 44). Visualizing chr9 and chr22 in all HAP1 cells shows matching strand states of the fused segments with a detectable strand switch in the expected 50% of the cells (Fig6A). As with inversions, a translocation breakpoint needs to be matched across multiple cells to distinguish it from a SCEs. Based on this, we reasoned that from hundreds of single cells we could generate genome-wide strand-state correlation maps that would highlight the location of translocations. We calculated the Pearson correlation between 1Mb sized bins in 439 HAP1 cells that each contained more than 20 reads per Mb and plotted the resulting correlation matrix (Fig6B, see Methods). We found strong positive correlation between the expected chr15 disomic region and chr19 as well as between the terminal points of chr9 and chr22 (FigS27A and Fig6C), while the negative correlation highlighted some of the inversions (FigS27D-E). With 62 high-coverage HAP1^lig4-^ cells, we observed both known translocations alongside a new balanced translocation that arose between chr13 and chrXp (Fig6D and FigS27B). Additionally, we explored using cells with lower coverage at less than 20 reads per Mb and with a lowered resolution using 5Mb bins.

**Figure 6.**
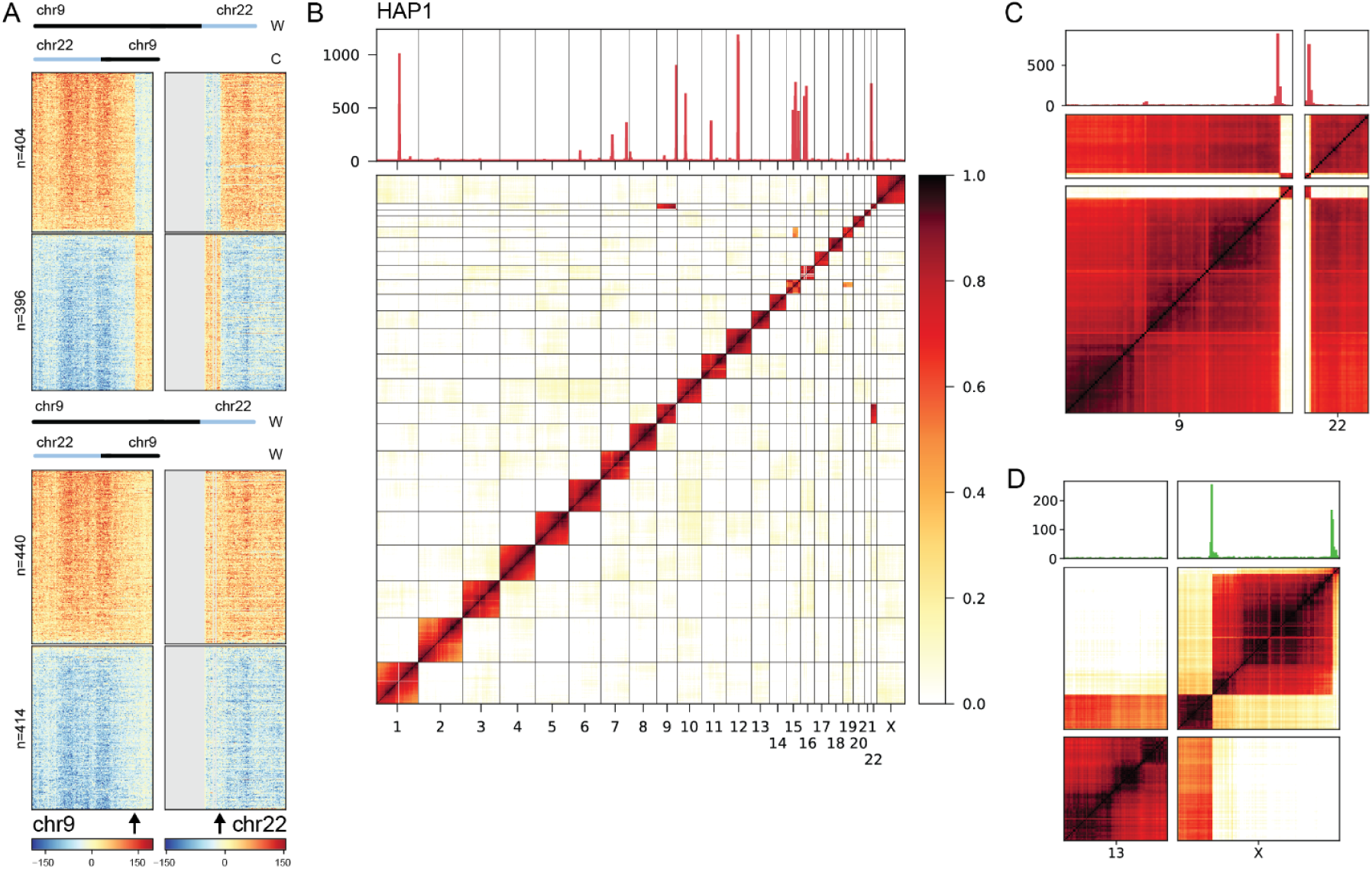
Positive Pearson correlation of strand states reveals translocations. **(A) Strand-state heatmap of a balanced translocation between chr9 and chr22 in the HAP1 cell line.** Each row represents a single cell. Each 200kb bin shows the subtraction of W read counts from C. Only the subset of cells without SCE on chr9 are depicted. The translocation appears as a strand switch in ∼50% of the cells (black arrows). Cells are ordered based on the strand state at the edges of each chromosome. **(B) Genome-wide heatmap of strand-state Pearson positive correlation in HAP1 cells**. Heatmap was plotted at a 1Mb bin resolution using 439 cells. Strand switches without the region filter are plotted in the top panel. Positive correlation *in trans* (off-diagonal) highlights mapped translocations. See also FigS27. **(C) A zoom-in of chr9 and chr22 in HAP1 cells.** The strand state of the terminal end of chr9 correlates with the strand state of the terminal start of chr22, highlighting a balanced translocation forming the Philadelphia chromosome. Top panel highlights the breakpoints (without the region filter) that pile-up at the translocation. Scale is the same as in (B). **(D) A zoom-in of chr13 and chrX in HAP1^lig4-^ cells.** The terminal start of chrX correlates with the strand state of chr13, highlighting a balanced translocation that was not present in the parental HAP1 cell line. The terminal end of chrX additionally shows a reduced correlation with the rest of the chromosome, highlighting an inverted duplication. Top panel shows strand switches without the region filter. Scale is the same as in (B).

From 689 cells we obtained three clear translocations with strong strand state correlations off the diagonal (FigS27C). Aside from translocations, the correlation heatmap also captured the inverted duplication event on the terminal end of chrX in HAP1^lig4-^ cells (Fig6D and FigS7F). We did not observe but expect chromosome-level positive correlation for telomere-telomere fusion of non-homologous chromosomes, and a decreased representation of WC chromosomes (50% expected) if homologs are fused. Neither BJ-5ta nor Patski cells contained any detectable translocations. For a subset of BJ-5ta cells the correlation heatmap likewise captured the inverted duplication at the start of chrX (FigS27D-F and FigS7E). Altogether, we show that sci-L3-Strand-seq enables us to detect both unbalanced and balanced translocations genome-wide.

### Resolving complex events provides new insights into the HAP1 disomic region

Having identified two inversions on chr15 in HAP1 cells, we unexpectedly found that one was located within the known disomic region and was never identified in previous studies (Fig7A) (29, 44). The duplication of chr15 was produced by an insertional translocation into chr19 (Fig6) and it must have originated from the lost parental chr15 prior to haploidization since it contains heterozygous SNVs (80). From this, we know the strand state of the disomic region is a composite of chr15 and chr19 (Fig7B).

**Figure 7.**
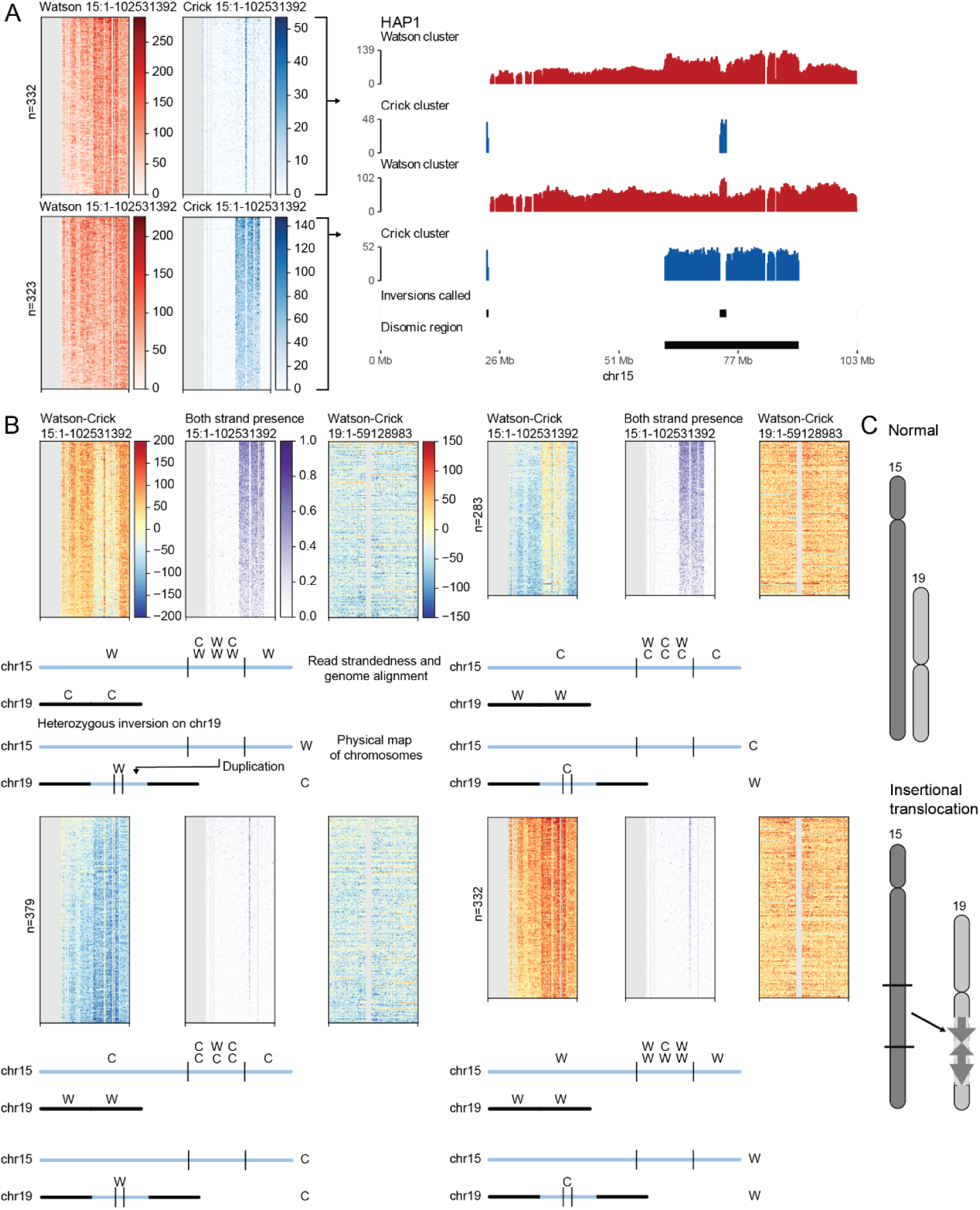
Resolving a complex rearrangement in haploid cells. **(A) A heterozygous inversion within the disomic region on chr15.** Each row represents a single cell. Pileup tracks of each heatmap show the median value for each 200kb bin, with an additional track showing inversion calls and the previously published location of the chr15 disomic region (right panel). **(B) The inversion is located within the newly gained copy created by an unbalanced insertional translocation.** Strand state heatmaps of chr15 and of the translocation chromosome. Each row represents the same single cell across all heatmaps. Cells were divided into four sets, each consisting of three heatmaps, based on the strand state of chr15 and chr19. In each set, the subtraction of W from C reads is shown on the first (chr15) and third (chr19) columns. The second column shows regions where both W and C reads are present. Below each set, we show a schematic plot of read strandedness and genome alignment on the top, and the corresponding physical map of chromosomes at the bottom. The other three possibilities of the inversion being within the original chr15 copy, or being homozygous on both copies, as well as the case without an inversion are depicted in FigS28. **(C) Diagram showing the final rearranged structure on chr19**. The original structure of chr15 (dark gray) and chr19 (light gray) is shown on the left, while the insertional translocation into chr19 containing an inversion is shown on the right.

Therefore, since chr19 shows a matching strand state with the gained copy, but opposite to the inversion, we know the inversion must be heterozygous and located on chr19 (Fig7B and C, FigS28). We can also confidently rule out the other two possibilities of 1) a heterozygous inversion on the chr15 copy, or 2) a homozygous inversion on both the chr15 and chr19 copies (see Supplemental Text). Resolving complex rearrangement involving duplication, translocation and inversion in the same region and providing the likely order of events highlights the power of sci-L3-Strand-seq for dissecting complex SVs.

## Discussion

Here we describe the extension of sci-L3 methods to Strand-seq that enables the detection of error-free SCE and mutational crossover outcomes in thousands of single cells (Fig1). The development of sci-L3-Strand-seq with complementary computational approaches offers a low-cost comprehensive assay for a multitude of applications including the study of HR repair, SVs and genome instability outcomes, along with high-throughput mutational screens.

With a 2-day experiment (FigS2), sci-L3-Strand-seq produces single cell whole genome libraries with strand information without any specialized equipment at a cost of $0.15 per cell for 10, 000 cells and appreciably less with greater cell numbers; on par with the previously published sci-L3-WGS (10). The scalability, and the resulting sub-saturation sequencing and low coverage of single cells motivated development of new computational approaches. Below we discuss the development of new genome segmentation methods, leveraging iteratively information at both the single-cell and population level, and new observations of genome instability enabled by these new developments.

Given the mechanistic nature of mitotic crossover, the resulting genome rearrangement is typically large, involving millions of bases. If we know the junction at a transition from heterozygous to LOH region, random allelic dropout at heterozygous sites due to low coverage is no longer an issue: hundreds of single-read covered SNVs to the left of the transition (heterozygous region) will collectively have a 0.5 haplotype fraction, and 0 or 1 for the LOH region on the right. The key to address sparsity in coverage is therefore proper chromosome segmentation. To our knowledge, we developed the first haplotype-specific strand switch-based approach for chromosome segmentation, which uniquely enabled detecting cnLOH and is conceptually novel from segmentation by HMM or fixed bin size (15, 20, 34, 35, 81, 82). NAHR between non-allelic repeats is typically more genetically tractable with various reporter assays (83–86) and genome-wide with population studies (87, 88), although for the latter, it is difficult to attribute the NAHR event to meiotic or mitotic origins. Allelic HR involving non-repetitive sequence homology from the sister chromatid or homolog has been difficult to map genome-wide. The original Strand-seq addresses the gap in mapping SCE, and here, haplotype-aware chromosome segmentation further allows us to dissect inter-homolog crossover with recombined chromatids co-segregating into the same daughter cell without causing cnLOH (FigS18C) or with recombined chromatids segregating apart into different daughter cells leading to cnLOH. We envision that the analysis framework pioneered here will eventually lead to knowledge on HR partner choice of: 1) homology on the sister chromatid, 2) allelic homology on the chromosome homolog, and 3) non-allelic repeats.

In addition to haplotype-aware chromosome segmentation, at the single cell level, we also provide SVM methods to integrate multifaceted genotype information, including strandedness, haplotype fraction using phasing inherent to the dataset, digital counting of copy numbers, and their combinations. The digital copy number counts were uniquely enabled by employing tagmentation-based genome fragmentation, overcoming unevenness in IVT-based linear or PCR-based exponential whole genome amplification in single-cell sequencing. These single-cell level analyses, together with haplotype-aware segmentation, were built into an iterative process to leverage population information from a large number of cells.

Here we summarize the five scenarios where our analyses go through single-cell level and population level cycles. First, we initially treat all strand switch calls as potential “SCE” (single-cell level), and examine single or double “SCE” hotspots for SV such as deletions, inversions and translocations (population level). This not only allows us to sensitively call subclonal SV sometimes present in only 1% of the cells, but also helps filtering potential SCE calls (single-cell) that are in fact boundaries of SVs.

Second, we start with all the cells for initial phasing of SNVs (population), and then call LOH events with strand-switch segments and imperfect phasing (single-cell). We also survey all the cells for regions with LOH calls and rescue the 50% of LOH expected to not have a strand-switch (population). Identifying the set of cells with LOH then collectively feedback to refine phasing (population level), orthogonal to strand-based phasing. Improved phasing accuracy allows better LOH calls and rescues (single-cell). Third, both LOH called with strand switch-defined segments (single-cell) and rescued LOH segments (population) were used as input of SVM for SV annotation. This strategy was devised to account for SV not associated with strand switches, but it also enabled better event calls in cells with particularly low coverage. Sci-L3 produces single cell libraries with 10-fold coverage range. Most ultra high throughput single cell sequencing methods have similar or higher variability of per cell coverage. Cells with higher coverage can thus serve as bait to inform segmentation. Fourth, similarly, cnLOH identified by haplotype-aware segmentation can rescue cnLOH without haplotype-specific strand switches. Fifth, we build clonal maps (population) from all annotated SVs (single-cell), and the clonal structure of specific events (population) help inform manual validation for training SVM (single-cell). In sum, we tackle the throughput vs. coverage trade-off by building iterative analysis processes that go through cycles of single-cell and population level information. We discuss eleven new bioinformatics tools in Supplemental Text developed and employed in this process, several of which are also applicable to sci-L3-WGS.

Combining improved analyses of all the seven types of mitotic crossover outcomes and the genome-wide and scalable nature of sci-L3-Strand-seq, we summarize new observations and insights gained from the examined mouse and human cell lines. We mapped an unprecedented number of 21, 761 SCEs from a total of 5, 826 single cells, which allowed us to examine subtle mutant phenotypes. Upon a knockout of the classical NHEJ pathway by deleting *LIG4*, we observed a moderate but significant increase in SCEs.

Despite its error-free nature, an increased rate of SCEs can be a sensitive indicator of genomic stress, instability, and possibly act as an early biomarker of tissue specific susceptibility to cancer (72–77). In addition to SCE, we also observed elevated levels of SVs in our *LIG4* knockout cells, including a somewhat surprising 2.5-fold increase in whole chromosome gain, suggesting the increase in HR repair may lead to an increase in gross chromosome rearrangement.

Sci-L3-Strand-seq enables examining crossovers emanating from a large-cluster of repeats like the centromere as long as there are enough unique sequences flanking the repeats. We examined the rates of SCEs within centromeres of human non-acrocentric chromosomes and observed a decreased rate of SCEs. The preference for HR in centromeres is central to their maintenance and rapid divergent evolution (89–91). Repeat amplifications and deletions in centromeres can be produced with single-strand annealing (SSA) and gene conversion between unequal repeats. With our observed decreased rate of SCE, it is possible crossovers are actively suppressed in favor of SSA and gene conversions (92). The size and composition of human centromeres can greatly vary between individuals or cell lines (93). Therefore, the CHM13 centromere lengths used may not be an ideal representation of our cell lines. Interestingly, in contrast to centromeres we found late replicating domains and several repeat classes, namely LINEs and LC regions, to be enriched for SCEs. These could be explained by either a higher rate of initiating lesions, such as non-B structure forming repeats, or a higher propensity of certain regions to utilize HR over other repair pathways (46, 47, 56, 57). Alternatively, it has been recently proposed that SCEs can arise independently of HR from stalled replication forks through POLQ-mediated end joining (94). The study utilized the haploid KBM-7 cell line (HAP1 parental line) and PARP inhibitor (Olaparib) treatment to induce replication fork stalling that produced under-replicated DNA in mitosis. In agreement with our data, they observed an increased rate of SCEs within CFS, suggesting our cells may sometimes be utilizing an HR independent repair pathway in difficult to replicate regions. We specifically chose to study spontaneous SCEs without damage-inducing agents. The correlation between SCE abundance with END-seq (profiles DSB) but not GLOE-seq (profiles SSN) suggests that DSB is the main source of spontaneous SCEs. Uniquely to inter-specific F1 hybrid mouse cells, we found a positive association with sequence divergence, suggesting that the sister chromatid is the favored repair template compared to diverged donor, likely due to the anti-recombination role of mismatch repair (54).

Aside from error-free SCEs, sci-L3-Strand-seq enabled us to examine different types and relative abundance of mutational SVs. With haplotype-aware segmentation that enabled cnLOH calls we found that segmental deletions are significantly more abundant compared to cnLOH (2x in BJ-5ta and 19x in Patski), contrary to analysis of mosaic chromosome alterations in the hematopoietic lineages (42) but in agreement with neuronal disorders (95). We also mapped a subclonal inversions involving as few as 9 cells (1% of the population), and resolved a complex rearrangement involving a duplication, a translocation and a previously unidentified heterozygous inversion present only in the translocated copy in the HAP1 cell line.

Mutational SVs allow us to build clonal lineages and dissect the sequential order of events. By examining subclonal structures, we found that: 1) monosomy seems to be a necessary intermediate for UPD in diploid cells; 2) DTZ in diploid arise in cells with pre-existing loss on the corresponding chromosomes; and 3) recurrent SVs from clonal map suggest certain chromosomes are hot for repeated duplication. In meiosis, crossovers engage homologs to provide necessary tension for chromosomes to be pulled into different poles in Meiosis I. Similar phenomena have been observed in the formation of reciprocal uniparental disomy (RUD) (8). However, we observed several lines of evidence that monosomy preceded UPD in the clonal histories of both Patski and BJ cell lines. Similarly, the observation that DTZ in cells with matching loss events in the population suggests DTZ in diploid cells requires a loss intermediate and thus is a two-step event. As for recurrent chromosome gain, McCulley and Petes (69) previously showed that yeast have significantly increased trisomy rate specifically for chromosomes II, VIII, X and XII in the checkpoint-defective diploid strains. Similar genome destabilization and/or selection mechanisms causing chromosome specificity could be shared in mammalian cells. Copy number changes are an almost universal feature of cancers, impacting fitness and cancer progression (96, 97). Patterns of whole chromosome and arm gains have been shown to be tumor specific and non-random, with recurrent events being linked to malignant transformation and growth (71, 98, 99). These recurrent events arising at different time points may also suggest that statistical modeling of tumor heterogeneity needs to assume polyclonal origins rather than using a strong prior belief that tumor is monoclonal (100).

The ability to simultaneously examine error-free SCE and mutational SVs could allow us to look at features differentiating these crossover outcomes, and their crosstalk. With the limited number of SVs observed in this study, we did not identify (epi)genomic features that clearly separate error-free SCE and mutational SVs. While they may share the same initiating lesions, repair partner choice of identical sister, homolog harboring heterozygosity, and non-allelic repeats remains to be explored in a larger scale study. We were able to link certain SVs with a variation in the level of SCEs, suggesting specific CNVs cause genomic instability. Similarly, in the example of chr13-chrX in HAP1^lig4-^, a translocation became a source of genome instability, resulting in a higher rate of SVs. In line with human cancer studies, these events may represent early aneuploidies that drive malignant transformation (99, 101), suggesting sci-L3-Strand-seq could be a powerful assay for detecting early cancerous states.

In addition to the Patski cells with gradual diversification, we found a surprising number of near-haploid cells. Several instances of haploidization have been previously described, including yeast cell haploidization in the absence of the homologous recombination protein RAD52 (102) and division without DNA replication observed in the skin cells of zebrafish (103). However, most notably, near-haploid karyotypes have been observed in leukemias and solid tumors (104, 105), with the HAP1 parental cell line (KBM-7) itself being derived from a chronic myeloid leukemia (CML) patient (106). We also observed cells with near-triploid karyotypes, suggesting they may represent reciprocal events of the near-haploid cells. Such punctuated bursts of karyotype alterations in a single cell division have been characterized in patient-derived tumor organoids of colorectal cancer (65). They showed such events can arise from multipolar spindle defects, where genome-duplication and mitotic failure can lead to tripolar division that produces three daughter cells with extensive karyotype alterations.

### Limitations

The limitations of sci-L3-Strand-seq include the low sequencing coverage that restricts the resolution of events (10). As a result, pinning down chromosomal elements as the source of events may be hard, especially for error-free SCE. Events such as microCNV as well as small and complex rearrangements in chromothripsis may not be fully detected (20). These resolution issues can be partially resolved by increasing the number of cells sequenced, if “mini-bulk” of cells with shared events can be identified by examining subclonal structures. While the resolution of SCE is difficult to refine for single events, piling up more SCEs genome-wide may eventually lead to better understanding of underlying genomic elements involved. Sci-L3-Strand-seq provides the technical throughput for exploring clonal lineages. However, it remains difficult to resolve the temporal order of all the events with certainty. This limitation implicitly affects accurately estimating the rate of mitotic crossover outcomes. For example, SCE rate is mapped for the single cell division for BrdU labeling, but the detection of inversions and translocations requires multiple single cells (nine cells in an example subclonal inversion we were able to map), preventing the identification of newly arisen events without minimal clonal expansion. Additionally, our assay uses nuclei without distinguishing their source of live vs. dead cells, although this has the advantage to look at raw DNA repair outcomes, staining for nuclei from live cells may be explored if needed for certain biological applications. Lastly, sci-L3-Strand-seq still requires BrdU labeling and thus is not applicable to fixed samples, rendering it less accessible for clinical samples. It is also less generalizable compared to sci-L3-WGS for model organisms without established BrdU labeling methods. Our development of haplotype-aware segmentation uniquely detects cnLOH. However, both cnLOH or co-segregated inter-homolog crossovers are rare in mammalian systems, except in early embryogenesis (107). Our methods have a greater potential in systems with smaller nuclei like yeast, for meiosis with homolog pairing, and organisms with somatic homolog pairing like dipteran insects including fruit fly and mosquitos.

### Conclusion

Sci-L3-Strand-seq substantially expands on the sci-L3 methods, enabling the detection of SCEs, inversions, balanced translocations, and chromosome level haplotype phasing useful for analyzing LOH and CNV. It additionally improves upon Strand-seq methods with Tn5-based digital counting and its linear amplification, high-throughput, and low-cost implementation. We develop computational methods that directly tackle the throughput vs. coverage trade-off, and iteratively leverage single-cell and population level information for comprehensive analyses.We demonstrated sci-L3-Strand-seq provides a quantitative view of crossover outcomes, with insights into clonal population dynamics. The generalizability of sci-L3 enables future adaptation of sci-L3-Strand-seq to other modalities such as RNA, enabling the examination of SCEs in populations of mixed cell types to address cell-type specific error-free and mutational HR (27). We anticipate sci-L3-Strand-seq will have a wide range of applications ranging from the study of basic molecular mechanisms to obtaining insights into complex diseases.

## Resource availability

### Lead contact and materials availability

Further information should be directed to and will be fulfilled by the Lead Contact, Yi Yin.

### Data and code availability

Raw sequencing data and processed data have been deposited at GEO as GSE292266 and are publicly available as of the date of publication. Additional raw data were obtained from GSE281238. All original code and supporting data has been deposited at Zenodo and are publicly available at 10.5281/zenodo.15036453 as of the date of publication.

## Acknowledgments

This work was supported by National Institute of General Medical Sciences (5R35GM142511 to Y.Y) and Damon Runyon Cancer Research Foundation (DFS-43-20 to Y.Y). T.R. was supported by the National Institute of Health Training Grant in Genomic Analysis and Interpretation (T32HG002536).

## Author Contributions

Conceptualization: P.C., T.R., Y.Y.; Data curation: P.C, T.R., Y.Y.; Software: P.C., T.R.; Formal analysis: P.C., T.R., Y.Y.; Funding acquisition: Y.Y., T.R.; Investigation: P.C., T.R., Y.Y.; Methodology: P.C., T.R., Y.Y.; Supervision: Y.Y.; Writing - original draft: P.C., T.R., Y.Y.; Writing - review & editing: P.C., T.R., Y.Y.

## Declaration of Interests

The authors declare no competing interests.

## Supplemental information

**Document S1.** Text and Figures S1–S28

**Table S1: Related to FigS3 and FigS4. Breakdown of cell counts across QC filtering steps.** Cells output from the sci-L3 pipeline are further filtered for coverage, background, and overall strand state percentage. In addition, several cells were removed by manual inspection of cells with more than 10 breakpoints. Each column shows the number of cells removed by each individual filter. The final cell counts show the cells passing all the QC filters.

**Table S2: Related to FigS7. Coordinates and annotation for hg19 and mm10 region filter.** The region filter primarily consists of strand switch hotspots that overlap with called inversion and translocations. The disomic region in HAP1 and HAP1lig4-cells was obtained from the manual merging of several hotspot regions. Similarly, the region on chr12 in Patski cells consists of two manually merged hotspots. The hg19 centromeric regions for chr1 and chr16 were blacklisted due to a large number of false breakpoints, in addition to the mm10 reference inversion on chr14. chrY in BJ-5ta cells, similar to chr16, also contained regions with high numbers of false strand switches corresponding to an inversion, an inversion proximal region, along with a region requiring blacklisting.

**Table S3: Related to Fig2D. Significance of SCE enrichment for 1Mb bins genome-wide.** SCE counts were normalized by the number of bins each breakpoint overlaps with, the total number of cells, and ploidy. SCE enrichment was determined by fitting a gamma distribution and calculating p-values using the cumulative density function, adjusted using the Benjamini-Hochberg approach (see Methods).

**Table S4: Related to Fig2D, Fig4 and FigS25. Coordinates of SCE breakpoints and SV annotations.** Each SV annotation (col1) is broken down by chromosomal arm, adjacent annotations (col2 - adjacent left; col3 - adjacent right) and classifies the event by location (col4; e.g. arm, terminal, interstitial or whole chromosome). In addition, regions derived from LOH rescue (col5), and cells with high number of gains, cells with high number of loss (near-haploid cells) or cells with spikes are marked (note). The strand states and deduplicated W and C counts are shown for each annotated region. Adjacent annotations include the chromosome left end (LE), chromosome right end (RE), and the centromeres (CEN) (FigS25A). Breakpoints are broken down into SCE and SV annotation adjacent.

**Table S5: Related to FigS10. Significance testing of each pairwise correlation between genomic features, SCEs and mutational SV breakpoints.** Results of a test for the correlation between paired samples. Pairwise Spearman correlation between SCEs or mutational SV breakpoints and genomic features was calculated and tested for significance (*** < 0.0005, ** < 0.005, * < 0.05). Bonferroni was used for multiple comparison correction.

**Table S6: Related to Fig3A. Patski cell phasing accuracy by chromosome.** Detailed breakdowns of phasing accuracy with respect to total heterozygous SNVs called, total heterozygous SNVs with WC coverage, and total phased sites for each of five phasing configurations and their combination: 1) raw phasing with StrandPhaseR, 2) inferred phasing where sites with only 1 phased base have the opposite base inferred for the missing haplotype, 3) phasing with conflicting sites removed (where the same base was assigned by StrandPhaseR for both haplotypes), and 4) phasing with chr4 clonal LOH, 5) inversion correction where the haplotype assignments are reversed inside inverted regions of chr14 and chrX.

**Table S7: BJ-5ta cell phasing statistics by chromosome.** Phasing statistics for BJ-5ta cells. Same as for TabS6, except missing accuracy statistics since BJ-5ta does not have a phasing ground truth. Total biallelic SNVs, SNVs with WC coverage, number of cells used for phasing, and phased sites for 3 different configurations: 1) raw phasing with StrandPhaseR, 2) inferred phasing, and 3) conflicting site removal.

**Table S8: Patski LOH calling statistics.** General statistics at each major step of Patski LOH calling, including raw LOH calls from breakpointR segmentations, chromosome arms, and rescues. Further statistics include the combined LOH calls for segmentations and whole arm fractions, where the LOH arm call is used when segmentation is unavailable on that chromosome arm, are also shown. LOH events without merging that were used as inputs to the SVM classifier are summarized in columns E & I, together with the statistics from LOH tables post-merging and post-manual updates.

**Table S9: BJ-5ta LOH calling statistics.** General statistics at each major step of BJ-5ta LOH calling, same as in TabS8.

**Table S10: Related to Fig3B and FigS13. Counts of LOH events segmented based on strand switches and arms versus rescues.** Based on our inability to segment LOH events in WW and CC regions (50%), the expectation is that we will not have the correct segmentation for 50% of segmental events. With the assumption of most events being clonal, we can use existing LOH calls to recover (“rescue”) the missing segmentation and enable LOH event annotation.

**Table S11: Related to FigS18 and FigS19. Manual examination of additionally annotated events resulting from haplotype-aware segmentation.** Examples from both Patski and BJ-5ta cells are shown with original segmentation (TabS4) (1), haplotype-aware segmentation with supplemental table merging (2), raw haplotype-aware segmentation (3), and manually resolved annotation that consists of a combination of raw haplotype-aware segmentation and manual merging into whole chromosome events (4).

**Table S12: Related to FigS16 and FigS20. Comparison of HAP1^lig4-^ annotated using SVM model from HAP1^lig4-^ or HAP1 cells.** HAP1^lig4-^ data was annotated independently with an SVM model trained using either HAP1 or HAP1^lig4-^ manual annotations. To match annotated events, at least a 20% overlap was required. All additional gain events annotated with the HAP1 SVM model correspond to the known chr15 disomic region. DTZ events not annotated with the HAP1 SVM model are dispersed across numerous chromosomes. Manual inspection of the DTZ events revealed they were false positive and borderline cases, leading to the use of the HAP1 SVM model for both cell lines.

**Table S13: Related to Fig4C and FigS24. Summary statistics of SCE counts per groups of mutational SV outcomes.** The number of SCEs for cells with whole chromosome clonal events (trisomy, UPD, monosomy) and a combination of thereof (chr3 and chr5 trisomy, chr4 and chr12 monosomy, chr4 UPD and chr12 monosomy) were examined to determine if there is a relationship between mutational crossover outcomes and the rate of SCEs. The “other” category contains all cells not present within the main category. The “chr19 no annotation” category is limited to cells with chr3 trisomy and chr4 monosomy, such that they match the cells with chr19 trisomy in all aspects except chr19 itself.

**Table S14: Related to Fig4 and FigS25. Categorical counts of annotated events and significant difference testing.** Significant differences between annotated events within and between cell lines were calculated using chi-squared. Pairwise comparison with contingency tables and fold change are included for between cell comparisons. Additional tables include breakdowns of events by their chromosomal location.

## Methods

### Cell culture

BJ-5ta (CRL-4001, ATCC) and Patski (gift from Disteche lab) cells were cultured in DMEM medium supplemented with 10% FBS and 1x Pen-Strep. HAP1 cells were cultured in IMDM medium supplemented 10% FBS and 1x Pen-Strep. All cells were cultured at 37°C, 5% CO2. For sci-L3-Strand-seq, cells were cultured with BrdU at 40 µM final concentration for 24 hours prior to fixation. HAP1 knockout cells for LIG4-(referred to as HAP1^lig4-^) are as described in (38) (gift from Shendure lab).

### Sci-L3-Strand-seq library generation

Fixation and nucleosome depletion was performed as previously described in ‘Methods and molecular design of sci-L3-WGS and sci-L3-target-seq’, subsections ‘Single Cell Preparation and Nucleosome Depletion’ and ‘Tagmentation (first-round barcodes) and ligation (second-round barcodes)’ (10). Notably, cells were trypsinized and fixed with 37% formaldehyde (final 1.5% concentration) in 1x PBS at a cell density of 1 million/mL for 10 minutes at room temperature with gentle tube inversion. For first round barcoding, nuclei were distributed into individual wells. Any remaining nuclei were stained with DAPI at a final concentration of 5 ng/µL and used for FACS as previously described (27). After the ligation reaction was stopped by the addition of the stop solution (lysis buffer (LB: 60 mM Tris-Ac pH 8.3, 2 mM EDTA pH 8.0, 15 mM DTT) with 0.1% Triton-X100 (LBT)), nuclei were pooled and stained with Hoechst-33258 to a final concentration of 5 ng/µL and the quenched population was sorted (100-300 nuclei per well) into a 96-well plate containing 3 µL of LB. Right after gap extension with the Bst WarmStart 2.0 polymerase at 68°C for 5 minutes and its inactivation with 80°C for 10 minutes, Hoechst-33258 was added at a final concentration of 10ng/µL and incubated at room temperature for 10 minutes in the dark. A 1 or 5 minute exposure in a Bio-Rad Gel Doc with a 365nm UV bulb, or a dose of 27-4000 mJ/cm2 was administered with a UVP crosslinker (CL-3000L) as previously described (27). At this stage, each well had a volume of around 7.7 µL, to which 0.3uL of USER enzyme (a mix of uracil DNA glycosylase and endonuclease VIII, referred to as UDG+EndoVIII in the text, NEB) was added and incubated at 37°C for 15 minutes. Next, 0.9 µL of its inhibitor UGI (NEB) was added with a further incubation at 37°C for 10 minutes. Finally, the T7 in vitro transcription system was assembled by adding 2 µL H2O, 2 µL T7 Pol mix and 10 µL rNMP mix (NEB, HiScribe T7 Quick High Yield RNA Synthesis Kit) and incubated at 37°C for 10-16 hours. The remaining section, ‘RNA purification, RT and SSS’, was performed exactly as previously described (10).

### Read processing and alignment

Sci-L3 read barcodes were identified and processed as previously described (10). Briefly, the main steps of de-multiplexing consist of:

1. Swapping read 1 (R1) and read 2 (R2) based on RT primer location such that all combinatorial barcodes are in R1. RT primers (GGGATGCAGCTCGCTCCTG) are identified based on Levenshtein distance of less than or equal to 3 from the reference sequence.
2. The 3rd round barcode (SSS, 6nt, no error allowed) and UMI (4nt) from R1 are attached to the read name. Reads are split into individual fastq files based on barcode sequence, enabling all subsequent steps to be performed in parallel. Reads without an identifiable barcode are discarded.
3. The remaining barcodes are extracted by first trimming the Tn5 Mosaic End double-stranded (MEDS) sequence from R1 (5’ adaptor: AGATGTGTATAAGAGACAG; maximum error rate: 0.2, minimum overlap: 19) and R2 (5’ adaptor: AGATGTGTATAAGAGACAG; 3’ adaptor: CTGTCTCTTATACACATCT; maximum error rate: 0.2, minimum overlap: 13) using cutadapt v3.0 (109) and parsing the MEDS adjacent sequences containing the 1st (Tn5, 8nt, 1 error allowed) and 2nd (ligation, 7nt, 1 error allowed) round barcodes (bc2 7nt - 4nt spacer - bc1 8nt - MEDS). Errors are calculated using Levenshtein distance. Successfully identified barcodes are written into the read name, else both R1 and R2 are discarded.

Subsequently, paired-end reads were aligned to a hs37d5 and mm10 hybrid genome reference using BWA v0.7.15-r1140 (110) and converted into BAM format with only primary alignments using Samtools v1.10 (111). Using the 1st and 2nd barcodes in the read names, the 3rd round barcode BAM files were additionally split into individual single cell BAMs. Barcode combinations with less than 10000 reads were discarded. Single cell hybrid genome BAMs were additionally split by mouse or human contigs, retaining only standard chromosomes. Each BAM was filtered for proper read pairs, a maximum insert size of 2000, and excessively soft-clipped reads based on a maximum soft-clipping ratio of 0.5 (defined as the number of soft-clipped bases over the number of matched bases). We found that excessively soft-clipped reads disproportionately contribute to higher background estimate levels (FigS3A). Finally, to avoid collisions, any split and filtered BAM that had less than 30% of total reads of the correct species or less than 10000 reads was discarded.

A general processing workflow implemented with snakemake (112) is available at https://github.com/recombinationlab/sciL3Pipe.

### Complexity estimation

Single cell library complexity was estimated using preseqR v4.0.0 (113, 114) by extrapolating the percentage of unique fragments being observed with an increasing sequencing effort. Per cell insert size was calculated as the mean of all cell fragment sizes. Percent coverage was calculated from the extrapolated duplication histogram by multiplying the unique fragment number by the mean insert size and normalizing by the mappable genome size. PreseqR was set to predict the expected number of species represented at least once in a given sample (r=1).

### Breakpoint calling and filtering

Strand breakpoints were called using breakpointR v1.10.0 (36). The window size for calculating ΔWs was set to 1Mb for haploid cells and 5Mb for diploid cells. The minimum number of reads required between two breaks for strand state assignment was set to 10 and 50 for haploid and diploid cells respectively. The smaller window size and lower minimum required reads were chosen for haploid cells due to the ability to filter false breakpoint calls based on strand states (strand state filter). Blacklisted regions for hg19 and mm10 (version 2) (115) were excluded from all analysis. For mm10, we additionally included a high signal region we consistently found on chr4-89540413:89544084.

breakpointR run command for haploid cells:

breakpointr(inputfolder = <>, outputfolder = <>, windowsize = 1000000, binMethod = “size”, pairedEndReads = TRUE, pair2frgm = FALSE, min.mapq = 20, filtAlt = TRUE, peakTh = 0.33, trim = 10, background = 0.05, minReads = 10, maskRegions= <>)

breakpointR run command for diploid cells:

breakpointr(inputfolder = <>, outputfolder = <>, windowsize = 5000000, binMethod = “size”, pairedEndReads = TRUE, pair2frgm = FALSE, min.mapq = 20, filtAlt = TRUE, peakTh = 0.33, trim = 10, background = 0.05, minReads = 50, maskRegions=<>)

### Haploid breakpoint filters

To filter incorrect haploid breakpoints we:

1. Breakpoints overlapping with our genomic region tables (see “Breakpoint region filter and SCE filter”) were removed (region filter; regions shown in FigS7). The region filter supersedes all other filters and clustered event rescues.
2. Remove breaks that are within (<=) 2Mb of each other (clustered breakpoint filter). The 2Mb cutoff was chosen based on the bimodal distribution of distances between nearest breakpoints (FigS6C). Breakpoints that have been removed by the region filter are not included when finding clustered events.
3. Only retain breaks with a strand state consistent with haploid cells (e.g. retain WW-CC and CC-WW, but remove WW-WC or WC-CC breaks) (strand state filter).
4. For pairs of breaks overlapping centromere coordinates or within 3Mb of the centromere, if the strand states of the outer segments for each break are identical, both breaks are removed (e.g. break 1 strand state: **WW**-CC, break 2 strand state: CC-**WW**) (centromere filter).
5. Rescue clustered events that have been filtered by examining the strand states of the outer segments of a cluster (e.g. **WW**-CC CC-WW, WW-**CC**), if the strand states differ, a new breakpoint is created by taking the outermost coordinates of the cluster. A rescued break supersedes the clusters event filter, centromere filter and the strand state filter.

### Diploid breakpoint filters

To filter diploid breakpoints we followed the same procedure as for haploid cells except the strand state filter was not used and for the clustered events filter and the centromere filter we used a distance of (<=) 10Mb (FigS6C).

### Breakpoint region filter and SCE filter

After an initial round of filtering (without a region filter) for both haploid and diploid cells, we identified regions with high density of breakpoints using the hotspotter function from the breakpointR v1.10.0 package (36). Hotspots for single and multi-breakpoints per chromosome were called separately and combined. For human cell lines, we additionally combined overlapping hotspots into single intervals and summarized the number of breakpoints and number of cell lines for each hotspot. Using these two numbers, we normalized the total number of breakpoints within a hotspot by the total number of cells and weighted each value by the number of cell lines across which a hotspot was found, giving us an “importance score” for each hotspot. An “importance score” above 0.1 was used to categorize hotspots into global and cell line specific regions. Ultimately, we found that regions with high breakpoint counts corresponded to either inversions (see “Inversion identification”), translocations (see “Translocation annotation”), duplications (FigS7E-F), or errors in the genome reference as previously described (18, 19). Manual inspection of hotspots without a structural variation (SV) or a reference error revealed these hotspots had no obvious direct strand switches despite breakpoints being called. We found these hotspots were adjacent to inversions, suggesting that inversions may be causing breakpointR to call false positive breakpoints, possibly due to a missed call directly at the site of the inversion. We therefore categorized these as “inversion proximal” in our region filter.

In addition to inversion proximal hotspots, we also found the human centromeric regions of chr1 and chr16 contained hotspots with a high frequency of false positive breakpoint calls. However, from chromosome-wide heatmaps of all cells we found that by applying a region filter, we were also removing a substantial number of true positive breakpoints. By clustering cells by the strand state on the end of each chromosome we could pick out cells with centromere breakpoints that were filtered out. To salvage these true positive breakpoints we implemented an extra centromere rescue procedure. For each cell with a breakpoint call within the centromeric regions of chr1 and chr16 (before region filtering), we examined the strand state of both chromosomal arms independently. If the composite strand state of the two arms indicated a strand switch (haploid: arm1 - WW, arm2 - CC | arm1 - CC, arm2 - WW; diploid: WW, CC | CC, WW | WW, WC | WC, WW | WC, WW | CC, WC | WC, CC) the centromere breakpoints were rescued.

The two sets of breakpoint calls (with and without the region filter) ultimately enabled us to classify breakpoints into categories corresponding to their origin (e.g. inversion, translocations, SCE, etc.). After each round of filtering, the strand state of each breakpoint and segment was re-evaluated with `background=0.1` and `minReads=10` settings.

Segmental structural variation can additionally result in a strand switch and be incorrectly classified as an SCE. To account for breakpoints resulting from SVs, we used the SV annotation from our SVM classifier (see “Annotation with Support Vector Machines classifiers) to define adjacent breakpoints.

### Haplotype-aware breakpoint calling

For haplotype-aware breakpoint calling, we first split aligned reads into separate BAM files based on the presence of haplotype 1 or haplotype 2 phased SNVs. Reads containing a mixture of both haplotype SNVs were excluded. For each cell, both haplotype BAMs were used as inputs into breakpointR with haploid settings described above. BreakpointR was not designed to detect drops to zero (see “Drop to zero annotation (DTZ)”) that are common features in cells split by haplotype. To overcome the limitation in calling breakpoints for DTZ regions, we devised two orthogonal approaches.

First, we reasoned that by artificially introducing background noise to each haplotype specific BAM, breakpoints could be called since the DTZ regions would contain sufficient reads to have a WC state assigned. To create a pool from which reads could be randomly sampled, we merged the BAMs of 50 cells with a background estimate of 0, i.e. every chromosome had both Watson and Crick reads. Based on the read depth of each cell we created dynamic bins (size dependent on coverage) across which reads would be randomly sampled. Random reads corresponding to 10% of each cell’s total reads were introduced as background noise. We found that with each random initialization, we recovered a disjoint set of breakpoint calls, leading us to perform five random initialization and taking the union of all breakpoints identified.

Second, we took advantage of the HAP1 SVM for identifying segments with DTZ (see “Drop to zero annotation (DTZ)”). For Patski cells we binned the entire genome into 4 Mb bins and calculated RCC and RCBC scores (see “Variables for Support vector machines (SVM) classifiers”) for each bin, which were passed to the SVM to enable identification of DTZ bins. The sparsity of SNVs in the human genome led us to use 10 Mb bins for BJ-5ta cells. To consolidate the bins identified as DTZ, we first removed any calls for bins with a mappable width of zero. Single bin gaps between consecutive DTZ bins were relabeled to DTZ and any single lone calls were removed. A final merging across two bin gaps was performed and the edges of each DTZ segment was used as a new breakpoint.

The final set of haplotype-aware breakpoints is a combination of the original haplotype-unaware breakpoints, breakpoints from haplotype split reads with five different background noise random initializations, and the edge bins of DTZ calls. For segmentation identified by more than one approach, we prioritized original haplotype-unaware calls, followed by calls from haplotype split reads, and finally DTZ calls.

### Sample selection and cell filters

To select high quality cells, we performed filtering using the background estimate, strand state percentage and coverage. The background estimate for each cell was calculated using breakpointR v1.10.0 (36).

Cells with a background estimate above (>) 0 and below (<=) 0.08 (FigS3B), and with more than (>) 2 median reads per megabase were retained. We grouped regions with a WC strand state and/or not biased enough to be assigned to WW or CC states together and defined these as having a “strand-neutral” state. Examining the strand-neutral percentage across diploid cells showed the expected centered distribution at 50%. We additionally observed a peak above 75% resulting from unsuccessful libraries (FigS4A). We therefore excluded any cells with a strand-neutral percentage above (=>) 75% from downstream analysis. Haploid cells displayed a tri-modal distribution of strand-neutral percentages, with the primary distribution representing good quality haploid libraries starting just below 25% (FigS4A). Since haploid cells should not have any regions with both W and C reads, we chose a stringent threshold that excluded any cell with a strand-neutral percentage of more than (=>) 15, leaving some margin to account for a known disomic region (44).

Cell barcode collisions in human and mouse cell mixing experiments were identified from the preferential alignment of read pairs to the species genome of origin. However, identifying collisions from alignments is not possible with experiments that mixed two cell lines of the same species. In such cases, we additionally utilized the presence of the Y chromosome to distinguish male and female cell lines, and examined SNVs private to each cell line to exclude misassigned cells (FigS3C).

To identify private SNVs, we first aggregated all cell line specific reads into a single BAM file (pseudo-bulk) and used BCFtools (111) to call SNVs. For BJ cells, an available bulk WGS dataset (SRA: SRP102259; (26)) was used instead of our pseudo-bulk. Records private to each cell line were output using the `isec` command. Multi-allelic SNV sites were split into biallelic records. For haploid cells, only homozygous SNVs were retained. For each single cell, a ratio of the number of reads containing a private SNV over the total number of reads covering private SNV positions was calculated. The ratios were scaled to fall between 0 and 1 and any cell above 0.5, meaning it contained more SNVs private to another cell line than its assigned line, was excluded from downstream analysis (FigS3C).

BCFtools SNV call run command:

`bcftools mpileup -O z --skip-indels --ignore-RG --redo-BAQ --min-BQ 13`

`--per-sample-mF -a ‘AD, ADF, ADR, DP, SP, SCR’ -f <genome.fa> <combined.bam> | bcftools call --multiallelic-caller --variants-only -O z -o <out.vcf.gz>`

We also manually examined cells with more than (>) 10 breakpoints and excluded any cells from downstream analysis that contained a high number of false positive breakpoints, leading to the removal of 2 BJ-5ta and 3 Patski cells.

### Chromosome heatmaps

To visualize the strand status across thousands of single cells we used an extension of the bioconductor package profileplyr v1.8.1 (116) and Deeptools v3.5.0 (117), implemented in sciStrandR, to plot heatmaps. Each row of the heatmap represents a single cell and each column a genomic interval. All heatmaps were plotted at a 200 Kb maximum resolution with counts per million (cpm) normalization, defined as: 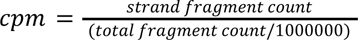 Bins overlapping blacklisted regions (115) and centromeres were masked. Count matrices were created for both Watson (W; positive) and Crick (C;negative) strands from R1 reads extended by 800bp. Additional matrices were produced from operations on the W and C count matrices. Heatmaps capturing the difference between strands counts were produced by subtracting the W from the C count matrix and setting empty bins to 0. Being able to plot the combined heatmaps with only a single scale makes it difficult to distinguish between WC states and low coverage regions as both will have values close to 0. To distinguish WC state regions, an additional matrix was constructed that highlighted the presence of both W and C reads. The matrix was constructed by dividing the parallel minimum by the parallel maximum of the W and C read counts for each bin.

A final matrix containing the number of breakpoints within each bin was created to help sort and categorize the heatmap matices: (1) Cells containing a single breakpoint within the plotting interval were split into the first two categories based on the strand orientation of the first 3% of bins. For cells with low coverage in the starting bins (sum of CPM less than 5), the first 10% of bins were examined instead. (2) Any cells with more than a single breakpoint were split into the next two categories. All cells with a breakpoint were additionally sorted by breakpoint position within each category.

### SCE distribution and correlation with genomic features

#### SCE enrichment and depletion within centromeres

The enrichment or depletion of SCEs from the centromeres of non-acrocentric chromosomes was evaluated using a permutation test from the regioneR v1.30.0 Bioconductor package (108). Before testing, the centromeres were resized to the widths within the CHM13 v2 reference (43, 118). Additionally, the centromere of the human chr1 was extended upstream to encompass a proximal inversion that masked centromeric breakpoints. For each non-acrocentric chromosome, the number of SCE overlaps with centromeres was counted after 1000 random permutations of SCE coordinates. An SCE was counted as an overlap only if it overlapped by more than (>) 40% of its size. Human acrocentric and mouse chromosomes were not evaluated due to poor mappability of the short p arms.

#### SCE hotspots

SCEs were counted across 1Mb bins and normalized by the total number of bins each SCE overlapped. Each bins counts were further normalized by the total number of cells and the ploidy of the cell line (HAP1 and HAP1^lig4-^: 1; BJ-5ta and Patski: 2). The normalized tracks were fitted with several common univariate probability distributions and based on Q-Q plots the gamma distribution was selected. P-values were calculated using the cumulative density function and further adjusted using the Benjamini-Hochberg approach as implemented in the core R stats package.

#### SCE correlation with genomic and epigenomic features

To examine the correlation of SCEs with various genomic and epigenomic features we downloaded processed datasets from previous studies, along with gene and various repeat class annotations. Repeat annotations were downloaded from the UCSC Genome Browser RepeatMasker (https://genome.ucsc.edu/cgi-bin/hgTables). The resulting collection of features were processed into either count data (gene and repeat annotations), signal tracks (genetic features, e.g. GC percentage) or normalized signal tracks (epigenetic marks). Obtained datasets varied in their window sizes and were further processes to obtain an averaged signal across 1Mb sized windows. SCEs were reduced to only the midpoint coordinate when counting overlap with 1Mb windows. Spearman correlations were computed on the resulting matrix and p-values were further adjusted using the Bonferroni method. For correlation domainograms, Spearman correlation was calculated between an increasing number of 1 Mb bins at windows of 10, 25, 50, 75, 100, 125 Mb. Feature tracks plotted alongside domainograms were additionally Z-score normalized.

For hg19, the following datasets were used: 1) HAP1 AB compartments, lifted over from hg38 to hg19 (119); 2) HAP1 and BJ Repli-seq replication domains (120); 3) HCT116 Gloe-seq (53); 4) NALM6 END-seq after etoposide treatment and no treatment control (NT) (52); 5) Chromosome fragile sites, lifted over from hg38 to hg19 (55); 6) HepG2 ChIP-seq for RAD21 and SMC3 (121).

For mm10, the feature matrix previously generated in (10) was subset and resized from 100Kb to 1Mb windows. The following datasets were used in the final matrix: 1) sequence divergence and 2) CNV gains between the strains (32); 3) END-seq after etoposide treatment and no treatment control and 4) CTCF and RAD21 ChIP-seq in C57BL/6 and Spretus activated B cells, along with 5) TOP2A and TOP2B ChIP-seq in MEFs (52); 6) Patski allelic ATAC-seq data and PolIIS5p ChIP-seq (122); 7) mESC Repli-seq replication domains (123).

### Phasing

#### Variant Calling

For Patski cells, a pseudo-bulk VCF was created from single cell BAM files using 1569 cells from the following libraries: nextseq190419 (yi293_GAACCG n=2, yi293_AGGACG n=55, total=57 cells), nextseq190830 (n=979), nextseq190906 (yi311_AATTGA n=38, yi311_ACTTAA n=8, yi311_CGGTGG n=5, yi311_GAGTTT n=4, yi311_GGGCCG n=29, yi311_GGGTGC n=24, yi311_GTCTAT n=12, yi311_TAAAGA n=6, yi311_TAAGCG n=29, yi311_TAATGC n=5, yi311_TATTCT n=8, yi311_TTTCGC n=21, total = 189 cells), and nextseq190907 (n=344). BAM files were merged, sorted, and indexed with Samtools (111). Human and mouse reads were split using a custom Python script. Variants were called using BCFtools `mpileup` and `call` commands against mm10 (111). The merged and filtered Patski VCF file was intersected with the published SPRET/EiJ SNVs (31) using the bcftools `isec` command and then filtered for biallelic SNVs to obtain the final set of SNVs for phasing. There were 33, 105, 073 total SNVs of which 28, 673, 432 had the heterozygous (0/1) genotype informative for phasing.

For BJ-5ta cells, SNVs were called with BCFtools `mpileup` and `call` against hg19 using previously published bulk WGS data (26), resulting in 4, 280, 029 total SNVs, with 4, 073, 787 SNVs for chromosomes 1-22 (sex chromosomes were excluded from phasing) of which 2, 569, 252 had the heterozygous (0/1) genotype informative for phasing. The 2, 847 multiallelic SNVs were not used for phasing.

#### Phasing with StrandPhaseR

Phasing was performed using StrandPhaseR (v1.0.0; R 4.1.0) (59). Phasing was performed using 949 Patski and 2180 BJ-5ta cells. WC regions were extracted using a modified version of the extract function in breakpointR (36), included in sciStrandR, and partitioned by chromosome for phasing (chr1-19 and X for Patski and chr1-22 for BJ-5ta). All cells that pass QC and are present in the WC regions file of a particular chromosome were used for phasing, except in two cases: 1) the phasing crashed prior to completion, in which case fewer cells were used (see “Errors Encountered During Phasing”), 2) the chromosome contained many LOH events, in which case a subset of heterozygous cells were used for phasing (see “Additional manual phasing for Patski”).

#### Inferred Phasing

StrandPhaseR generates phased haplotypes where substantial fractions of loci between haplotypes 1 and 2 are disjoint (818K of 1, 931K phased Patski sites and 107K of 174K phased BJ sites had both alleles assigned to haplotypes). SNV loci that were assigned for only one of the haplotypes were inferred for the other using the reference. In some cases, the same reference allele was assigned to both haplotypes, despite us only using heterozygous SNVs. In these instances, the ambiguously phased SNVs were filtered out as “conflicts”.

#### Correction of phasing over inversions

Previously published inversions were lifted over from mm9 to mm10 using the UCSC Liftover Tool (18, 124). The large inversions on chr14 and on chrX were also identified by our inversion calling. We therefore chose to use only these two inversions for correction, whereby alleles within any inverted regions are exchanged between the haplotypes 1 and 2.

#### Additional manual phasing for Patski

We found that Patski chr4 phasing accuracy was significantly lower than for other chromosomes. After an early pass of LOH calling where 761 out of 934 Patski cells were found to have Haplotype 1 LOH on chromosome 4, it became clear that most of the Patski cells were uninformative for phasing chr4. Initially, a subset of 10 heterozygous Patski cells were used for phasing of just 22K and 63K sites for haplotypes 1 and 2, respectively, but better SNV coverage was desired. As the majority of cells contain LOH, they are effectively pre-phased without the need for Watson-Crick strand information. Cells with the whole-chromosome haplotype 1 LOH fraction >= 0.9 were aggregated and sites deemed to be homozygous (allelic depth of the most frequent allele / allelic depth of all observed alleles >=0.75) were manually phased as the first haplotype. We required that at least 2 cells have a read at each SNV site and assigned the most frequent allele across all LOH cells as the Haplotype 1 allele. Because the Patski VCF was filtered for biallelic SNVs, the second unobserved allele at each position could be assigned to Haplotype 2. Unlike StrandPhaseR that requires Watson-Crick information to phase heterozygous sites, all SNVs in chr4 LOH cells with a coverage of 2 or more were phased. Similarly, we collected 281 Patski cells with hap1 LOH on chr12 (only 13 had hap2 LOH and were excluded) and attempted the same manual phasing. However, the accuracy failed to improve over StrandPhaseR and was therefore not used.

#### Phasing Accuracy Calculation in Patski

Phasing accuracy is calculated for both haplotypes using the Patski ground truth alleles with maternal SNVs from the REF/B6 and paternal SNVs from the ALT/SPRET. Because haplotype 1 and 2 assignments are made arbitrarily for each chromosome (in other words, each chromosome is phased into 1 of the 2 haplotypes and not the entire genome), we do not know ahead of time which haplotype corresponds to the REF or ALT ground truth. Accuracy was calculated by binarizing bases matching the REF allele from the Patski VCF to 0 and those matching the ALT allele as 1, then adding these values to their respective haplotype vectors. This results in one haplotype having a very large sum (almost exclusively 1s) and the other haplotype having a very low sum (almost exclusively 0s). Therefore, the haplotype with a large sum is determined to correspond with the ALT ground truth while the haplotype with a small sum corresponds to the REF ground truth. As these assignments are arbitrary and vary by chromosome, the max of each haplotype vector sum is taken (i.e., suppose 9 ALT alleles and 100 phased sites, then taking the maximum of 9 or 100-9=91 gives 91 and 50 ALT in 100 phased sites would reflect a random guess between the 2 haplotypes). Finally, the max is divided by the relevant number of heterozygous SNVs, het SNVs with WC coverage, or phased sites (using same toy example, 0.09 accuracy with respect to ALT or 0.91 accuracy with respect to REF alleles is reported as 0.91 phasing accuracy for that particular haplotype and 0.5 accuracy would represent the worst possible phasing performance). Invalid bases are counted as inaccuracies in the cases of raw and inferred phasing accuracy calculations; they are accounted for in the accuracy denominator after the conflict removal step. Plots were created using 1Mbp binned accuracies, while the phasing accuracies in supplemental tables and plotting labels are unbinned.

### Structural variation annotation

#### Inversion annotation

Using the heatmap count matrix categorization (see “Chromosome heatmaps”), we picked cells with no breakpoints and with whole chromosome WW or CC states. This chosen subset of each count matrix was collapsed into a bedgraph format by counting the percentage of cells at a particular position with a value greater than (>) 0 (percentage column operation). Subsequently, peaks above 0.4 (reads present in at least 40% of the cells with whole chromosome WW or CC states, roughly 20% in the population) were called as inversions with the exception of Patski, where due to a low number of cells with whole chromosome WW or CC states we increased the peak threshold to 0.5 (reads need to be present in at least 50% of the cells). Using the heatmap count matrices limits the inversion resolution to the 200 kb bins of the matrix, but it allows us to detect larger inversions (e.g. human chr8 in BJ-5ta cells), inverted duplications (e.g. chrX in BJ-5ta and HAP1^lig4-^ cells), and problematic regions (e.g. mouse chr14 in Patski cells). Inversion calling was run for WW and CC chromosomes with and without the region filter to ensure the filtering did not influence the calls.

Breakpoints resulting from inversions should appear as double breakpoints. Because of our clustered breakpoint filter, the double breakpoints from inversions less than 2 Mb in haploid and 10 Mb in diploid cells are removed, allowing these chromosomes to be used for inversion calling (only chromosomes with no breakpoints are used). However, larger inversions retaining their double breakpoints would not be used. Additionally, because of the chosen cutoff of 40% in human cells and 50% in mouse cells, double breakpoints originating from sub-clonal inversions would also not be identified. To overcome these two possible instances of missed inversion calls, we additionally examined breakpoint hotspots for chromosomes with more than one breakpoint. We identified a low frequency inversion in HAP1^lig4-^ cells on chr3 that we included in our final inversion call set. We also found a problematic region on chr12 in BJ-5ta cells that caused false double breakpoint calls. The same analysis also revealed double breakpoints on chr4 in BJ-5ta cells that matched a region of segmental loss that had a missed annotation in several cells. For downstream analysis we filtered both the chr12 and chr4 double breakpoints.

#### Translocation annotation

Translocated chromosomes, in the absence of a sister chromatid exchange, share the same template strand states. We used the correlation of strand states between chromosomes across N number of cells to identify potential translocations. A measure of template strand states was calculated for 1 Mb bins, using the W/C ratio as defined in breakpointR v1.10.0 (36):

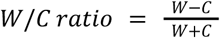

where W is the number of Watson fragments, and C is the number of Crick fragments within each bin. Fragments in blacklisted regions for hg19 and mm10 (version 2) (115) were excluded. A ratio was calculated only if a bin contained more than (>=) 10 fragments. Bins for which a ratio was not calculated in more than half of all cells were masked. To maximize the bins with sufficient read depth, we limited the analysis to cells with between (>) 20 and (<) 80 median reads per Mb. Any missing values were set to 0 and the resulting W/C ratio matrix was used to calculate a Pearson correlation of the strand states between all bins. For translocation, only the positive Pearson correlation was used, as the negative correlation highlights inversions. It is also possible to use cells with less than (<) 20 median reads per Mb to identify translocations, but at a reduced resolution (5Mb bins size tested).

#### LOH calling

Pileups were generated from single-cell bams with bcftools `mpileup` using the bulk SNVs described in the “Variant Calling” section. From the bulk VCF, homozygous sites were defined as 0.1 < REF allelic depth / (REF allelic depth + ALT allelic depth) < 0.9 while heterozygous sites were defined by a REF allelic depth >= 0.9 or <= 0.1. After phasing, sites with a total allelic depth > 0 are binarized as (1, 0) or (0, 1) for homozygous sites and (1, 1) for heterozygous sites.

LOH was scanned across chromosomes 1-19, X for 931 Patski cells (female) and 1-22 for 2178 BJ-5ta cells (male). In humans, chromosomes 13, 14, 15, 21 and 22 are acrocentric and only contain phased haplotypes on the q-arm. Strand state region files were generated - retaining regions of all sizes except for the mitochondria - using a modified version of the extract function in breakpointR (36), included in sciStrandR. The strand state regions, together with chromosome arms, were used for chromosome segmentation of each single cell. For chromosome arm segmentation, any segments spanning centromeres were split into 2 segments with the left segment terminating at locus centromere start and the right segment beginning at locus centromere end+1. Acrocentric chromosomes (all Patski and 5 in BJ-5ta) begin at centromere end+1.

For each single cell, SNVs from the joint phasing file were intersected with each cell’s pileup file, then mapped to each of the chromosome segments for calculating the haplotype fractions. Segment fractions are calculated as the sum of binarized tuples containing haplotype 1 (1, 0) or (1, 1) divided by the sum of binarized tuples containing either haplotype 1 or 2 (1, 0) (0, 1) (1, 1) within the defined segment; hence, each segment fraction is defined as the “haplotype 1 fraction”. Any segments that contain >= 20 phased SNVs and that span >= 10 Mb and exhibit a haplotype 1 fraction >= 0.9 or <= 0.1 are called as LOH candidates. Additionally, haplotype 1 fractions are calculated for each of the chromosome arms (a single arm per chromosome for Patski, two arms per chromosome for BJ-5ta). The LOH calling priority is: 1) chromosome segments that met the LOH candidate criteria above, 2) if a chromosome also meets the arm criteria, use the segmental LOH calls, 3) if a chromosome does not have segmental calls, but does have full arm LOH call, then use the arm call. The LOH calling itself is done in two stages, 1) the segments from strand state regions are examined via a “segment scan”, and 2) chromosomal arms are examined via an “arm scan”.

There were 3 cases where non-adjacent LOH segments were manually merged. In the first case, an LOH region was too short to be called but meets the fraction threshold (14 cells). In case 2, a short interstitial heterozygous region was between two LOH regions of the same type (both Hap1 or both Hap2). Here, both LOH regions are merged to include the heterozygous region, after which the haplotype 1 fraction is re-calculated (2 cells). Case 3 is similar to case 2 except that the interstitial heterozygous regions contain an annotated inversion and may be longer than 10Mb. Adjacent chromosome segments with extreme haplotype 1 fractions of the same type (Hap1 <= 0.1 or Hap2 >= 0.9) are merged into one segment in the LOH event table (63 cells). In all cases, the merge column of the LOH table is marked ‘M’ versus ‘Y’.

#### Haplotype 1 fractions and evidence thresholds

We tested three different haplotype 1 faction thresholds: 0.1 & 0.9 [0.9], 0.05 & 0.95 [0.95] and 0.85 & 0.15 [0.85]. With Patski cells, there were 6 fewer cells with LOH calls at the 0.95 haplotype threshold vs 0.9. These calls mostly appear to be artifacts resulting from noisy segmentation and aneuploidy (Fig3D). Additionally, there were 7 fewer Patski cells with LOH calls when the haplotype 1 fraction threshold was reduced again from 0.9 to 0.85. This time, however, many strong LOH candidates were not picked up, even in cells that contained many LOH events across chromosomes. For BJ-5ta cells, chromosomes 2, 10, and 11 exhibited the greatest range in raw LOH call counts, depending on the thresholds chosen. Some example cells demonstrate that many LOH calls are missed at the 0.95 threshold, while questionable calls are introduced at the 0.85 threshold. We also implemented an “evidence” threshold that required a minimum number of phased SNVs to be observed within a segment for it to be considered for LOH calling. We found many short “rescue” regions (see “LOH rescue”) in BJ-5ta cells that were called using evidence of 20 and 30, but were not seen when the evidence threshold was increased to 40 and 50. We began to see true positive LOH missed at the 50 evidence threshold. Based on these tests, we chose to use the 0.1 & 0.9 [0.9] thresholds, requiring all LOH segments to span at least 10 Mb and contain at least 20 phased SNVs within the candidate LOH region for calling, and at least 40 phased SNVs for BJ rescues.

#### LOH rescue

First, overlapping candidate LOH regions from the segment and arm scans are combined. These represent all the cells with a candidate LOH in a particular region. Subsequently, chromosomes without existing LOH calls are scanned, generating candidates of possible rescue segments. Rescues do not span a full arm, otherwise they would be detected by the arm scan. Each cell and chromosome pair can have numerous LOH rescue candidates, so the best rescue segments are first chosen by picking the longest non-overlapping segments for each cell. If LOH segments of the same class (both having fraction >= 0.9 or both <= 0.1) are overlapping, then the overlapping regions are automatically merged and the number of phased SNVs in the rescue segment - evidence - is recalculated, along with haplotype 1 fraction and segment size. As an example, if cell A has an LOH rescue candidates at chr1:10, 000, 000-50, 000, 000 (shared with cell B) and at chr1:40, 000, 000-60, 000, 000 (shared with cell C), then the LOH rescued for cell A is chr1:10, 000, 000-60, 000, 000. In rare cases where opposite haplotype 1 fractions overlap - less than 0.1 across chr2:10, 000, 000-chr2:40, 000, 000 (Hap1) and more than 0.9 across chr2:35, 000, 000-chr2:90, 000, 000 (Hap2) - the longer haplotype is fully retained with the shorter one being truncated so it is non-overlapping - less than 0.1 chr2:10, 000, 000-chr2:34, 999, 999 and more than 0.9 chr2:35, 000, 000-chr2:90, 000, 000.

### Data preparation for annotation

#### Deduplication

Fragments - read pairs merged into a single interval - were deduplicated using only read 1 (R1) coordinates after MEDS sequence trimming (FigS11). First, each R1 start position was extended upstream by up to 3 bp (offset), creating placeholder fragments that together with the original fragments were used to examine any start position overlaps. Fragments - original or their placeholder - with an identical start position to an initial fragment - the first fragment in the stack of overlapping fragments - were marked as duplicates. Finally, the R1 end positions after unmasking soft-clipped bases were examined and any fragments with identical end positions to the initial fragment were marked as duplicates. The union of both start and end position duplicates were considered IVT or PCR duplicates and removed from downstream analysis (FigS11).

We based the deduplication on only R1 due to the secondary enzymatic activity of T7 RNA polymerase called RNA-dependent RNA polymerase activity (RdRP), where T7 RNAP is capable of initiating transcription from RNA primed RNA templates resulting in RNA-RNA duplexes (125–127). In the event of self-priming or cross-priming of two transcripts, the 3’ end of the resulting transcript duplex would be truncated by RNases treatment resulting in a truncated R2 despite originating from an identical gDNA fragment (10, 26). We additionally observed soft-clipping of R1 start and end positions after trimming of MEDS. When soft-clipped bases were unmasked, we found these fragments had identical end coordinates. Consequently, to circumvent any potential issues with MEDS trimming, we incorporated the secondary deduplication step based on the end of R1 after unmasking soft-clipped bases.

#### Segmentation

Chromosomes of diploid cells were segmented for annotation in the following three ways:

A. Strand state region files containing all strand orientations (CC, WC, WW, Undefined) were parsed for each chromosome and single cell (36). The region files contain segments as defined by the filtered - without the region filter (see “Breakpoint calling and filtering” - breakpoint boundaries. Regions that completely overlap centromere annotations are split into two segments with the upstream segment ending at centromere start −1 and the downstream segment beginning at centromere end +1.
B. Each chromosome without breakpoints is additionally segmented by its constituent arms based on centromere coordinate. Acrocentric mouse chromosomes are segmented from the downstream centromere end+1 to chromosome end. In BJ-5ta, p-arms are segmented from 1 to centromere start-1 and q-arms from centromere end+1 to chromosome end.
C. Rescue segments (see “LOH rescue”) are defined by breakpoint boundaries from other cells that contain candidate LOH calls. Rescue segments are tested in every cell except for those already containing an LOH candidate on the same chromosome. Because a cell-chromosome pair can contain many rescue regions, the longest non-overlapping regions that exhibit haplotype fractions passing a threshold become the rescue segments. In the case that rescue segments overlap, they are merged. If merged segments overlap a centromere then they are split into two rescue segments with the upstream segment ending at centromere start-1 and the downstream segment beginning at centromere end+1.

Segments shorter than (<) 10 Mb or with less than 100 unique fragments were excluded from gain annotation. For drop to zero and loss annotation, only segments shorter than (<) 10 Mb were filtered.

For haploid cells, segmentation was performed as described for diploid cells in A, with the exception of still using the breakpoint region filter. Instead of keeping breaks within our marked regions as we did for diploid cells, we instead chose to manually add segmentation for the disomic region on chr15 (60471098-60471099 and 90518644-90518645, translocations on chr9 (132330486–134723723) and chr22 (22806017–24576823) and additionally on chrX (31719597–33113806) for HAP1^lig4-^ cells. The duplication on chrX in HAP1^lig4-^ cells was not added as it would have resulted in a 7Mb segment that would have been filtered out (10Mb segment filter). Manually adding breaks for known events, rather than retaining called breakpoints over these regions, yielded better annotation of the chr15 disomic region and the translocations in haploid cells without a noticeable impact on annotations across the rest of the genome (see “Post-processing and analysis of annotated events”).

#### Identification of high spike cells

We observed a subset of cells containing sharp peaks and extensive bin-to-bin variation in read counts. To avoid introducing noise into our SVM annotation classifier, we developed a “spiky score” to remove such cells from annotation. Fragment counts independent of strand were first counted across 1 Mb bins for each cell. A running median across 11 Mb was used to smooth the fragment counts and enabled sharp changes in fragment counts to stand out (FigS5A). The number of peaks (bins) with a smoothed absolute value above (>) 5 were counted for each cell. This threshold was chosen based on the general distribution of peaks across all cells. Because the number of spikes within each cell was dependent on sequencing depth, we additionally applied a loess normalization to the peak counts (FigS5A). Any cells with a normalized peak count - spike score - of more than (>=) 5 were labeled as “spiky” cells.

### Variables for Support vector machines (SVM) classifiers

#### Digital overlap counts (DOC)

The Tn5 transposase leaves a 9bp overhang after cutting. To distinguish adjacent from overlapping fragments, the pairwise overlap of deduplicated fragments was calculated and any pair with more than (>) a 12bp overlap was counted as an overlapping fragment region. In addition, the strand orientation of each overlapping fragment pair was categorized into Watson (DWOC), Crick (DCOC) - same strand overlaps - or opposite strand overlaps (DOOC). Finally, the digital overlap count for a particular genomic segment was normalized by mappability and sequencing depth (FigS17C):

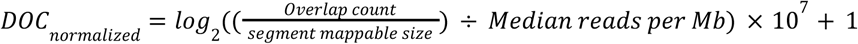

#### Relative chromosome coverage (RCC) score

The RCC score for a genomic segment is calculated as follows:

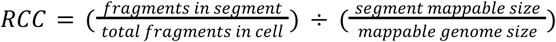

Genome mappability is defined as the proportion of a segment or genome that does not contain ambiguous bases (N) over the total number of bases. Due to differences in coverage between chromosomes, we additionally centered each segment’s RCC score by the median RCC score across all cells, such that a score of 2 could be generally interpreted as aneuploid (FigS17B):

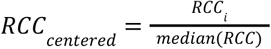

#### Background estimate and relative chromosome background coverage (RCBC) score

RCBC allows us to distinguish true signal from background noise and define regions with no signal as DTZ. Background estimates were calculated using breakpointR v1.10.0 (36). The background estimate is calculated from segments with a combination of WW or CC template strand states for which the opposite strand reads represent the background. The background estimate for a cell is the mean of the background from WW segments 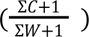 and CC segments 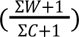, where C and W represent the total count of fragments with the respective strand state. From cells with high coverage that contained segmental and chromosomal deletions, we noticed that the background is strand independent and displays an even distribution across the genome (FigS17A). We used these observations to derive the RCBC score that captures the relative contribution of background to the true template strand signal:

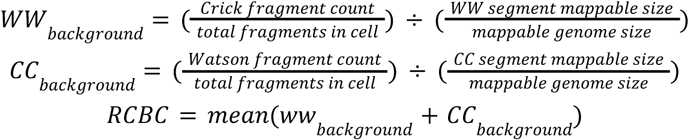

To account for differences in coverage between chromosomes, we additionally centered each RCBC segment score by the median RCC segment score:

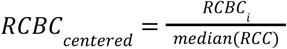

#### Final strand states

In haploid cells a key feature for segmental gains annotation is the WC strand state (FigS16). In instances where a breakpoint was missed, a false WC strand state can be assigned that negatively impacts annotation. A similar case can arise in diploid cells, where a segmental or whole chromosomal loss can result in a false strand state assignment. To circumvent such edge cases, we additionally examined strand states of each segment at a 5 Mb resolution, assigning states for each bin as defined in breakpointR v1.10.0 (36). Any segment with a W/C ratio of more than (>) 0.8 or less than (<) −0.8 (90% of fragments) would be assigned a WW and CC state respectively. A segment with a W/C ratio between (>) −0.2 and (<) 0.2 is considered a WC state. Everything else is marked as an unassigned state (set asby breakpointR). If the strand state of the entire segment does not match the dominant state from the 5 Mb bins (more than (>=) 50% of bins), the segment strand state is reassigned to a composite of the top two most frequent bin states - the “final strand state”.

#### Annotation with Support Vector Machines (SVM) classifiers

All SVM classifiers were run using the R package e1071 v1.7-9 (128) with the radial basis kernel and default settings. For classifier prototyping we used the Orange data mining toolbox v3.30 (129). For each cell line, a manual annotation dataset was curated. To evaluate the SVM classifiers we randomly sampled and split the manual annotation into a 70:30 training and test datasets. Confusion matrices were calculated using the R package caret v6.0-90 (130). To obtain a representative rate of (mis)classifications, we performed 70:30 random sampling of the manually annotated datasets 20 times and used the mean value of all the confusion matrices. We use F1 score to evaluate SVM performance. The importance of each feature was assessed by randomly permuting the values of each feature 20 times, one feature at a time, and measuring the resulting decrease in F1 score. For haploid cells, the same 70:30 test and training dataset split was used for assessing feature importance. In diploid cells the number of features used in each classifier was much higher (4 in haploid versus 9-11 in diploid), meaning the importance of certain features may not be apparent in particular random samplings of the manually annotated dataset. As a result, for diploid cells we used the entire manually annotated dataset to assess each feature’s importance. For the final annotation of all cells we trained each classifier on entire manually annotated datasets. Despite having a separate manual annotation for HAP1^lig4-^ cells for performance evaluation, the final annotation of both HAP1 and HAP1^lig4-^ cells used the HAP1 manual annotation.

#### Drop to zero annotation (DTZ)

Segments - especially in high coverage cells - contain background fragments that can prevent calling complete deletions, or “drops to zero”, based solely on empty bins (FigS15A). To identify DTZ segments, we trained an SVM classifier using a manually curated training dataset with RCC centered and RCBC centered scores as the input variables (FigS16B, FigS17B and D). The DTZ annotation setup is equivalent between haploid and diploid cells.

#### Loss of heterozygosity (LOH) annotation

Loss (monosomy) and cnLOH (copy neutral LOH) annotations only apply to diploid cells. The equivalent of loss in haploid cells is DTZ. In diploid cells we found the most important features for identifying LOH were haplotype fraction and final strand state. We therefore chose to annotate both loss and cnLOH with a single SVM (FigS17). All segments previously classified as DTZ were left out of LOH classification.

Additional features, such as digital overlap counts, were important for distinguishing between loss and cnLOH, and between LOH and gain. We ultimately used the following input variables to train an SVM classifier for LOH annotation: RCC centered, RCBC centered, mappable width, coverage percentage, final strand states, ROC normalized, RWOC normalized, RCOC normalized, haplotype 1 fraction (FigS17B and D).

When manually annotating cnLOH, we found a substantial bias of events toward a single haplotype (Hap1 or Hap2) (FigS14). The resulting underrepresentation of one of the haplotypes in the training datasets caused the classifier to completely overlook cnLOH events with haplotype 1 fraction either above 0.9 or below 0.1 when applied to the full dataset. To overcome this haplotype bias, we inverted the haplotype 1 fraction below 0.5 (|1-haplotype fraction|) so that both haplotypes were now represented by a single fraction going from 0.5 to 1 (instead of 0 to 1).

#### Gain annotation

To simplify classification, we combined any increase in copy number of a segment or a whole chromosome under a single umbrella term, gain. In haploid cells, we used the final strand states, RCC centered, ROC normalized and coverage percentage as the input variables for classifying gain. A small number of haploid gain annotations contained a composite final strand state - WW-CC or CC-WW (FigS16C and “Final strand states”). These states arose from missed breakpoint calls that resulted in a false positive gain call. From the composite strand state we know these classifications are false and therefore we corrected them during post processing.

For diploid gains annotation, we mostly used the same input variables as for LOH (FigS17B and D). We first removed all segments classified with LOH before performing gain classification. We additionally modified the same strand overlap scores. Normalized same strand overlaps (RWOC and RCOC normalized) are informative for identifying gains in WC state segments. In WC segments they have the potential to identify chromosome imbalance arising from only a single chromosome duplication (trisomy). However, with WW and CC segments we found instances where the normalized DWOC and DCOC score proved detrimental and the singular normalized DOC score was sufficient for an accurate classification. We therefore set the normalized DWOC and DCOC score to 0 for WW and CC state segments respectively.

Finally, we added the “strand-neutral” state percentage (see “Sample selection and cell filters”) as additional variables for gain annotation in diploid cells. The strand-neutral state percentage were important features for identifying BJ-5ta cells with a high percentage of gain events that likely correspond to 4n cells. Despite also including these two features in the Patski cell classifier, their importance for gain annotation was negligible (FigS17B). The exclusive presence of high percentage gain cells in BJ-5ta suggests a cell line specific feature. Removing the “high gain” cells before evaluating feature importance put the features important of the BJ-5ta classifier in line with the Patski classifier (FigS17D). The “high gain” BJ-5ta cells display a range of strand-neutral state percentages that are also present in cells without any gain classifications, showing that these two features are not amenable to a simple cutoff that could have been performed during initial sample selection (FigS20D).

#### Post-processing and analysis of annotated events

For HAP1 and HAP1^lig4-^ cells we ran the SVM annotation using segmentation with and without the region filter (see “Segmentation”). Even though we obtained better event calls with the region filter and manually added regions, in HAP1 cells there were four instances where the coordinates better reflected the annotated event without the region filter, and were therefore switched out. In HAP1^lig4-^ there were only two such instances that were also switched out. In addition, there were four false positive gain event calls in HAP1^lig4-^ cells that were not present without the region filter, and were removed from the final annotated set.

We can be confident that any segments in haploid cells with a WW-CC or CC-WW state are false for gain as these composite states arise from missed breakpoint calls in single copy segments. Therefore, we manually corrected any such misclassifications of gain. Similarly in diploid cells, any segments without an LOH annotation, but with a WW-CC or CC-WW state were reclassified as loss. The two exceptions to this rule were for the composite state segments on chr14 in Patski cells and chr1 in BJ-5ta cells. In both cases, the composite state was the result of an inversion. Therefore, any composite state segments on chr1 in BJ-5ta or chr14 in Patski and with a haplotype fraction between (>) 0.2 and (<) 0.8 were not reclassified as loss. Due to the relative low frequency of composite state segments in diploid cells we had to exclude CC-WW or both CC-WW and WW-CC states from the classifier due to their absence in the training data.

We next consolidated the segment annotation by merging adjacent segments with identical labels. To retain a contiguous segmentation for merging, segments shorter than (<) 10 Mb and segments longer than 10Mb but with less than 100 unique fragments were readded (see “Segmentation”) and labeled as short and low coverage segments respectively. Chromosomes with a single unique annotation label (ignoring short and low coverage segments) were considered whole annotations, with everything else falling under segmental annotation. For whole chromosome annotations, all individual segments were merged into a single interval. The haplotype 1 fraction difference between adjacent segments was additionally examined in diploid cells to avoid merging segments with a haplotype switch. Merging was allowed only if the difference did not exceed 0.2. If the difference was greater, all whole chromosome segments were relabeled as segmental and further examined if segmental merging was required. For segmental annotation merging, the following rules were applied:

● All consecutive segments with identical annotations are merged together.
● Annotated segments flanked by short segments (<10Mb) will be expanded to include a single short segment on either side. However, if a short segment is situated between conflicting annotations, no merging will be performed.
● Consecutive segments with identical annotations separated by a single short segment (<10Mb) will all be merged together with the short segment.
● For segments in diploid cells that have met the above criteria, the haplotype 1 fraction of each annotated segment is additionally examined to avoid merging segments with a haplotype switch. If the haplotype 1 fraction difference of the two segments does not exceed 0.2, the segments are merged.

Following merging, annotated events were examined for a set of location features such as the right side of the centromere (RCEN) and the right end of the chromosome (RE) that enabled us to further classify events into arm, interstitial, terminal and whole (FigS25D). In the example of RCEN and RE, the annotation would be labeled as an arm event.

#### Clustering and diffusion maps of annotated events

To construct the distance matrix, each cell was first partitioned into 1Mb bins and each bin that overlapped an annotated event was assigned a value 1 to 8, depending on the event category. To improve the clustering resolution we additionally added a weight for each set of annotation categories. For haploid cells, DTZ and whole chromosome DTZ was given a weight of 0, while gain and disomy was given a weight of 40. For diploid cells, DTZ and whole chromosome DTZ was given a weight of 0, loss and monosomy was given a weight of 20, cnLOH and UPD was given a score of 40, and gain and trisomy was given a weight of 60. The weights were chosen based on the copy number of each event category.

Alternatively, to prioritize cluster separation by chromosome 4 monosomy vs UPD - allowing us to highlight the mirrored nature of clonal populations whose sole branch point was the status of chromosome 4 - we altered the weights for DTZ and whole chromosome DTZ to 40, and cnLOH and UPD to 0. The weighted count matrix for each cell line was subsequently used to obtain the Canberra distance matrix using the factoextra package in R (131). We chose Canberra distance as it was the only distance metric in combination with ward.D2 hierarchical clustering that separated “high gain” from near-haploid cells (FigS22B). An optimal number of clusters was chosen using the silhouette method implemented in factoextra (131). With the maximum number of clusters to consider set to 200, the optimal number of clusters for HAP1 cells was 60, while for HAP1^lig4-^ cells it was 48, and for BJ-5ta cells it was 58. For Patski cells the optimal number of clusters was 177. However, with 177 clusters, most clusters contained only a single cell. This led us to split Patski cells into only 29 clusters that still preserved the pattern of clonal divergence. Lastly, we chose to manually reorder the 29 clusters to better highlight the gradual accumulation of events (Fig3D).

Diffusion maps were made using the destiny R package v3.8.1 (68). Assigning the highest weights to gains and trisomy raised concerns that the diffusion map clusters would be driven by the weights, rather than categories of events. We tested numerous distance metrics and chose to use Canberra, the same distance used for hierarchical clustering, due to its tolerance of different weight combinations (FigS23A). With Canberra, we observed minimal impact on the structure of the diffusion maps when the weights were inverted to 60 for DTZ and whole chromosome DTZ and 0 for gain and trisomy. Other weights, such as the default euclidean distance used by destiny, turned out to be substantially influenced by different weights (FigS23A).

For Patski and BJ-5ta cells, the percentage of the mappable genome covered by gain or trisomy was calculated for each cell within a cluster and the median of all cells was used to determine which clusters contained cells with unusually high percentage of gain - “high gain” - annotations. Similarly, the percentage of loss or monosomy was used to determine “high loss” or near-haploid clusters. A threshold of greater than (>) 45 percent median coverage was chosen to identify “high gain” and “high loss” clusters. Despite the speckled nature of gain annotations in Patski cells classed as “high gain” - compared to the contiguous gains in BJ-5ta cells - we chose to exclude “high gain” cells from downstream analysis for both cell lines.

#### Comparing annotated event rates within and between cell lines

We used chi-squared tests to determine significant enrichment of annotated events (SVs) within and between cell lines. In addition, we applied chi-squared tests to ascertain the significance of chromosomal location for each event class. For BJ-5ta cells, the sex chromosomes were excluded. For contingency table generation, events were counted on a per chromosome arm basis, accounting for cell ploidy. For whole chromosome events, counting by chromosome arm produces a count of two - except for acrocentric chromosomes - and counting segmental events by chromosome arm produces a count of one.

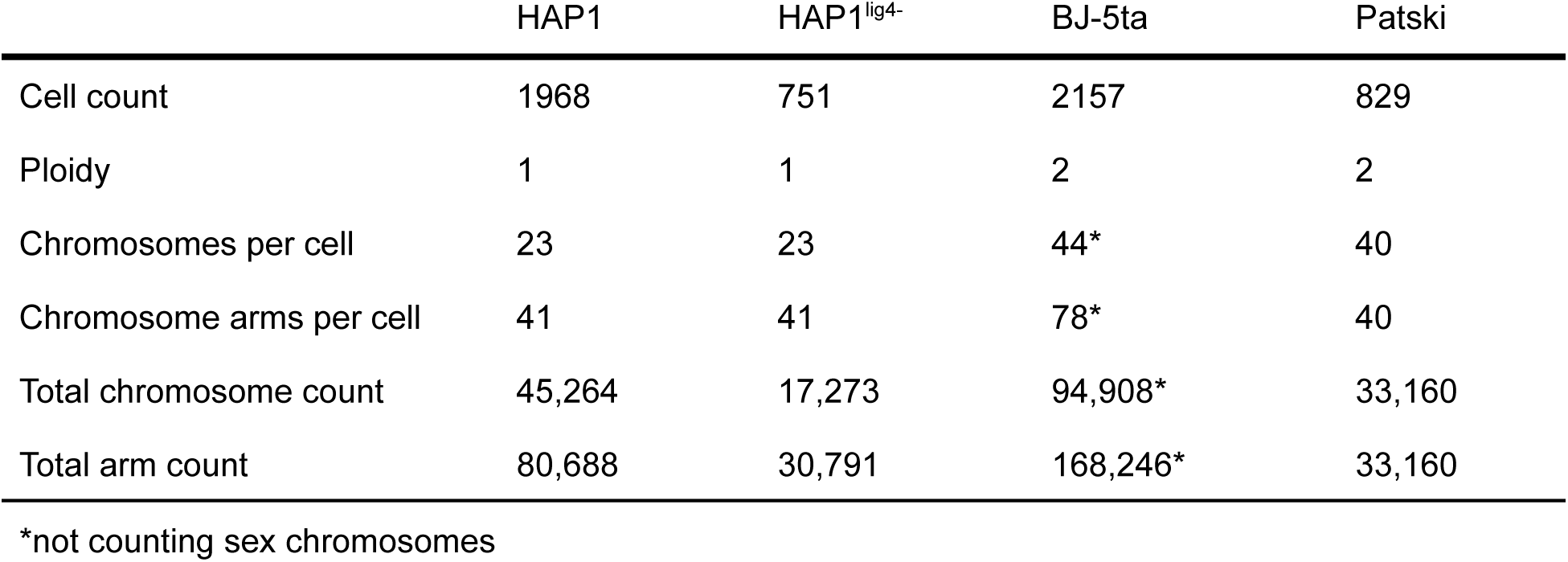

## Supplemental text

### Expansion of sci-L3 to Strand-seq

The quality of strand information obtained for each cell is dependent on the levels of BrdU incorporation within a single division. Insufficient incorporation of BrdU leads to incomplete nascent strand removal that produces interspersed regions of reads on both Watson (W) and Crick (C) strands. Even when achieving near absolute levels of nascent strand removal, the residual amounts form a continuous background throughout the genome. The level of background can be estimated from regions with a majority of only W or only C reads, where for diploids the same template strands are inherited from both parents (WW or CC regions). Cells that have entered S phase before the addition of BrdU will contain a punctate background corresponding to regions with no BrdU incorporation. On the other end, cells that have started incorporating BrdU a second time during the next S phase will contain regions with no reads (1). Aside from the quality of labeling, we also found that excessive soft-clipping of reads performed by BWA creates an additional source of background. By removing the excessively soft-clipped reads we observed a significant decrease in background levels (p-value: 2.02824e-18; Mann-Whitney U) (FigS3A). Ultimately, we devised a set of filters based on the adjusted levels of background, overall strand state percentage and evenness of coverage to eliminate cells with low quality strand information (FigS3-5).

### Sci-L3-Strand-seq enables accurate phasing of heterozygous variants and detection of LOH

We phased SNVs covered by WC reads with StrandPhaseR (2–4) that produced disjoint sets of either one or both SNVs getting phased at each site; consequently, 59.2% (13, 315, 574 SNVs out of 22, 507, 199) of sites in Patski and 43.3% (930, 997 SNVs out of 2, 149, 131) in BJ-5ta required the second haplotype to be inferred from the reference (FigS12A-C). Unexpectedly, we also found an average 1.3% and 6% of heterozygous sites with the same variant assigned to both haplotypes in Patski and BJ-5ta cells respectively, which we labeled as conflicting sites and excluded them from downstream analysis. After inference and removal of conflicting sites, the revised phasing accuracy for Patski cells ranges from 90.7% (chr14) to 98.9% (chr10), with an average accuracy of 98.1% (22, 191, 474 phased SNVs) (Fig3B and FigS12D-G). Additionally, we noticed that despite a 97.2% accuracy, chr4 in Patski cells had the lowest percentage of phased SNVs, at only 14% of total SNV sites, due to a widespread LOH that greatly diminished the number of heterozygous cells with Watson-Crick regions for chr4 phasing. Since the majority of chr4 was from a single haplotype, we were able to manually infer the other haplotype using the reference SNVs, boosting chr4 phased SNVs from 82, 448 to 1, 576, 283 (FigS12H-J). Lastly, we managed to attribute a significant source of the drops in accuracy and conflicting sites to inversions and centromere-proximal regions (p-value: 0.001; Permutation test; conflicting sites correlation coefficient 0.24 in Patski and 0.29 in BJ-5ta) (FigS13A-C). Using a combination of annotations lifted over from mm9 and our inversion calls, we corrected a reference inversion on chr14 and an inversion on chrX (5), both of which increased our overall phasing accuracy (FigS12K). For LOH detection in Patski cells, we were able to identify 1653 regions with extreme haplotype fractions from whole chromosome arms and 1063 from segmentation by strand state (TabS8 - raw LOH calls), which can be merged to 392 LOH calls with 404 corresponding rescues (TabS10). In BJ-5ta cells, we identified 166 regions with extreme haplotype fractions (a union of 94 regions from breakpoint segments and 92 from chromosome arms), and 62 from segmental rescues, a total of 228 as mentioned in the Main text (TabS9 - raw LOH calls).

### Strand switch- and clonal structure-based segmentation with machine learning classifiers facilitate accurate annotation of SVs

For HAP1 and HAP1^lig4-^ cells we implemented separate classifiers for gain and DTZ that achieved a 0.99 accuracy using a 70:30 training to test data split (FigS16A). DTZ was the most accurately identified event across both haploid and diploid cells (FigS16A, FigS16A-D), relying primarily on the centered RCC score (FigS16B). Gain classification in haploid cells relied strongly on regional strand states (FigS16B; final states - see methods), which can make them susceptible to misclassification in the event of missed breakpoint calls. In other words, having both WC strands indicates at least two copies of chromosomes, i.e., a gain in a haploid. However, a chromosome with a missed SCE call can also have both W and C regions, and thus an overall WC chromosome state. To avoid such false positive calls, we devised a bin-based method to evaluate whether regions are a composite of discrete strand states that enabled us to filter calls in post-processing (FigS16C). Despite having trained independent SVM classifiers for HAP1 and HAP1^lig4-^, we chose to only use the HAP1 classifier to ensure any observed differences were not stemming from training different SVMs (TabS12).

For BJ-5ta and Patski cells, we created additional multiclass classifiers for LOH events that achieved a 0.98-0.99 accuracy (FigS17A-D). The gain classifier used the same nine features as LOH with the addition of strand state percentages and the removal of regions previously identified as LOH, achieving an accuracy of 0.971-1 (FigS17A-D). The strand state percentages were key to identifying tetraploid-like cells (4n) in BJ-5ta (FigS17D, FigS20A and D). Regardless of the cells having all the characteristics of tetraploid cells with a mixed strand state percentage below 75% (FigS20D), we cannot confidently rule them out as doublets. It was not possible to have the same gain and LOH classifiers for BJ-5ta and Patski cells due to the disparity in SNV densities between the two cell lines.

### Haplotype-aware segmentation addresses lack of strand switch in cnLOH, enabling their identification, and improves other SV calls

We found most of the new gain annotations enabled by haplotype-aware analyses to be trisomies with missed SCE calls (5 out of 7 manually examined events in Patski cells). Overall, trisomy with an SCE constitutes a small proportion of all trisomy events in Patski cells (146/1652 events in Patski cells), suggesting missed SCEs are the leading cause of unannotated whole chromosome gain. Similarly, haplotype-aware segmentation also enabled us to identify segmental high-copy number gain that would otherwise appear as a whole chromosome gain due to the information being averaged across an entire chromosome (FigS19D).

### Clonal lineage suggests recurrent SVs

We used the same Canberra distance for the diffusion map as for hierarchical clustering of our SVM annotations (FigS23A). The first diffusion component (DC1) separated out clonal populations rooted by segmental loss on chr2 (Fig3F top left panel). Chr4 LOH is present in the other branch in 85% of Patski cells, making it the dominant backbone event around which further mutational events arose. The second diffusion component (DC2) describes the events surrounding the chr4 LOH, broadly separating out cells with chr12 monosomy and chr5 trisomy on one branch and cells with chr19 trisomy onto another (Fig3E top right and bottom panel). Projecting chr4 LOH onto the diffusion map reveals no obvious structures that identify a divergence point between monosomy and UPD with respect to the other mutational events (FigS23C left panel). For chr4 monosomy itself, the detection of both haplotypes suggests it occurred at least twice, while the domination of one haplotype (C57BL/6J) suggests possible selection bias (FigS22D). We observed a similar haplotype imbalance (*M. spretus*) for the monosomy on chr12 and trisomies on chr3 and chr19 (FigS22D). Interestingly, chr12 monosomy and chr19 trisomy are almost entirely mutually exclusive (Fig3E and FigS23B). An alternative to the gradual accumulation of events are punctuated bursts, previously described in human cancers and genetic disorders, that could have produced an initial cell containing multiple trisomies (6, 7). Unlike LOH, trisomy can revert to normal heterozygous disomy and produce the stepwise clonal structure we observe surrounding chr4. However, reverting to a disomic state should be equally likely to result in UPD, which we rarely observe on chromosomes with trisomy. Additionally, for the trisomies on chr5, the choice of which parental chromosome is doubled appears random with an almost perfect balance of both within an otherwise clonal population (Fig3D and FigS22D). The chr5 balance contrasts the biased haplotypes of every other mutational event that we examined in Patski cells (FigS22D). One explanation for the lack of haplotype selection could be that chr5 trisomies are recently acquired events. Lastly, we focused on DC4 as it provides a better separation of trisomies, while DC3 primarily separates out cells with segmental gains (FigS23B). These cells with multiple trisomies are likely to have originated from the chr4 LOH clones containing pre-existing chr3, chr5, and chr19 trisomies and having subsequently acquired the unique trisomies on chr6, chr10 and chr17 (Fig3D). The founder populations of trisomies on chr15 and chr19 provide the first evidence of the independent origin and recurrent nature of events in Patski cells.

Likewise, the trisomies on chr1, being within separate clonal populations with irreversible LOH events and containing both haplotype combinations, provides evidence of their independent origin.

### SVs alter the rate of SCEs

Breakdown of cells into five categories allowed us to disentangle the contribution of the different events to the rate of SCE in the following manner: 1) Cells with only a UPD on chr4 have a similar rate of SCEs to controls, which consist of cells without chr4 UPD, monosomy or chr12 monosomy (p=3.15x^-01^; Mann-Whitney U). 2) Cells with only chr4 monosomy again showed a significant reduction in SCE (p=1.15×10^-05^; Mann-Whitney U), implying the duplication of chr4 (UPD) restores the rate of SCE to normal levels (FigS24A). The addition of chr12 monosomy appears to have the same restorative effect for cells with only chr4 monosomy. 3) The combination of chr4 UPD and chr12 monosomy has an additive effect that significantly increases SCEs by 1.25-fold over cells with chr4 UPD alone (FigS24A) (p-value: 1.14e-04; Mann-Whitney U).

### Measuring the rates of SVs

We noticed that haploid cells contain a substantially higher amount of whole chromosome DTZ compared to diploid cells. As cells without whole chromosomes are unlikely to be viable, these could represent a snapshot of cells about to undergo apoptosis, or alternatively, they may correspond to the single copy of chromosome sequestered in micronuclei. With the various detergent and heat treatments in sci-L3, it is possible micronuclei dissociate and are lost during wash steps.Chromosomal (fragments) in micronuclei have a high chance of subsequent loss (8). Our characterization of DTZ could thus be biologically meaningful. In diploid cells, segregation errors of broken or whole chromosomes that produce micronuclei should not lead to a concerted loss of both homologs, yet in both Patski and BJ-5ta cells we still observed DTZ. We found 74 Patski cells containing a whole chromosome or arm DTZ, including 63 near-haploid cells for which micronuclei could still explain DTZ. In the remaining 11 cells we found all had a fully matching complement of monosomy or arm loss events in another cell. For BJ-5ta we found 3 cells containing a DTZ, out of which one had a matching monosomy in another cell. The observation that DTZ in cells with matching loss events in the population suggests DTZ in diploid cells requires a loss intermediate and thus is a two-step event.

### Opposite strand reads identify inversions

As an example of inversion identification from the opposite strand read, we plotted the Watson reads from an otherwise Crick-Crick chr8, revealing a large inversion at the left end of the chromosome (Fig5A left panel). Similarly, for cells with both chromosomes in the Watson strand orientation (WW), the chr8 inversion appears on the Crick heatmap as a column of reads above the background across all cells (FigS26A top panel). We used the heatmap read count matrices to generate genome-wide pileup tracks across all cells with WW or CC chromosomes that showed inversions as peaks and enabled us to identify their coordinates (see methods). With the pileup tracks of chr8 from CC strand state cells we mapped the BJ-5ta cell specific inversion to a 4Mb region (Fig5A right panel). The tracks also highlighted the inversion is heterozygous with a 1 to 1 ratio of Watson to Crick reads. Within WC strand state chromosomes, a heterozygous inversion is visible as a region with a WW or CC strand state, while for a homozygous inversion only a haplotype switch would be visible, since the inversion itself would still contain both strand orientations (FigS26A bottom panel).

### Identifying large and sub-clonal inversions from breakpoint hotspots

Calling inversions by examining double breakpoint calls is also helpful in filtering breakpoints that would otherwise be called as SCEs. SCEs are not clonal in nature but inversions are and in theory should be called as double breakpoints. However, in practice, breakpoints are not always perfectly called and have limited resolution of 128-255kb. For short inversions, <200kb for example, both or one of the strand switches could be easily missed from calling. Medium sized inversions could be subject to the 2Mb and 10Mb size filters we implemented for adjacent SCEs in haploid and diploid cells respectively (FigS6C). Therefore, for inversions with sizes <2Mb (or <10 Mb for diploid cells), if no or both strand switches flanking in the inversion is called, there will be no breakpoints called in that region, and thus this region will be called using opposite strands in otherwise fully WW or CC chromosomes. However, if only one of the two strand switches flanking the inversion is called, it will be first mis-classified as a single breakpoint event rather than an inversion. The two strand switches flanking the inversion could be randomly missed from calling, particularly in low-coverage cells. Therefore, when we aggregate breakpoints in all the cells, these inversions shared by a population of cells would show up as double breakpoint hotspots. Similarly, inversions >2Mb (or >10 Mb for diploid cells) would show up as adjacent SCE hotspots. To identify such “twin” hotspots, we could examine either the pileup of all “SCEs” (before filtering out SVs, FigS7) or SCEs from chromosomes with at least two breakpoints (Fig5C-D)

### Strand-state correlation identifies known and novel translocations

The translocation and inverted duplication on chrX in HAP1^lig4-^ cells are both absent from the parental HAP1 cells (FigS7F and FigS23B) and based on the ubiquitous nature of the two SVs it is possible they arose during the CRISPR-Cas9 editing. Examining the correlation gradient of the chr13-chrX translocation suggests the chrX p-arm segment is fused to the short arm of the acrocentric chr13 rather than to the CRISPR-Cas9 cut site, which is located within the terminal end of chr13 (Fig6D). Interestingly, the other two translocations also involve acrocentric chromosomes, suggesting acrocentrics may be more susceptible to translocations. A study examining chromosomal instability after CRISPR-Cas9 editing in HAP1 cells observed a high rate of aberrations of chr13, such as chromosome fusions, that fits with our observations (9).

### Resolving complex events provide new insights into the HAP1 disomic region

We can confidently rule out 1) a heterozygous inversion on the chr15 copy, since if the inversion was on chr15, the strand state of the inversion would be the opposite of the state of chr15 in cells with different strand states on chr15 and chr19 (WW or CC), while remaining the same in cells that have the same strand state across both chromosomes (WC); and 2) a homozygous inversion on both the chr15 and chr19 copies, since a homozygous inversion would not be visible in the disomic regions with a WC strand state and would only appear in cells that share that same strand state across both chr15 and chr19 (FigS28).

### Supplemental discussion on new bioinformatic tools developed for sci-L3-Strand-seq analyses

We summarize new bioinformatics tools developed and employed to process sci-L3-Strand-seq data and link them to sections in the **Methods**. We note tools used in the iterative process (marked with “iterative”) between single-cell and population level analyses, and tools that are also applicable to sci-L3-WGS (marked with “WGS”). At the aligned read level, we first implemented a filter to remove excessively soft-clipped reads to reduce background noise based on strand directionality (**Read processing and alignment**). We then used both background (below 8%) and strand-neutral state (15% at most for haploid cells and 75% for diploid cells) to filter at the cell level to pick successful Strand-seq libraries (**Sample selection and cell filters**). We also mark high spike cells for haploid in SV annotation (**Identification of high spike cells**). Strand switches are called in single cells with successful Strand-seq library preparation. We developed five layers of breakpoint filter (**Breakpoint calling and filtering**) including: 1) a region filter (“iterative”) with pre-existing SV identified by examining strand switch hotspot not due to SCE; 2) clustered breakpoint filter (“WGS”); 3) strand-state filter; 4) CEN-proximal filter; and 5) rescue of clustered events by checking strand state for outermost breakpoints. Most strand switches map SCEs, but they could also define boundaries of segments as candidates for mutational SVs.

Phasing facilitates SV calls. We implemented phasing with StrandPhaseR, but also improved phasing accuracy with resolving haplotype conflicts (predominantly near centromeres), leveraging cells with clonal LOH as such events expose phased sites, and correcting over inversions (see **Phasing**). For counting copy number, we implemented digital counting (**Digital overlap counts**) of Tn5 insertion patterns, which involves: 1) a rigorous deduplication scheme to enable overlapping read analysis (“WGS”); 2) extracting strand-specific overlaps; and 3) extracting haplotype-specific overlaps (“WGS”). We further identified and developed tools to extract additional features that improves SV annotation with SVM, including: overall and strand-specific overlaps, strand states, percentage of regions in WC state (a single value for each cell), percentage of regions in unassigned strand state (a single value for each cell), RCC score, RCBC score (a single value for each cell), read depth for the cell (a single value for each cell), mappable width of a segment, and haplotype fraction. We also added haplotype-aware features including haplotype-specific overlaps and haplotype-specific strand state. We experimented on methods to build clonal maps (**Clustering and diffusion maps of annotated events**) with various distance measures (“WGS”, and “iterative” for validating SV calls as TP events should be more clonal). We provide a binned approach and a “double SCE hotspot” approach to call inversions **(Inversion annotation**, “iterative” as these help filter SCEs). Plotting negative correlation of strand state also helps validate large inversion. We call translocations by examining strand-switch hotspots. We further verify translocations by co-segregation of the same strands, and positive correlation of strand state in otherwise unlinked regions. After SV annotation, we devised SCE filters (“iterative”, see **Breakpoint region filter and SCE filter**) to remove apparent “SCE” calls due to subclonal detentions, inversions, and segmental gains.

## Supplemental figures

**Figure S1.**
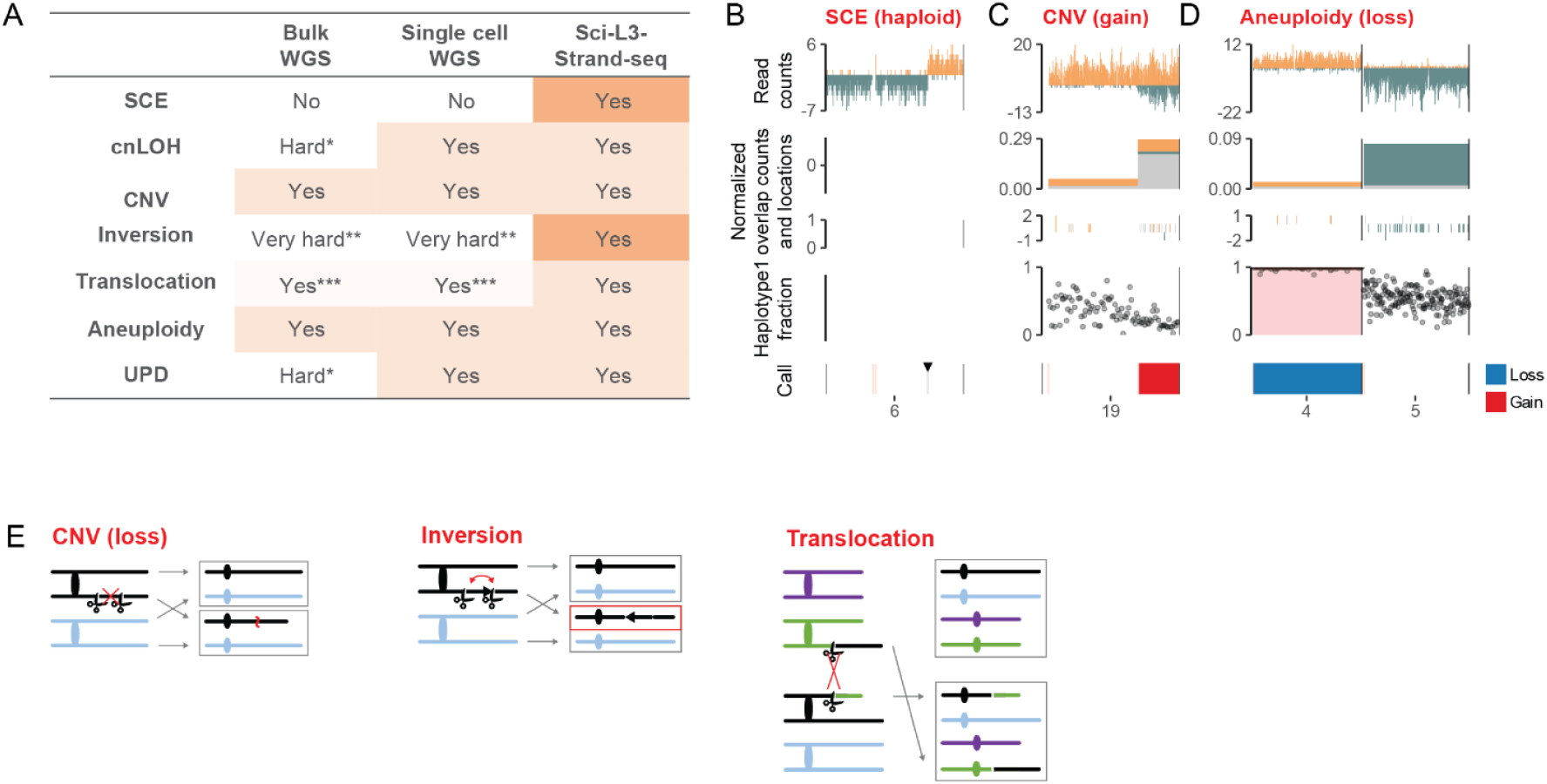
Mapping of mitotic crossover outcomes with additional examples of SCE, CNV and aneuploidy. **(A) The ability of different assays to detect the seven outcomes of mitotic crossovers.** Sci-L3-Strand-seq uniquely detects SCE, inversion and balanced translocations. Note, for reliable inversion and translocation detection, multiple cells with the same event are required. * Bulk WGS cannot distinguish reciprocal events if both daughter cells are sequenced, as every position will appear heterozygous. It is possible to clone each daughter cell, but at a considerable expense of time and labor. **Inversions do not alter heterozygosity or copy number in the resulting daughter cells and can only be detected with junction-spanning reads. ***Unbalanced translocations cause CNV, detectable with WGS. However, like inversions, balanced translocations do not alter heterozygosity or copy number and can only be detected with junction-spanning reads. **(B-D) Panel plots for example error-prone mitotic crossover events.** Each point for the haplotype1 fraction is the phased allelic fraction of approximately 50 binned SNVs for Patski and 15 for BJ-5ta. Black triangle denotes the location of SCE, while the red vertical line represents the centromere. **(B) An example of SCE in haploid cells.** In haploid cells (HAP1), an SCE produces a complete switch from one strand (C) to the other (W). **(C) An example of a segmental gain on chr19 in a Patski cell.** A substantial increase in the number of opposite strand overlapping reads (gray bars; Watson same strand overlaps: dark yellow bars; Crick same strand overlaps: dark green bars), accompanied by a skew of the haplotype ratio, is observed over the region of the segmental gain (red). The haplotype skew reflects the 2:1 haplotype1 to haplotype2 ratio of the segmental gain. **(D) An example of monosomy on chr4 (blue) in a Patski cell.** Chr5 from the same cell is included to highlight the lack of same strand overlapping reads (dark yellow bars expected at equal frequency to chr5 dark green bars) along with only a single copy of the homozygous chromosome (LOH - shaded pink). **(E) CNV, inversions and translocations as a result of end joining require two initiating DSBs.** Same as in Fig1C-E, with white bar and scissors indicating positions of the DSBs.

**Figure S2.**
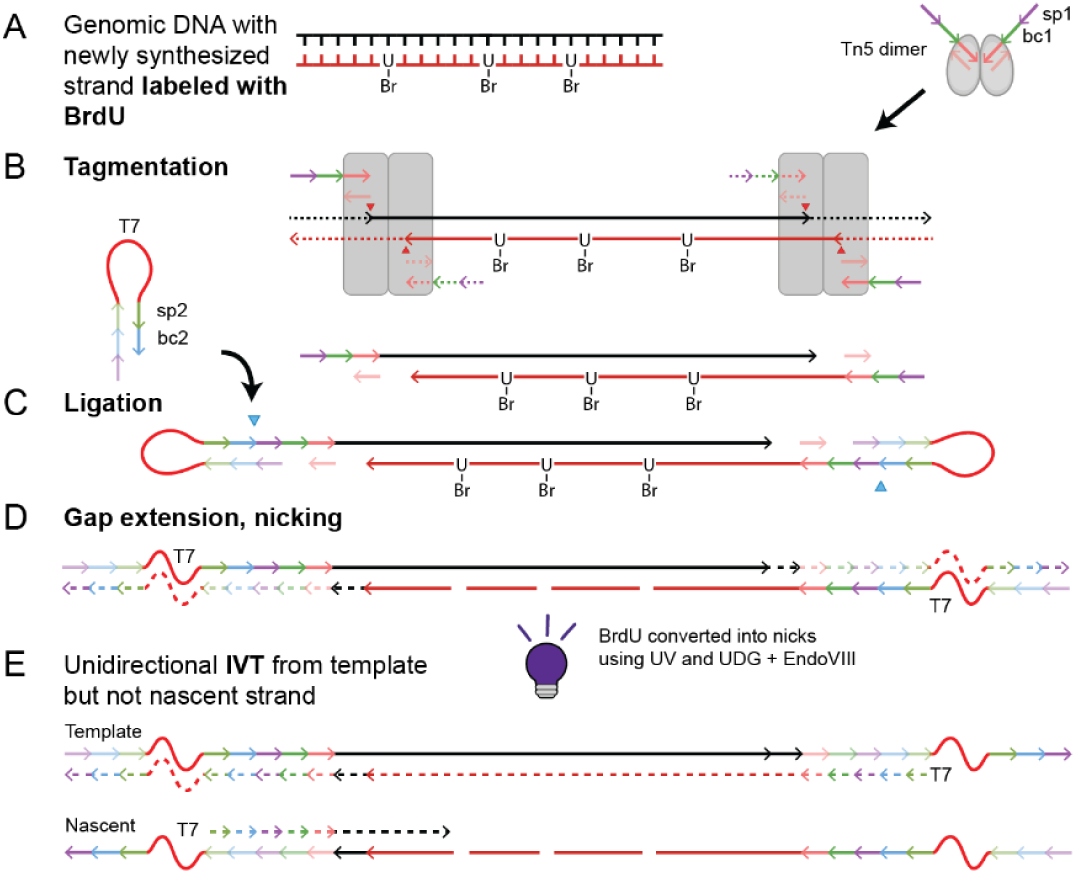
Molecular structures at each step of sci-L3-Strand-seq. (A) BrdU is added into the cell culture media and cells are maintained for a single cell cycle. During replication the BrdU is incorporated into the newly synthesized strand, and cells are harvested in the next G1. (B) Tn5 inserts the first round barcodes (bc1) with a spacer (sp1) that will be used for ligation. Tn5 transposase catalyzes staggered dual nicking (off by 9nt) and single strand transfer such that strand directionality of the replication template strand is maintained. (C) After pre-annealing to form a hairpin-loop, the ligation oligo is ligated via an overhang with sp1. Sticky end ligation again maintains strand directionality. The ligation oligo contains a T7 promoter for in-vitro transcription (IVT), a second round barcode (bc2), a spacer (sp2) that is the priming site for downstream second strand synthesis (SSS) step (after E, additional detail can be found in (10)). (D) Residual gaps from Tn5 insertion are filled and the polymerase further makes the T7 promoter double stranded. Only a single successful ligation is required to enable downstream IVT. The nascent strand is nicked at the sites of BrdU incorporation, using UV and the Hoechst dye for debromination and uracil DNA glycosylase and endonuclease VIII for excising the U, effectively removing it from downstream steps. (E) IVT is performed unidirectionally from the replication template strand, but not from the nascent strand, preserving strand information. The subsequent RT and SSS with a UMI and third-round barcodes are performed as previously described in the original sci-L3 method. Both are mediated by self-looped priming and/or annealed primers to maintain strand directionality.

**Figure S3:**
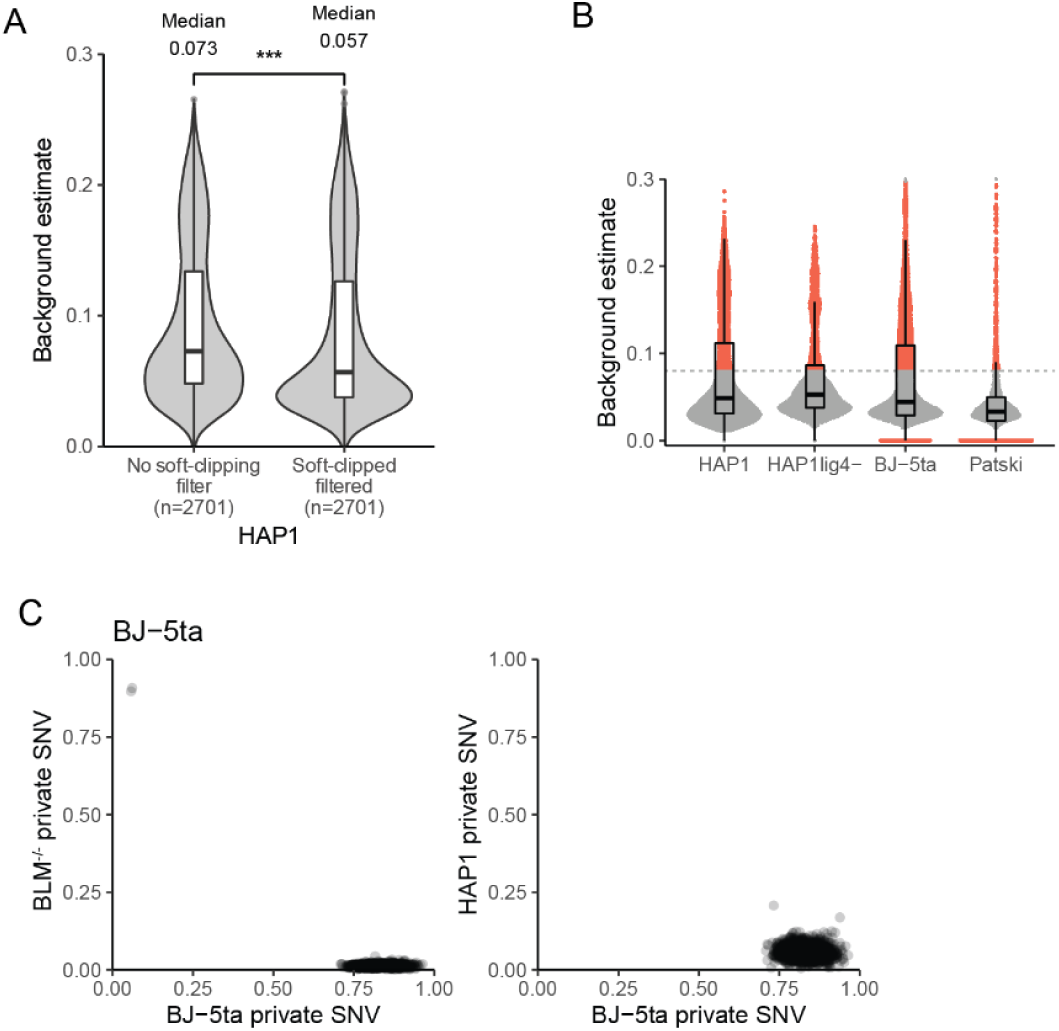
Library selection and filtering. **(A) Impact of removing excessively soft-clipped reads on background estimates.** Reads that had more than half of its length soft-clipped (ratio of 0.5), were removed from downstream analysis (soft-clipped filtered). Only data for HAP1 cells shown. Significant difference level (*** < 0.001, ** <0.01, * <0.05) was calculated using Mann-Whitney U (p-value: 2.03×10^-18^). **(B) Background estimates for each cell line.** Each dot represents a single cell. Cells with a background above 0.08 or exactly 0 (unable to calculate background due to lack of WW or CC state chromosomes) were removed from downstream analysis (red). Cells in gray were used in downstream analysis. Number of cells filtered by background for each cell line is provided in TabS1. **(C) Cell identity based on private SNVs.** To filter out mislabeled cells from mixing experiments, the cell identity of each human cell line was additionally verified using SNVs private to each cell line. Only cells filtered for coverage, background levels and percentage of WC chromosomes are plotted. Two cells misclassified as BJ-5ta cells, but containing private SNVs to a *BLM*^-/-^ cell line were excluded from downstream analysis.

**Figure S4:**
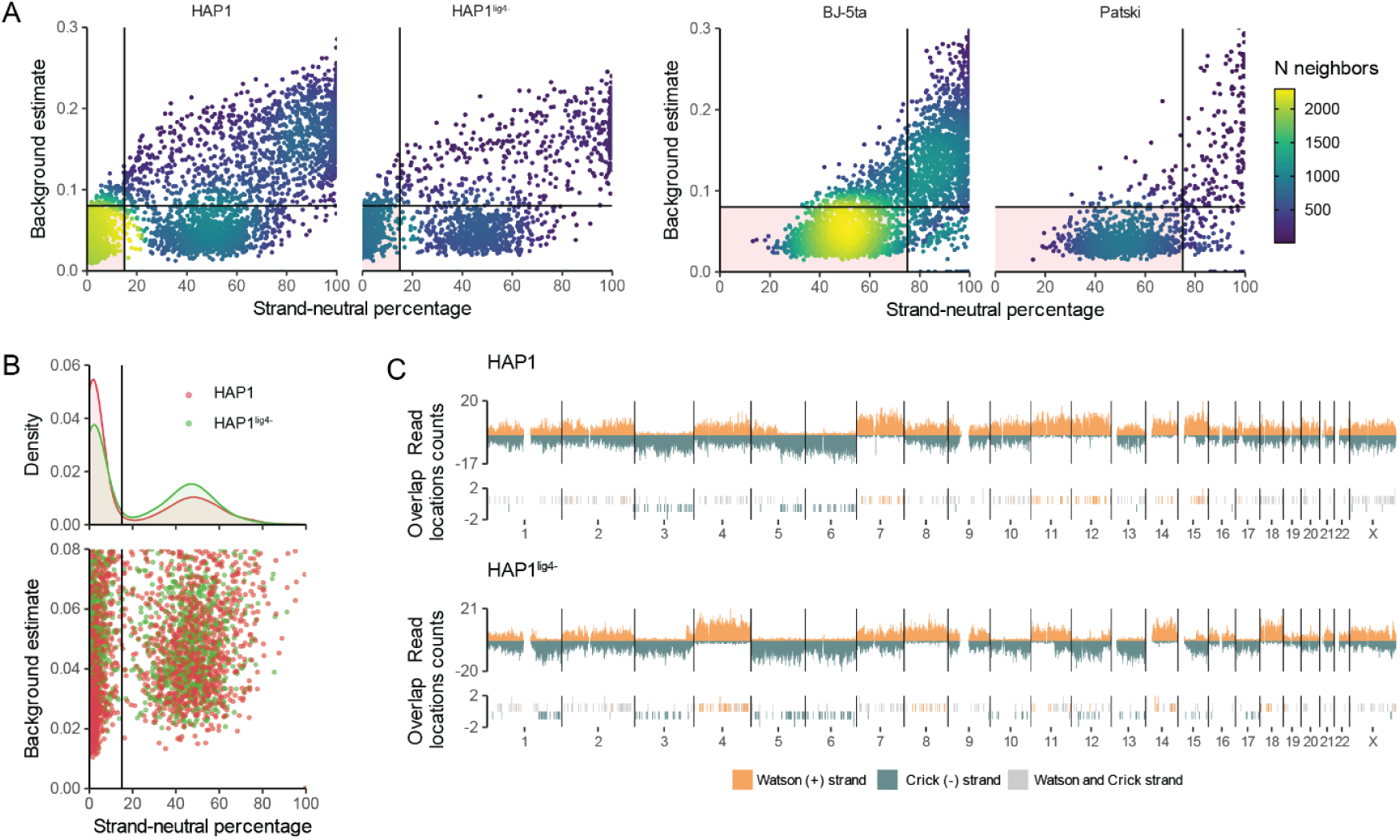
Filtering of cells with high strand-neutral percentage. **(A) Filtering cells based on strand-neutral percentage and background estimate.** We defined regions that have a WC strand state and/or not biased enough to be assigned to WW or CC states as “strand-neutral”. Each dot represents a single cell. Haploid cells with less than 15% strand-neutral regions (vertical black line) and background estimate of less than 0.08 (horizontal black line) were retained for downstream analyses (pink shaded quadrant). Density of points is shown as the number (N) of neighbors. Diploid cells with less than 75% of strand-neutral regions (vertical black line) and background estimate of less than 0.08 (horizontal black line), were retained for downstream analysis (pink shaded quadrant). State percentages shown were calculated before breakpoints filtering. Cells with a high strand-neutral percentage have a uniform distribution of both Watson and Crick reads across the entire genome, suggesting the BrdU labeling or nicking failed to remove the nascent strand. Number of cells filtered by strand states is provided in TabS1. **(B) Haploid cells with diploid levels of strand-neutral states.** Each dot represents a single haploid cell. Haploid cells that have between 15% (vertical black line) and 75% of strand-neutral states and less than 0.08 background estimate resemble the strand-neutral distribution observed in diploid cells, and are thus removed from downstream analyses. HAP1^lig4-^ cells have slightly more diploidized cells. **(C) Example of HAP1 and HAP1^lig4-^ cells with high levels of strand-neutral state chromosomes**. The disomic region on chromosome 15 is still present, confirming the cell identity. High number of overlapping fragments across all chromosomes supports the known tendency of these cells to revert to diploids. It is reasonable to expect a different number of Tn5 insertions between individual cells that results in different levels of coverage. The additional observed equal coverage within these cells suggests that the high overlapping read counts and WC regions are not the result of doublets, but rather single cells that have become diploid.

**Figure S5:**
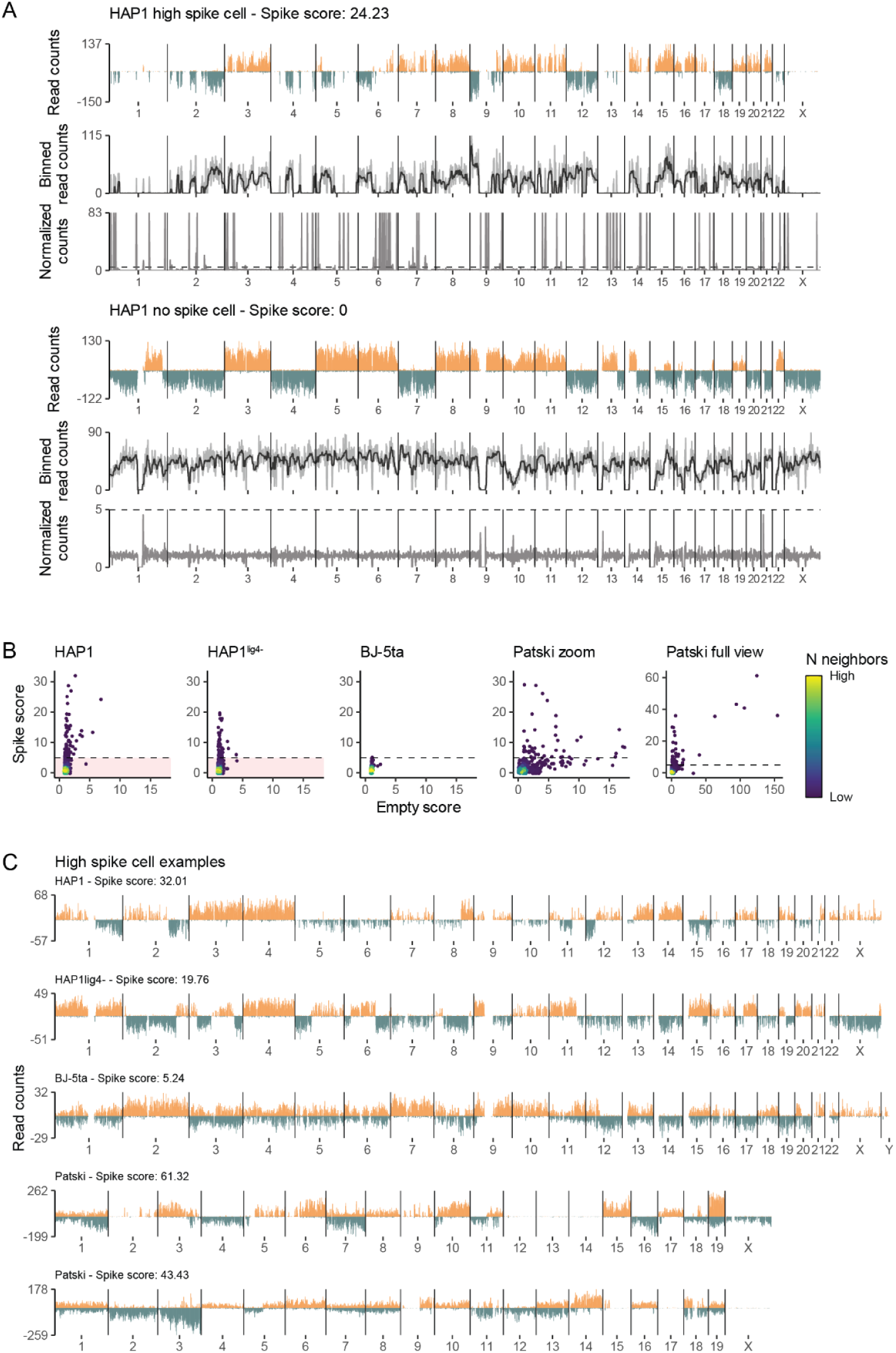
Identification and removal of cells with high levels of spike regions. **(A) Example cells with and without high levels of spikes.** The top panel shows a cell with a large spike count. The bottom panel shows a more representative cell with very few spikes. Spikes within cells are identified using a running median, shown as a black line in the middle plot within each panel. Using the running median, read counts are normalized to highlight spikes as shown in the bottom plot within each panel. The dashed line in the bottom plots of each panel represents our chosen threshold for calling spikes. Reads were counted within 1 Mb sized bins. **(B) Filtering of high spike cells.** Haploid cells with a spike score above (>=) 5 (dashed line) were excluded and only cells in the pink quadrant were used for downstream analysis. Only a single cell fell above the threshold in diploid BJ-5ta cells, while Patski cells had an unusually high empty score that can be attributed to a large number of haploid-like cells. Because of the ability to distinguish background spikes from real events with SNVs, the lack of spikes in BJ-5ta cells, and a biologically relevant reason for a high empty score in Patski cells (haploid-like cells), we chose not to apply the spikey filter in diploid cells. **(C) Examples genome-wide plots of cells with spikes from each cell line.** Reads counts are plotted at 200Kb bin resolution. We include a near haploid cell (above) and a diploid cell (below) for Patski high spike examples.

**Figure S6.**
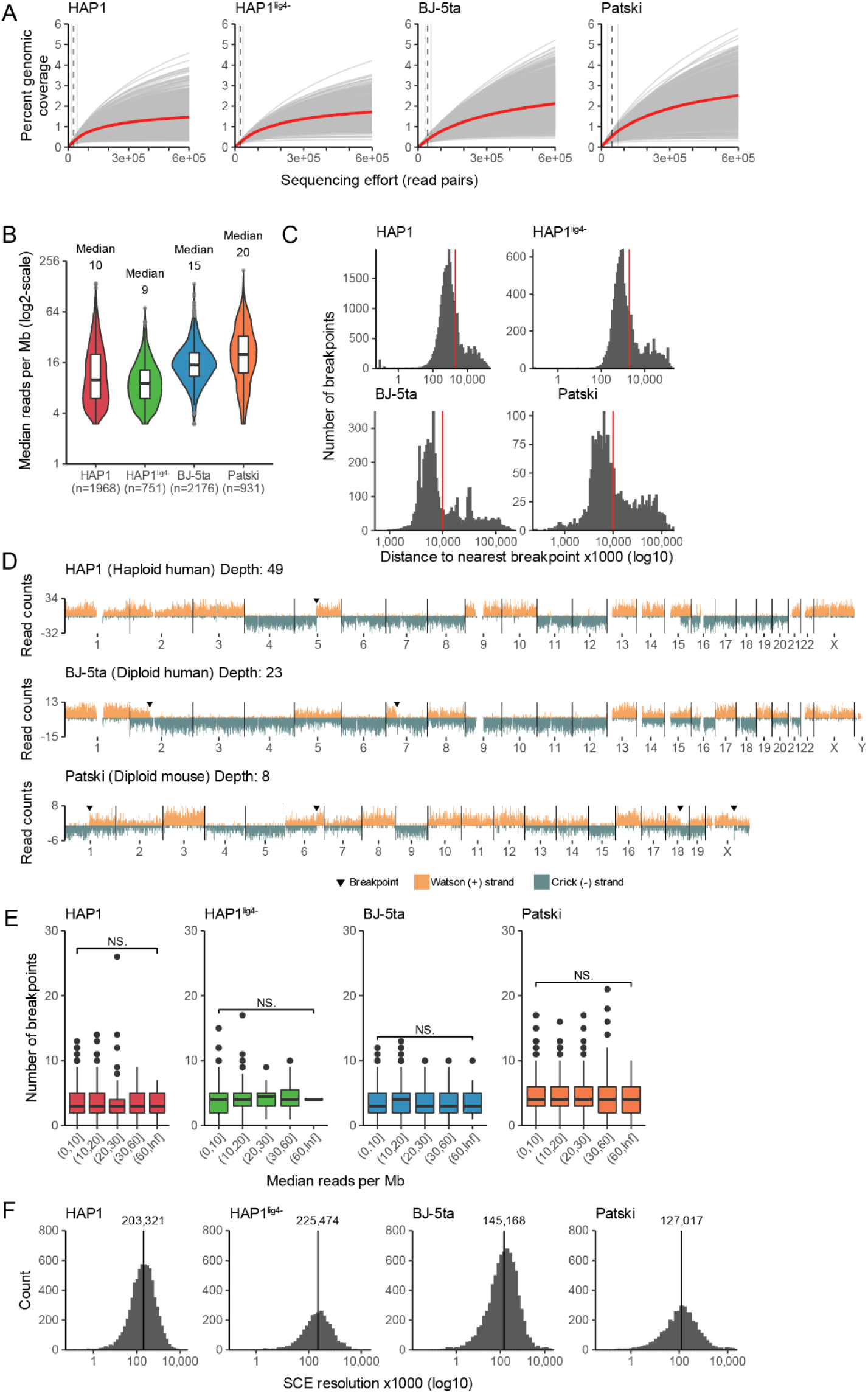
Impact of sequencing depth on SCE calls. **(A) Sequencing effort required to achieve coverage saturation for each cell.** Each gray line is the predicted sequencing effort required as a function of genomic coverage percentage for a single cell. The red line represents the smoothed conditional mean of all single cells. Vertical dashed line represents the median read pairs actually sequenced, showing that our sequencing effort is sub-saturation (adjacent lines represent the 25th and 75th quantile). **(B) Distribution of read coverage (median reads per Mb) within each sample.** Median absolute deviation (MAD) values for HAP1: 7.4, HAP1^lig4-^ 4.4, BJ-5ta: 7.4, Patski: 14.8. A median of 9 reads per Mb roughly corresponds to 0.26% coverage. **(C) Distances between nearest breakpoint events.** For haploid cells, breaks within 2Mb of each other were filtered (red vertical line). For diploid cells, the filtering distance for nearest breaks was 10Mb. Cutoffs were chosen based on the bi-modal distributions observed. **(D) Genome-wide plots of W and C reads with different levels of coverage**. W and C reads were counted within 200kb bins. We show example cells with depths of 49, 23 and 8 median reads per Mb.Black triangles denote identified breakpoints. **(E) Number of breakpoints are not significantly different at different levels of coverage.** Comparison of the lowest coverage cells (0-10 median reads per Mb, less than 0.3% coverage) with the highest coverage cells (60+ median reads per Mb, more than 1.8% coverage) was performed using the Mann-Whitney U test (NS.: p-value > 0.05). **(F) Resolution of breakpoints within each sample.** Vertical lines denote median break resolution in base pairs (bp). Breakpoint resolution was determined with breakpointR (11) (MAD HAP1: 212886.5; HAP1^lig4-^: 221783.6; BJ-5ta: 166416.7; Patski: 149988.0).

**Figure S7.**
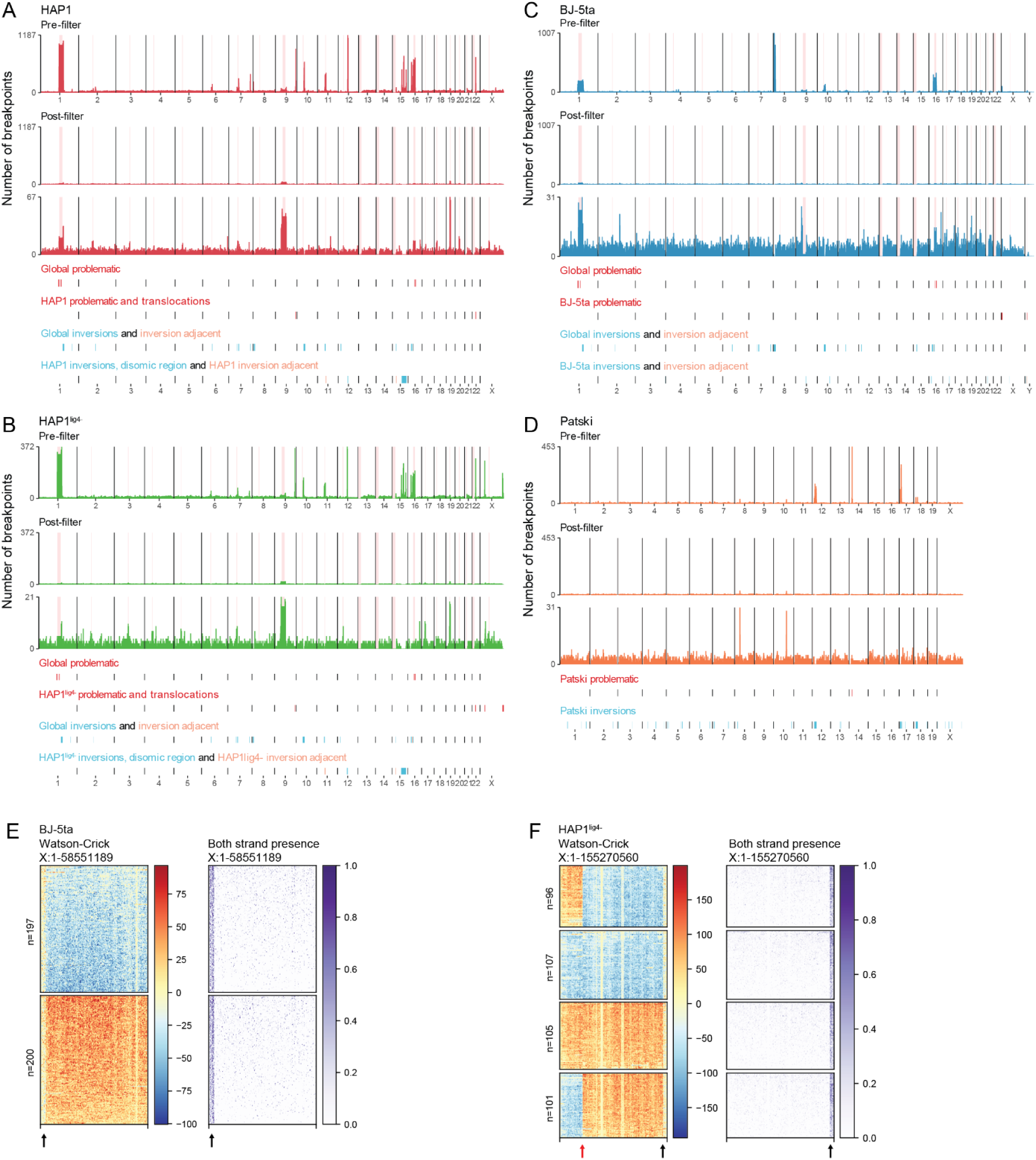
Region filter for breakpoints matching inversions, translocations and problematic regions. (A-D) The number of breakpoints identified by breakpointR without any filters are shown in the top bar plot for (A) HAP1, (B) HAP1^lig4-^, (C) BJ-5ta, and (D) Patski cells. Breakpoints remaining after applying the region, clustered events, strand state, and centromere filters (see **Breakpoint calling and filtering** section in **Methods)** are shown in the post-filter bar plots. Bottom post-filter plot has the y-axis scale adjusted. Note that breakpoints were counted across 1Mb bins without any normalization, which leads to low resolution breaks such as those across centromeres to appear as hotspots. Note, the post-filter raw counts used in Fig2D were additionally normalized. Centromeres are shaded in pink. Coordinates used for the region filter are plotted below the bar plots. The filtered regions (“region filter”) can be separated into several categories, with the global problematic and global inversion (including inversion adjacent) tracks showing regions with breakpoint hotspots that are common to all three human cell lines. We were unable to make these distinctions for (D) Patski as it was the only mouse cell line analysed. Global inversions and global problematic regions stem from issues in the genome assembly such as incorrect orientation of placed sequences with examples on mouse (mm10) chr14 (5). In human cells, the centromeres of chr1 and chr16 are additionally labeled as global problematic regions due to their high rate of false positive breakpoints. For these two centromeres, we performed an additional centromere rescue step that examined the strand states of the flanking arms to determine if a real SCE was present. Cell line specific problematic regions encompass identified translocations in (A) HAP1 and (B) HAP1^lig4-^ cells. Additionally, chrY in BJ-5ta cells contains a problematic region with high false positive breakpoints that could stem from its repetitive nature and incomplete assembly (12). The problematic regions (B) on the rightmost part of chrX in HAP1^lig4-^ and (C) on the left side of chrX in BJ-5ta cells are present within haploid chromosomes despite contain both W and C reads, suggesting these regions are duplicated. In addition to global inversions, we identified cell line specific inversion that were present in the vast majority of cells of a single cell line with notable examples (A-B) on chr12 of HAP1 and HAP1^lig4-^ cells and (C) on chr10 in BJ-5ta cells. We found several breakpoint hotspots without any obvious strand switches that were adjacent to global and cell line specific inversions, suggesting that inversions may be causing false positive breakpoint calls within their proximal area (light brown). Finally, we also chose to filter breakpoints resulting from the known disomic region located on chr15 in (A) HAP1 and (B) HAP1^lig4-^ cell. The list of problematic regions, inversion and translocations with their coordinates is provided in TabS2. **(E-F) Heatmaps of inverted duplications on chrX in (E) BJ-5ta cells and (F) HAP1^lig4-^ cells.** Red arrow highlights translocation, while the black arrows highlight the duplicated regions. Each row represents a single cell plotted at a 200kb bin resolution. Left heatmaps for each figure show subtraction of W read counts (red) from C (blue) for each bin. Right side heatmaps highlight regions with both W and C reads by plotting the ratio between the minimum and maximum read counts for each bin (purple, **Chromosome heatmaps** in **Methods**).

**Figure S8.**
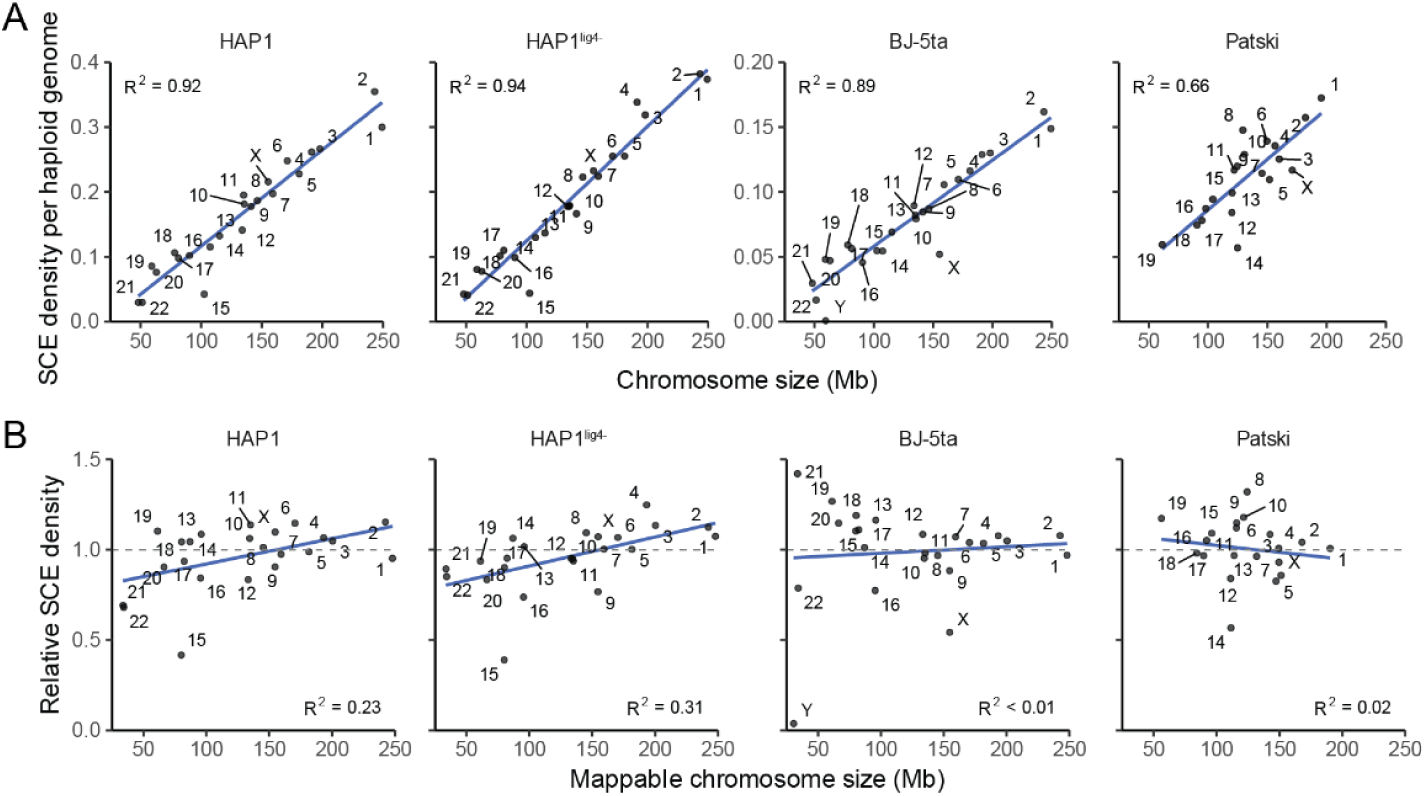
SCE density is directly proportional to chromosome size. **(A) Correlation between chromosome size and the number of SCE.** SCE counts were normalized by the total number of cells and ploidy. Chromosome X and Y stand out as outliers in the male BJ-5ta cell line due to their single copy nature, while chromosome 15 in HAP1 and HAP1^lig4-^ cells stands out due to the removal of breakpoint calls within the known disomic region. **(B) Correlation between mappable chromosome size and the relative number of SCE.** Relative counts were defined as the average number of SCE per chromosome normalized by the mappable chromosome size, over the average number of SCE per genome normalized by the mappable genome size. The mappable chromosome and genome sizes in this instance were adjusted by only removing gaps (Ns and blacklisted regions) from the start and end of chromosomes. The reasoning being, as long as there is enough flanking sequence on either size of an unmappable region, a breakpoint can still be detected. Gray horizontal dashed line represents the expected relative SCE density.

**Figure S9.**
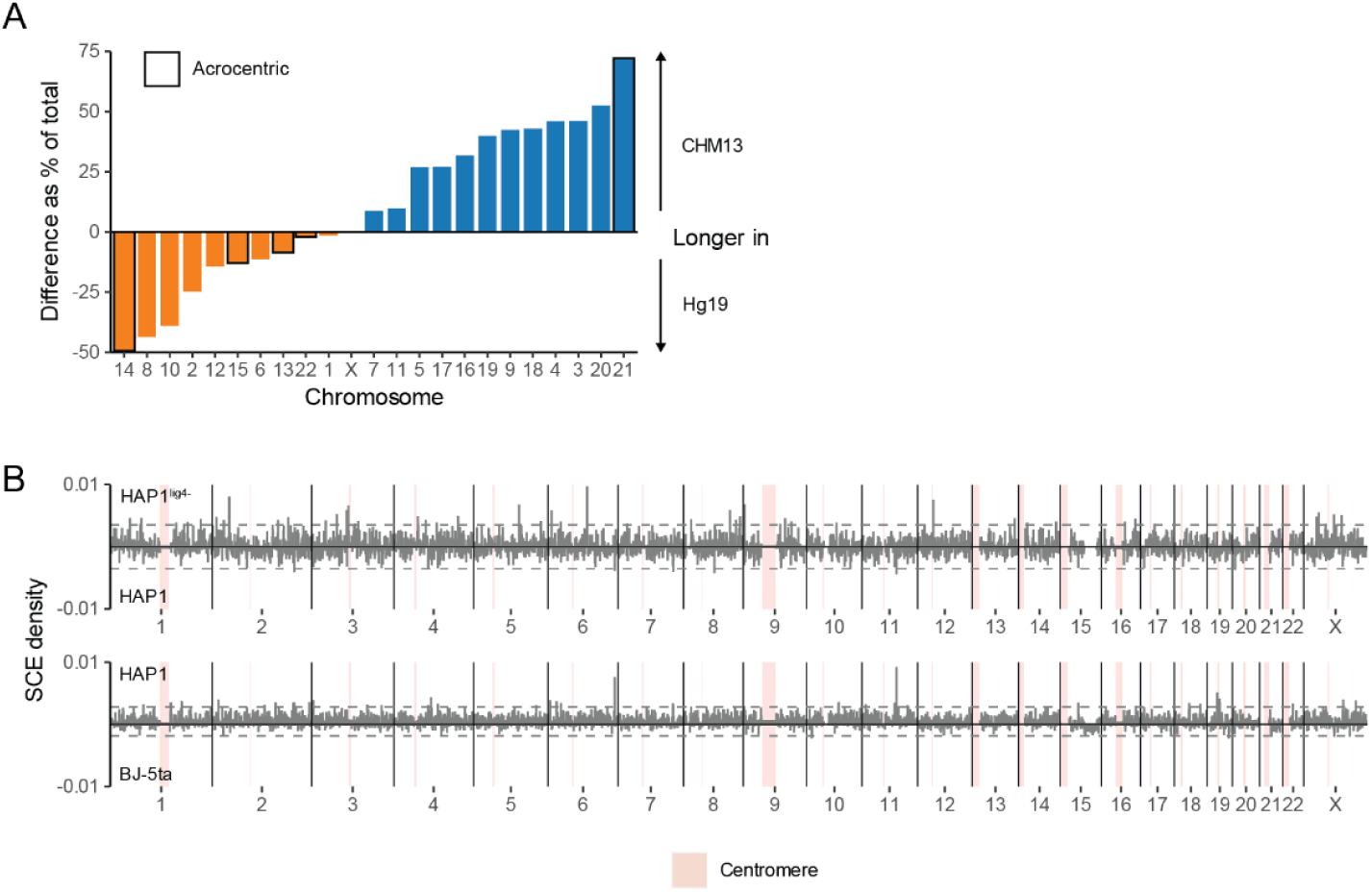
Centromere size disparity and breakpoint density differences between cell lines. **(A) Size differences of centromeres in hg19 versus the T2T-CHM13 human genome reference.** Acrocentric chromosomes are highlighted with black boxes. **(B) SCE density differences between HAP1 and HAP1^lig4-^, and BJ-5ta and HAP1 cell lines.** Densities are plotted at 1Mb bin resolution. The centromere (red highlight) sizes were resized to the T2T-CHM13 sizes to more accurately reflect SCE frequency within the centromeres. SCE counts were normalized by the number of bins each SCE overlaps with, the total number of cells, and the ploidy of each cell line. Gray dashed lines show the median +/- 3 times the median absolute deviation (MAD) for each comparison.

**Figure S10:**
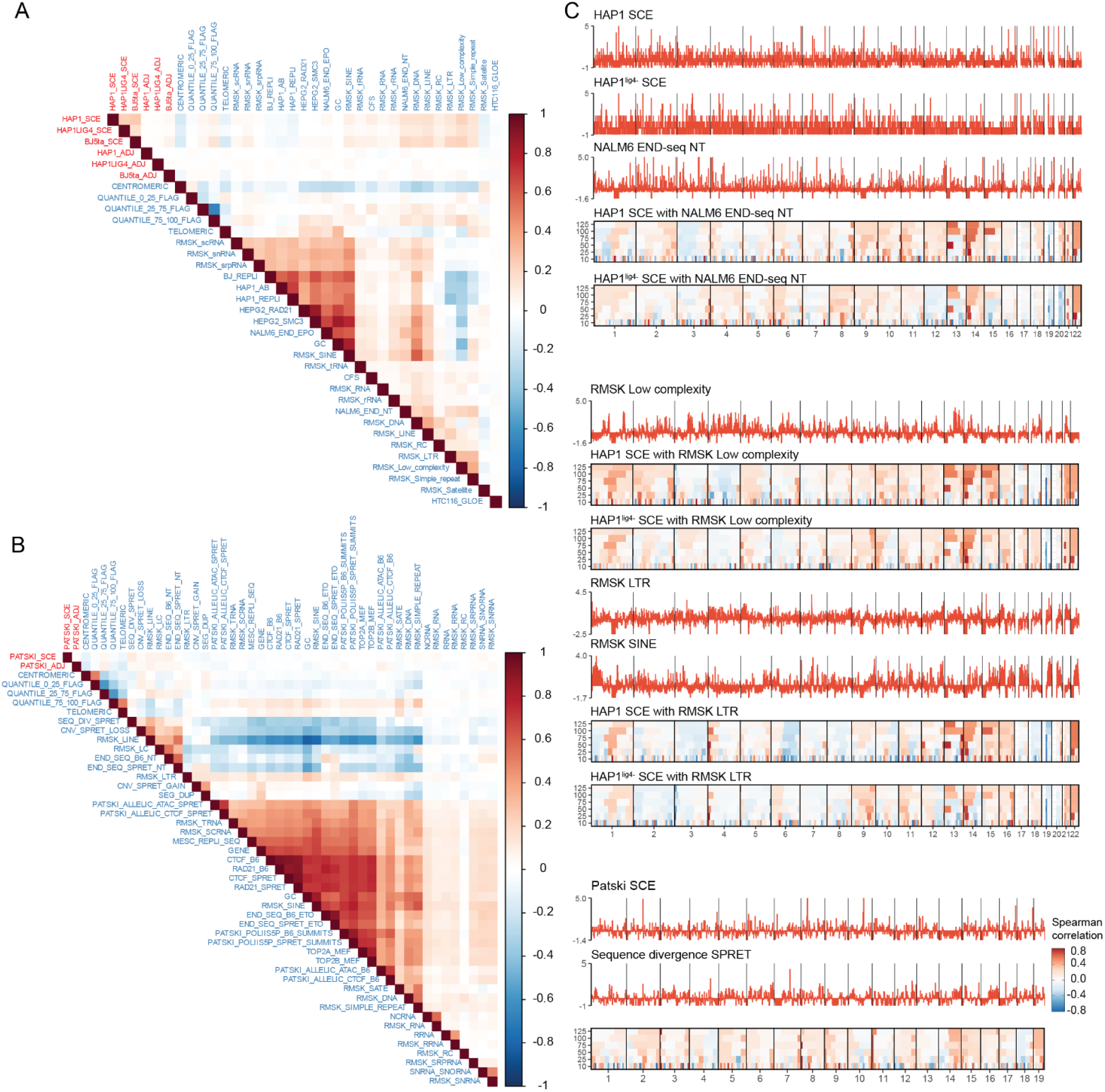
Spearman correlation between genomic features, SCE and mutational events. All pairwise Spearman correlations (positive correlation in red; negative correlation in blue) are shown between various genomic features, SCE pileups, and annotated mutational event breakpoints (ADJ - breakpoint adjacent) calculated using 1Mb sized windows. Cell line specific SCEs and mutational event breakpoints were manually positioned at the top of the matrix and are highlighted in red. The remaining features were sorted using hierarchical clustering. Results of the significance test for each pairwise correlation is provided in TabS5. (A) Correlation of SCEs and mutational events from HAP1, HAP1^lig4-^, and BJ-5ta cell human cell lines and general hg19 features. (B) Correlation of SCEs and mutational events from Patski cell (C57BL/6J x *M. spretus*), general mm10 and *Spretus* strain specific features. (C) Genome-wide distribution of scaled features and SCE correlation domainograms. Domainograms show Spearman correlation between increasing number of 1 Mb bins at windows of 10, 25, 50, 75, 100, 125 Mb. HAP1 and HAP1^lig4-^ cell feature tracks are plotted at 1 Mb resolution, while Patski are plotted at 3 Mb. Top panel shows feature tracks and correlation domainogram between HAP1 and HAP1lig4-SCE and NALM6 END-seq no treatment control (NT). Middle panel shows feature tracks (SCE same as top panel) and correlation domainogram between HAP1 and HAP1lig4-SCE and low complexity and LTR repeats. Correlation domainogram for SINE not shown (similar to LTR). Bottom panel shows feature tracks and correlation domainogram between Patski SCE and sequence divergence (SPRET).

**Figure S11.**
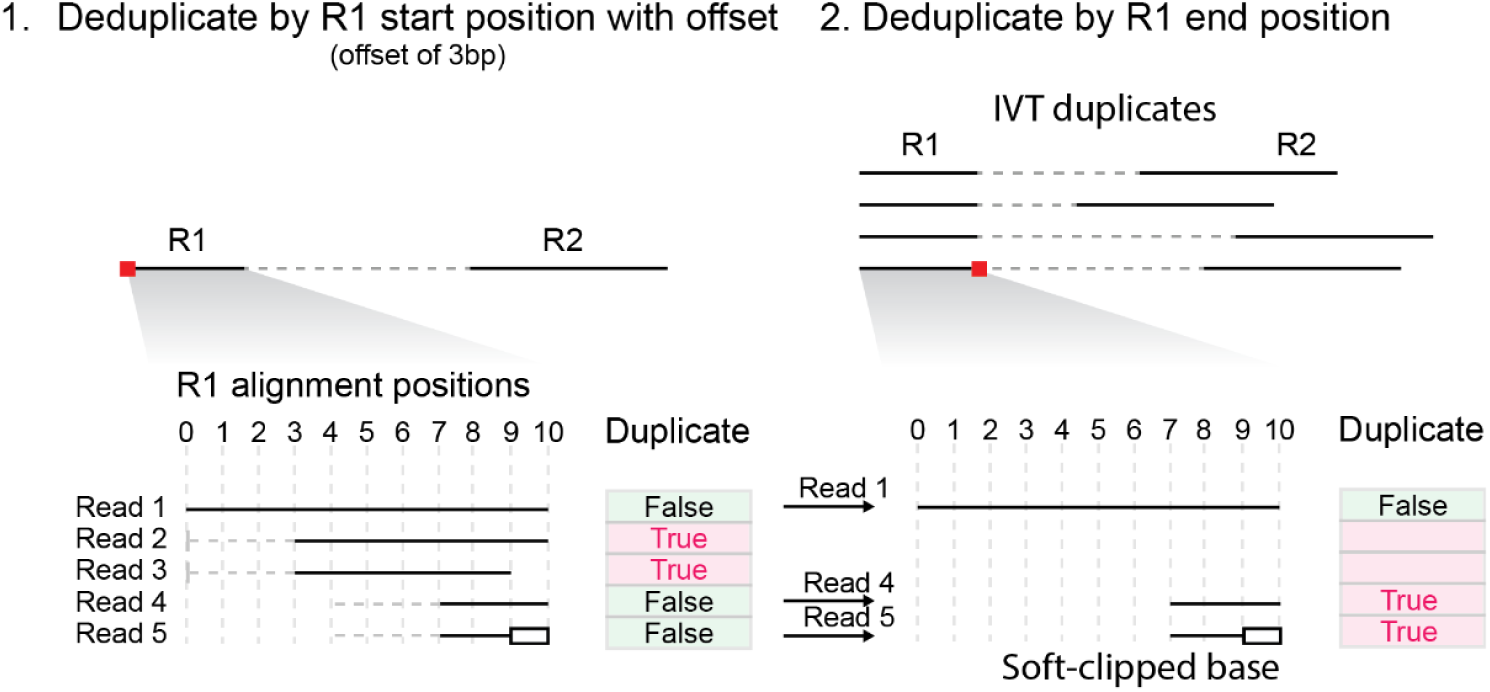
Deduplication scheme used to remove IVT and PCR duplicates. The two-step deduplication first identifies reads with the same R1 start positions, allowing for a 3bp offset upstream of the start position (gray horizontal dashed lines). Next, any soft-clipped bases on the end of R1 are unmasked, followed by the identification of any reads with identical end positions. Reads identified as duplicates from either R1 start and/or R1 end are removed from downstream analysis (see **Methods**). Deduplication for W and C reads is performed separately. Black empty boxes represent soft-clipped bases that were unmasked. In the example diagram, reads 2-5 are considered duplicate reads.

**Figure S12.**
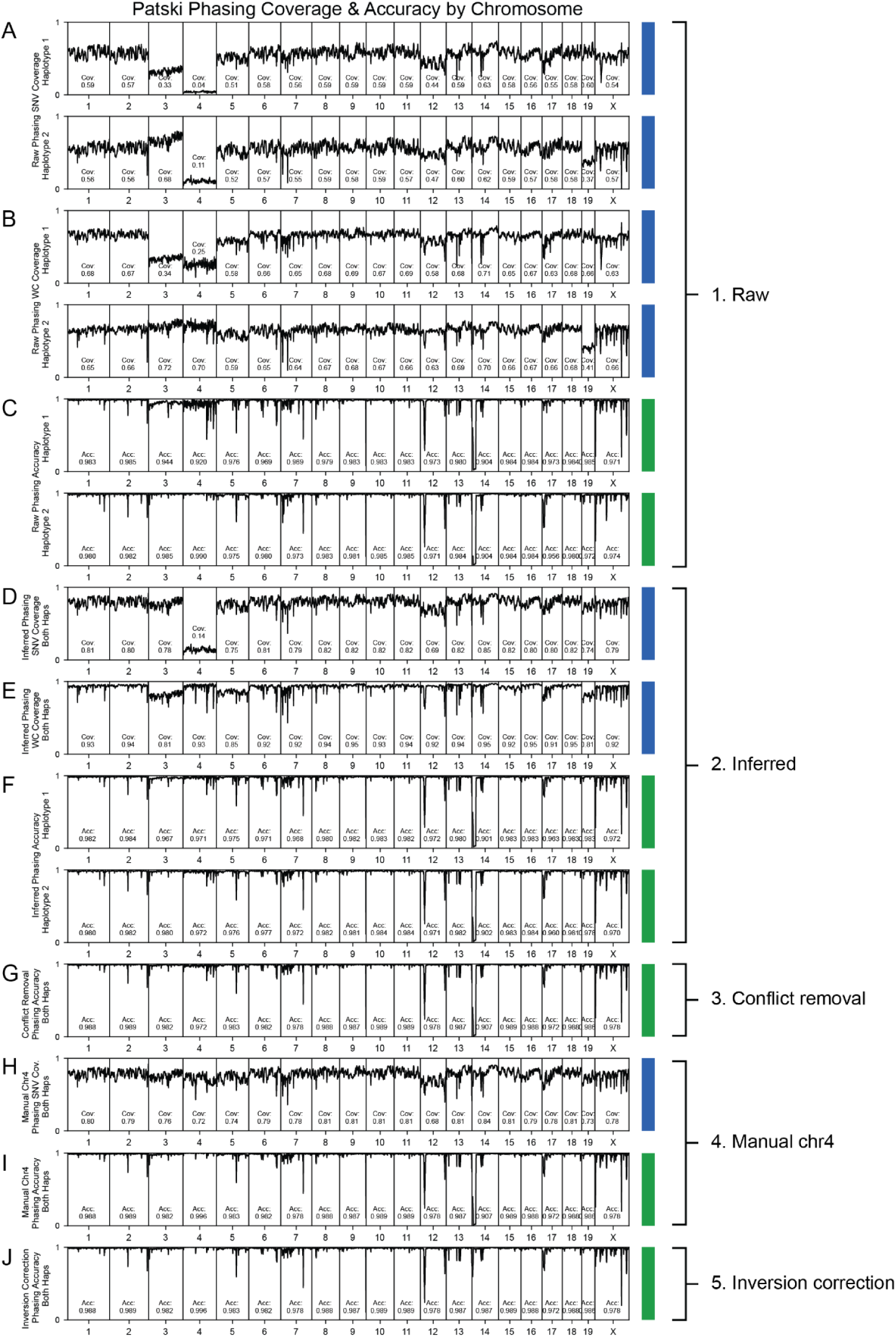
Phasing coverage and accuracy. (A) **Phasing coverages at heterozygous SNV sites with raw StrandPhaseR output for haplotypes 1 and 2.** Each point is the fraction of phased SNVs divided by the total heterozygous SNVs from the Patski bulk VCF file within a 1Mb segment of the genome. Roughly 55% of the heterozygous SNVs (54% hap1 & 56% hap2) are phased by StrandPhaseR (2–4), except chr3 hap1, chr19 hap2, chr4 and chr12. Chromosomes 4 and 12 have many loss events and consequently lower coverage; chr4 phasing used only a subset of 10 heterozygous Patski cells with WC regions to optimize the raw accuracy with respect to phased sites (panel C) which explains the low SNV coverage. Chromosomes 3 and 19 have trisomy events that introduce phasing conflicts on the duplicated haplotype. (B) **Phasing coverages at heterozygous SNV sites with WC coverage using the raw StrandPhaseR output for haplotypes 1 and 2.** Each point is the fraction of phased SNVs divided by the total heterozygous SNVs in the Patski merged VCF with WC coverage within a 1Mb segment of the genome. Roughly 65% of the heterozygous SNVs with WC coverage (63% hap1 & 66% hap2) are phased by default, except chr3, chr4, and chr12 hap1, and chr19 hap2. Larger chromosomes including chr1, chr2, chr3, and chr5 were run on a subset of cells (between 3-500) due to computation memory constraints, so their actual phasing coverage with respect to WC sites could be higher if all samples were used for phasing. Chr4 phasing used only a subset of 10 heterozygous Patski cells with WC regions. Chromosomes 3 and 19 that have whole chromosome gains for haplotypes 1 and 2, respectively. These haplotypes have decreased coverage because cells may have conflicting bases in WC regions. (C) **Phasing accuracies with raw StrandPhaseR output for haplotypes 1 and 2.** Each point is the fraction of accurately phased SNVs divided by the total phased SNVs within a 1Mb segment of the genome. Phasing accuracies are generally above 97% (97.3% hap1 & 97.5% hap2). Lower phasing accuracies are observed for chr3 & chr4 hap1 due to whole chromosome gain and loss events, respectively. Homozygous inversion at the start of chr14 and heterozygous inversions on chr12 both negatively affect phasing accuracy. (D) **Phasing coverage at heterozygous SNV sites after haplotype inference.** Because all SNV’s in this analysis are biallelic, we infer the opposite haplotype for the unassigned SNV. Consequently, phasing coverage increases from 55% to 78% except chr4 and chr12 which have LOH events for a large proportion of cells. (E) **Phasing coverage at heterozygous SNV sites with WC coverage after haplotype inference.** Phasing coverage of WC-covered SNVs increases from 65% to 92% except for chr3, chr5 and chr19 that have trisomies. (F) **Phasing accuracy after haplotype inference for each 1Mb bin of the genome.** Phasing accuracy is above 97% (hap1 97.3% & hap2 97.4%) with a 35% increase in the number of phased sites (22.5 million phased sites for both haplotypes vs 15.5 million for hap1 and 16.1 million for hap2 previously). (G) **Phasing accuracy after haplotype inference and removal of sites with phasing conflicts for each 1Mb genomic bin.** Phasing conflicts are sites where both haplotypes are assigned the same SNV; these are removed due to their ambiguity which improves overall phasing accuracy to 98.1%. (H) **Leveraging clonal LOH on chr4 improves phasing coverage.** Chr4 harbors both monosomy and UPD, unlike chr12 which has predominantly the deletion type of LOH. UPD can contribute to WC coverage and affect both phasing coverage and accuracy. We thus used LOH to directly phase otherwise heterozygous sites. (I) **Phasing accuracy improves with phasing chr4 with LOH.** We observed a 10-fold increase in phased chr4 SNVs with increased phasing accuracy from 97.2% to 99.6%. (J) **Phasing accuracy improves after inversion correction.** Using inversion annotation from sci-L3-Strand-seq, regions of chr14 and chrX had their haplotypes reversed at these positions which increases phasing accuracy by 8% on chr14 but only slightly for chrX as the inversion is small.

**Figure S13.**
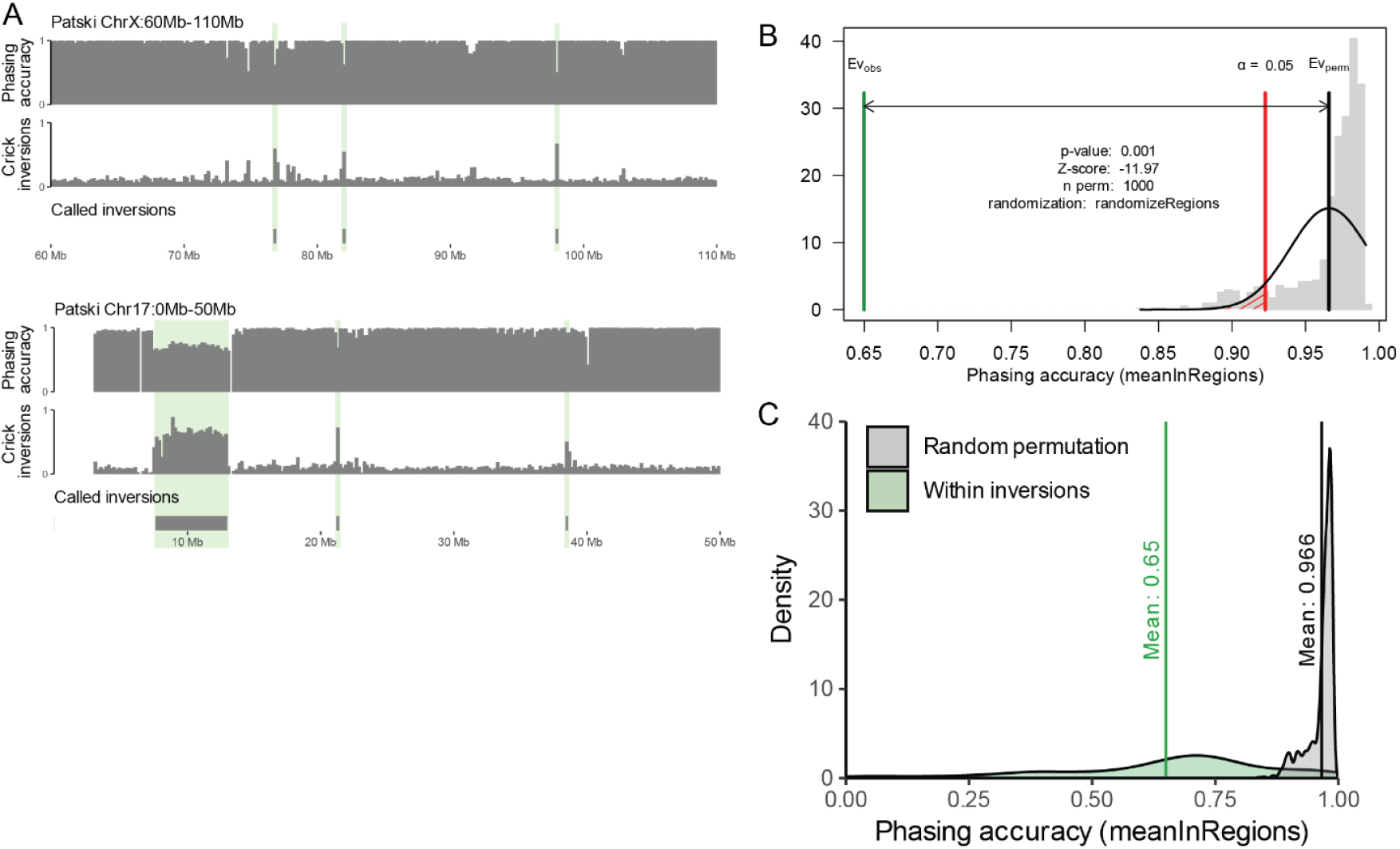
Phasing accuracy decreases over inversions. **(A) Identified inversions overlap with decreased performances in phasing accuracy.** Top panel shows a 50Mb window on chrX, while the bottom panel shows a 50Mb window on chr17 in Patski cells. Accuracy of SNV assignment to each parental strain of origin (C57BL/6J x *M. spretus*) is shown in top tracks of each panel. The phasing accuracy tracks are a zoom-in on the genome-wide final phasing accuracy presented in Fig3A. Middle tracks within each panel show the proportion of cells that contain reads opposite of a region’s strand state, which are indicative of an inversion (Crick reads within a WW strand state region). Bottom tracks within each panel show called inversions. An inversion is called if more than half (0.5) of all cells within a bin have reads with an opposite strand state to the rest of the chromosome. Green highlights show regions of lowered phasing accuracy that overlap a called inversion. There are other regions with a drop in phasing accuracy that may represent inversions that did not meet the call threshold. **(B) Permutation test for the association between inversions and lower phasing accuracy.** Results of the permutation test from the regioneR package (p-value: 0.001) used to assess the difference between the randomly permuted control set (Ev_perm_) and the observed values (Ev_obs_) (13). The bar chart (gray) shows distribution of the mean phasing accuracy (meanInRegions evaluation function) obtained after 1000 random permutations (randomizeRegions function) of inversion coordinates. **(C) Distribution of phasing accuracy within inversions (green) and randomly permuted regions (gray) from A.** Within inversion mean: 0.65, SD: 0.22; random permutation mean: 0.966, SD: 0.026.

**Figure S14.**
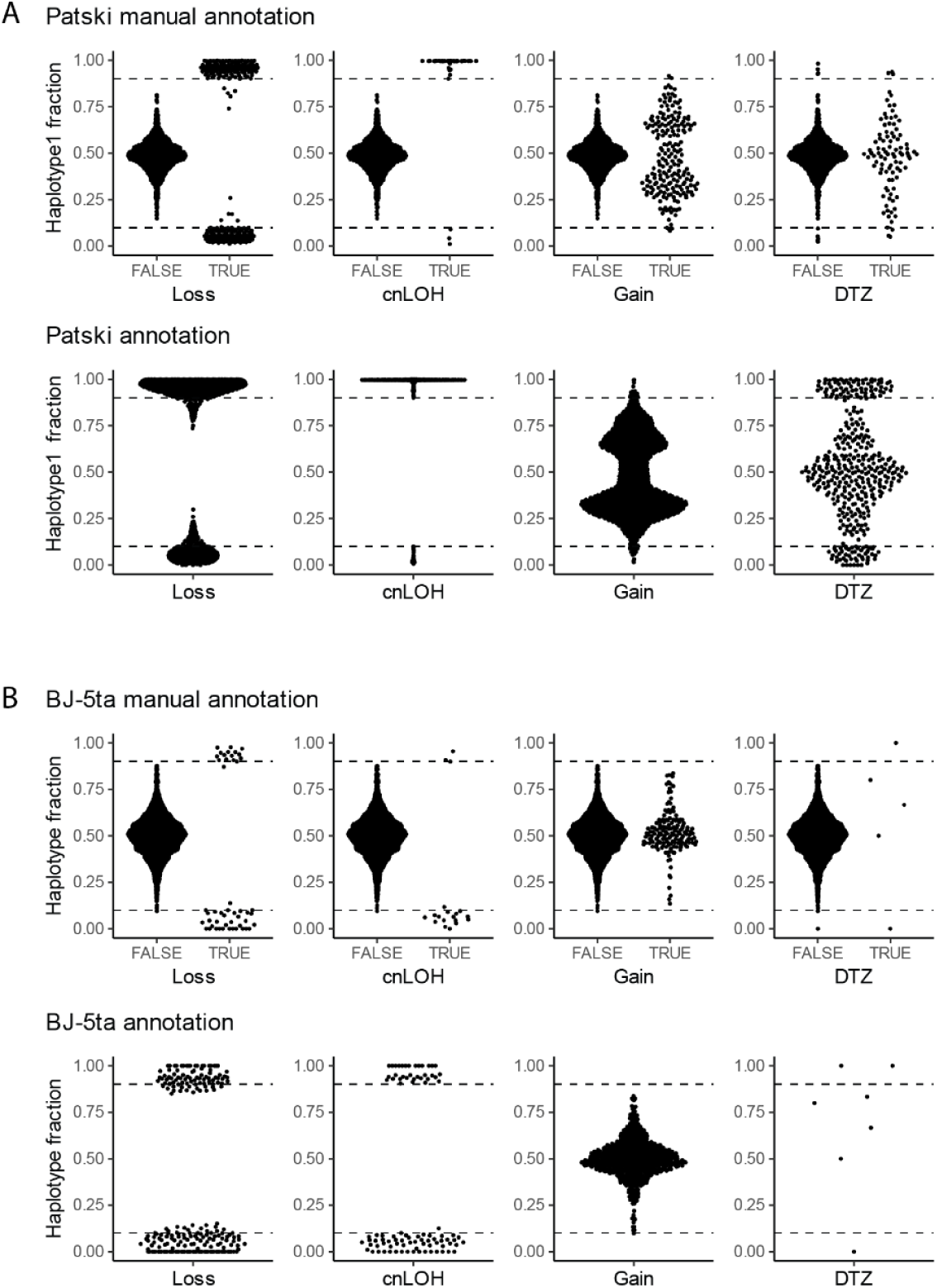
Haplotype fraction distributions within different classes of annotated events. (A and B) The top panels show the distributions of haplotype1 fractions (calculated as haplotype 1/(haplotype 1+ haplotype 2)) in the manually annotated training data. The bottom panels show the haplotype fraction distribution for each SVM annotated event class in all (A) Patski and (B) BJ-5ta cells. The horizontal dashed lines highlight the 0.1 and 0.9 thresholds used for LOH calling.

**Figure S15.**
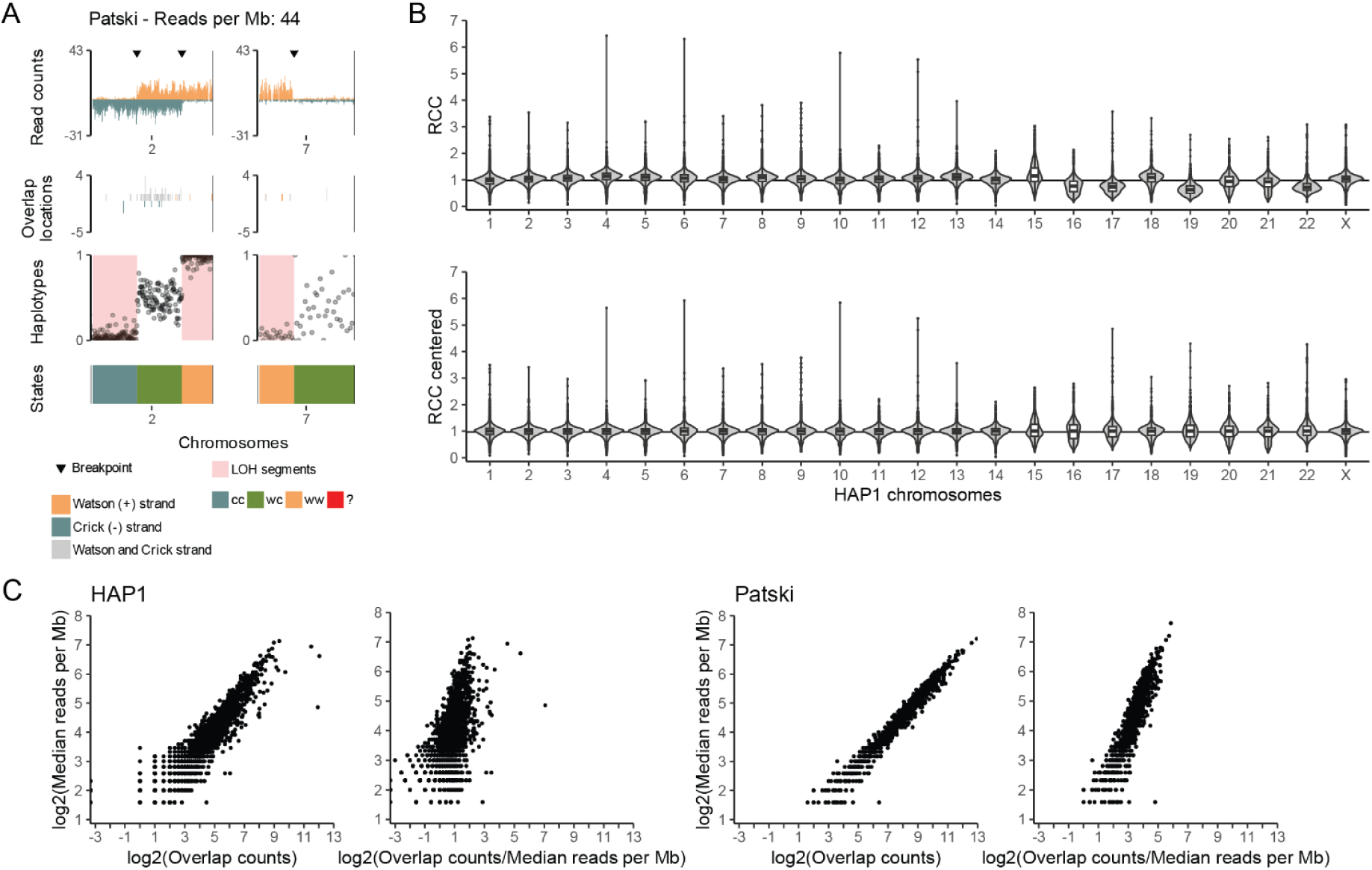
Scores and normalization used for SVM inputs. **(A) Relative chromosome background coverage (RCBC) metric**. Example cell (Patski) showing regions of only background reads (right side of chr7 - drop to zero) that can be distinguished from true signals (chr2 and left side of chr7, regions with LOH) by establishing the level of background within each cell. Cells with higher coverage have higher levels of background, sometimes making regions of drop to zero difficult to distinguish from true signal. The background levels are determined from W or C only regions, such as the LOH regions on chr2. Right side of chr7 consists of only background reads equally represented on W and C strands that match the levels within the LOH regions on chr2. **(B) Relative chromosome coverage (RCC) centering.** RCC is skewed in shorter chromosomes (HAP1 cells shown as example). The median RCC across all cells is expected to be 1 for all chromosomes (black horizontal line). However, some of the shorter chromosomes fluctuate away from 1 (top panel) and require normalization to center the score across all chromosomes (bottom panel). **(C) Relationship between overlapping reads and coverage.** The number of overlapping reads linearly correlates with median reads per Mb (left panel) and was normalized for downstream analysis (right panel). Overlapping reads are from duplicated regions in HAP1 and Patski cells.

**Figure S16.**
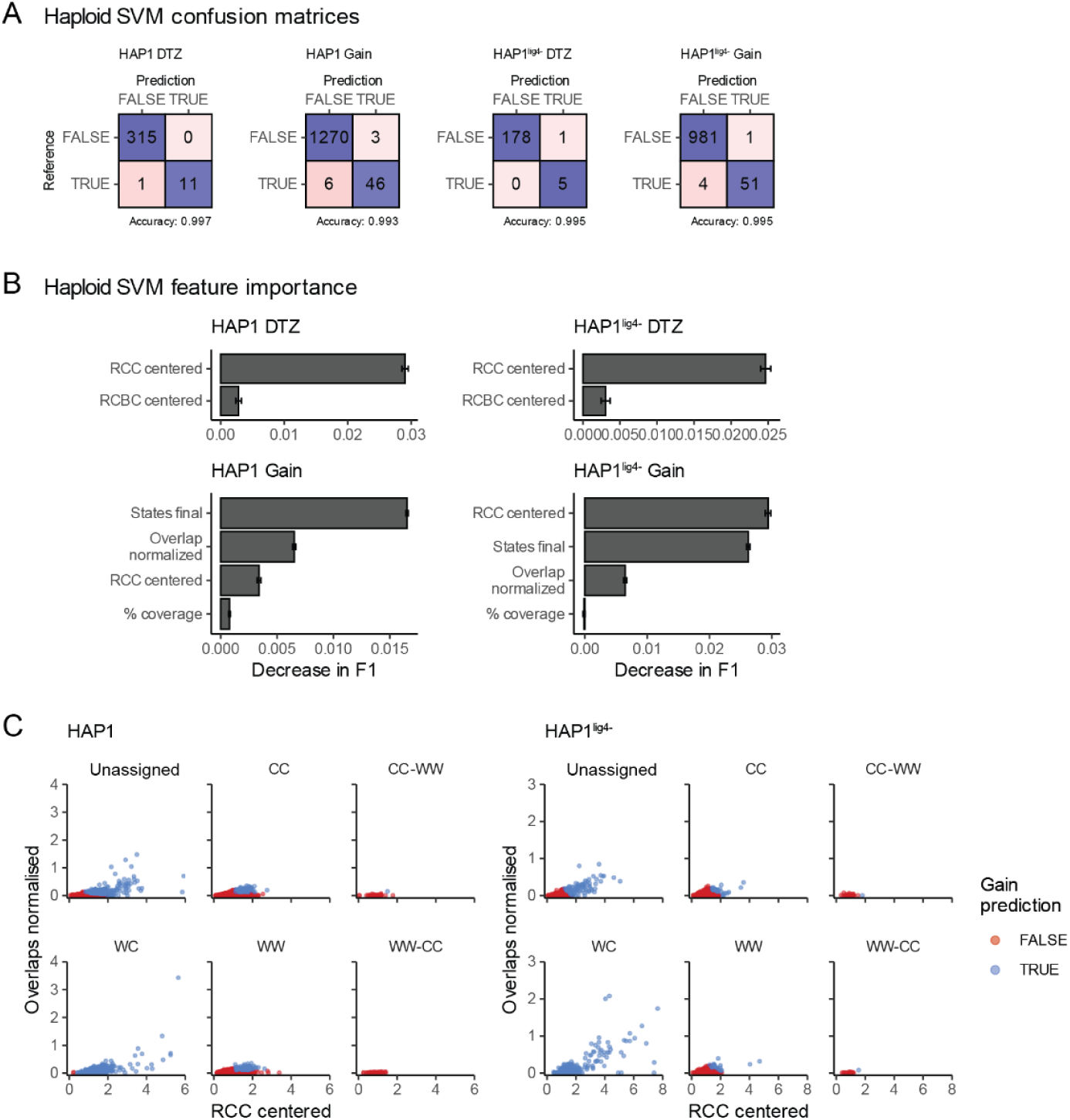
Training evaluation for the haploid SVM event annotation. **(A) Confusion matrices evaluating drop to zero (DTZ) and gain event annotations in HAP1 (right two panels) and HAP1^lig4-^ (left two panels) cells.** True positive and negative events are in purple; false positive and negative events are in pink. **(B) Importance of each feature used in the classification of events in haploid cells.** The feature importance was determined by the decrease of the performance score after 20 random permutations of a feature’s values (**Annotation with Support Vector Machines (SVM) classifiers** in **Methods**). Each bar shows the mean score decrease, with the error bars showing the standard error of the mean (SEM). **(C) Final strand states highlight gain misclassification in haploid cells.** In haploid cells, any region with both Watson and Crick reads must in fact be the results of a gain event. In HAP1 cells (left panel), two segments with a WC state are called as not having a gain. These two segments have a missed breakpoint call and a composite strand state of CC-unassigned, making the original WC state incorrect. We know all CC-WW and WW-CC composite strand state segments are false for gain and so we can confidently correct misclassified segments with these states (**Gain annotation** in **Methods**). Only a single event with CC-WW strand state was mis-classified in HAP1 cells and corrected. In HAP1^lig4-^ cells (right panel), two events with WW-CC and CC-WW strand states were mis-classified as gain and corrected.

**Figure S17.**
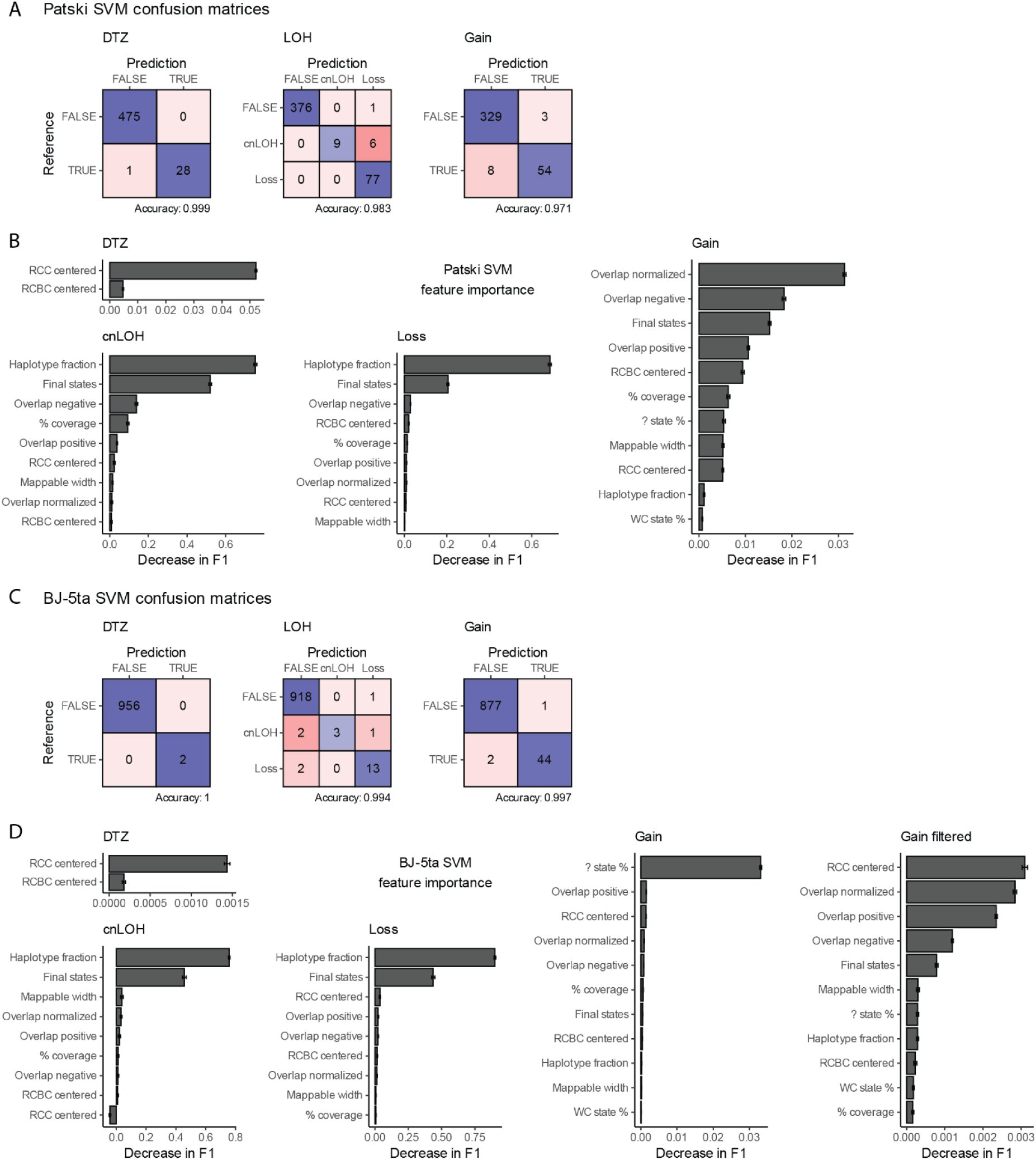
Training evaluation for the diploid SVM event annotation. **(A and C) Confusion matrices evaluating drop to zero (DTZ), loss of heterozygosity (LOH) and gain event annotations in (A) Patski and (C) BJ-5a cells.** True positive and negative events are highlighted with shades of purple, while false positive and negative events are highlighted with shades of red. **(B and D) Importance of each feature used in the classification of events in (B) Patski and (D) BJ-5ta diploid cells.** The feature importance was determined as in FigS16B. The presence of 4n like cells in the BJ-5ta dataset skews the feature importance towards the unassigned state percentage. The rightmost panel for BJ-5ta shows the feature importance without high gain cells, bringing the feature ranks more in-line with Patski cells.

**Figure S18.**
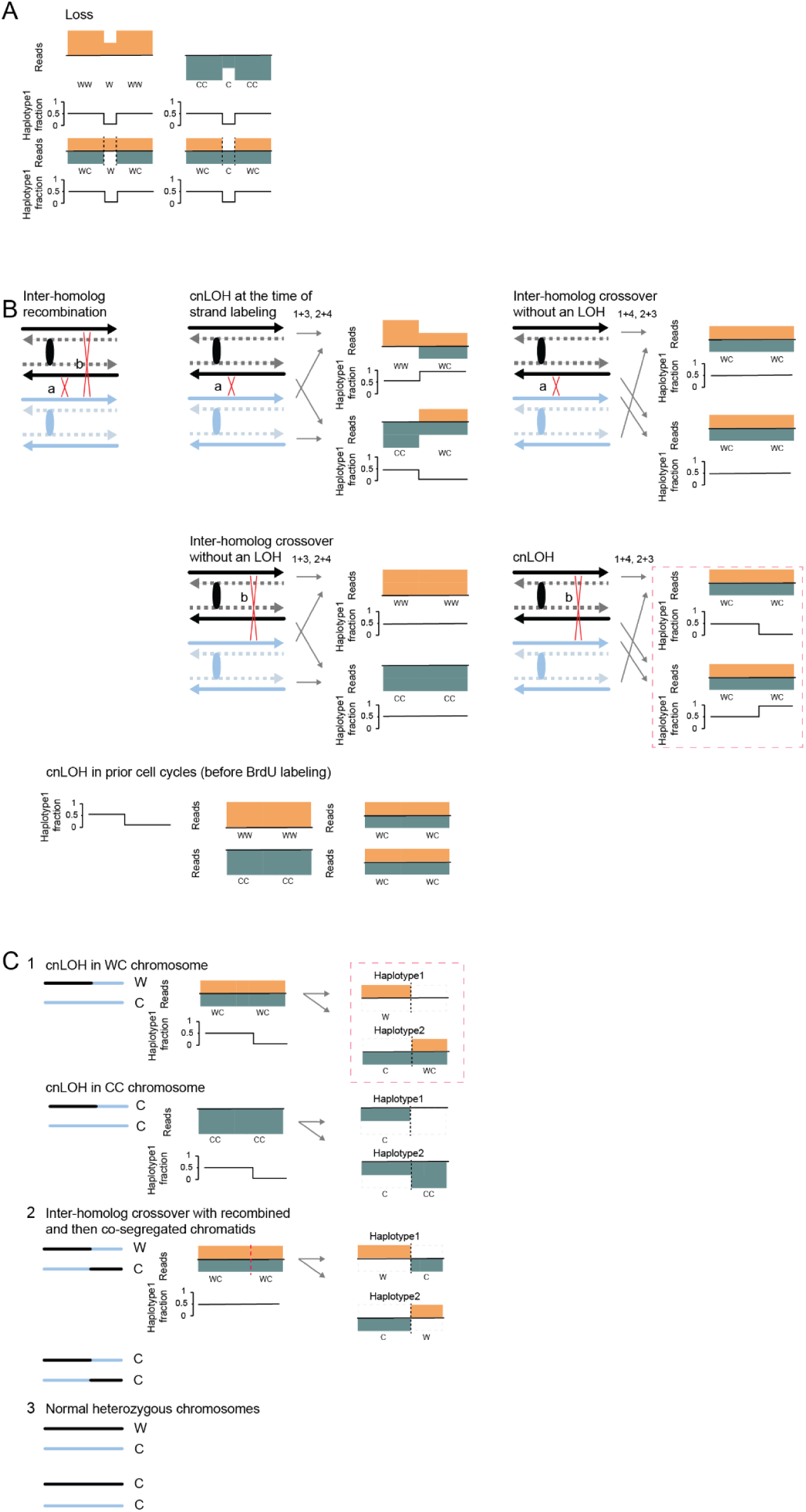
Haplotype-aware segmentation for detection of LOH events. **(A-C)** Vertical dashed lines highlight strand switches. Haplotype1 homologs are shown in black, while Haplotype2 are shown in blue. The homologs each contain two identical sister chromatids. Oval indicates centromere. Solid line depicts the template strand and the dashed line shows the degraded nascent strand. Top strand going left to right is W and the bottom strand going right to left is C. **(A) Strand state inheritance patterns for an interstitial loss.** Loss within regions that have inherited both a W and a C DNA template strand can be identified with a strand switch from WC to W or W to WC. However, low-pass sequencing makes copy number changes harder to identify, which can prevent segmentation and identification of loss in regions that inherited the same orientation template strands (WW or CC). As a result, it is expected that only 50% of loss will be identified from our strand switch-based segmentation. A portion of the other 50% can be rescued from the successful segmentation of other clones containing an identical event. **(B) Inheritance patterns of a terminal cnLOH.** An inter-homolog recombination can happen between templates with (a) opposite or (b) same strand orientations. A cnLOH acquired at the time of strand labeling will contain a strand switch from a recombination between strands of opposite orientation (a - top left panel). A recombination between strands with the same orientation will not contain any strand switches, even if acquired during labeling (b - middle panel). Red dashed box highlights WC chromosomes where reads of each haplotype switched strand orientation if we were to plot strand state for each haplotype. The bottom panel illustrates a cnLOH acquired in the cell cycles prior to strand labeling, showing there are no strand switches despite a haplotype1 fraction change. Haplotype-specific plots would also reveal underlying strand switches. **(C) Haplotype-aware calling of cnLOH events without a strand switch.** The following diagrams are in regards to copy-neutral chromosomes without apparent strand-switches in haplotype-unaware analyses. The top panel (1) shows cnLOH in WC chromosomes (50%) and CC chromosomes (WW chromosomes are similar but not shown here; 50%). Reads containing phased SNVs can be separated by haplotype, creating a C to WC switch for WC chromosomes that allows segmentation in a haplotype-aware manner (highlighted by red dashed box). In the same cell, the reciprocal haplotype containing a W to DTZ switch can further validate the segmentation. Both haplotypes combined improve the segmentation and detection of cnLOH. The C to DTZ and C to CC transitions for CC chromosomes are hard to detect due to low coverage, but as long as the segmentation was successful in WC chromosomes for clonal cnLOH, we can rescue cnLOH segmentation. The middle panel (2) shows a linkage switch without an cnLOH. Haplotype-aware segmentation can also detect inter-homolog crossovers (highlighted by dashed red line) where the recombined chromatids co-segregate into the same daughter cell. The bottom panel (3) shows normal heterozygous chromosomes without inter-homolog recombination. Note that copy-neutral chromosomes that only have SCE will have apparent strand-switch.

**Figure S19:**
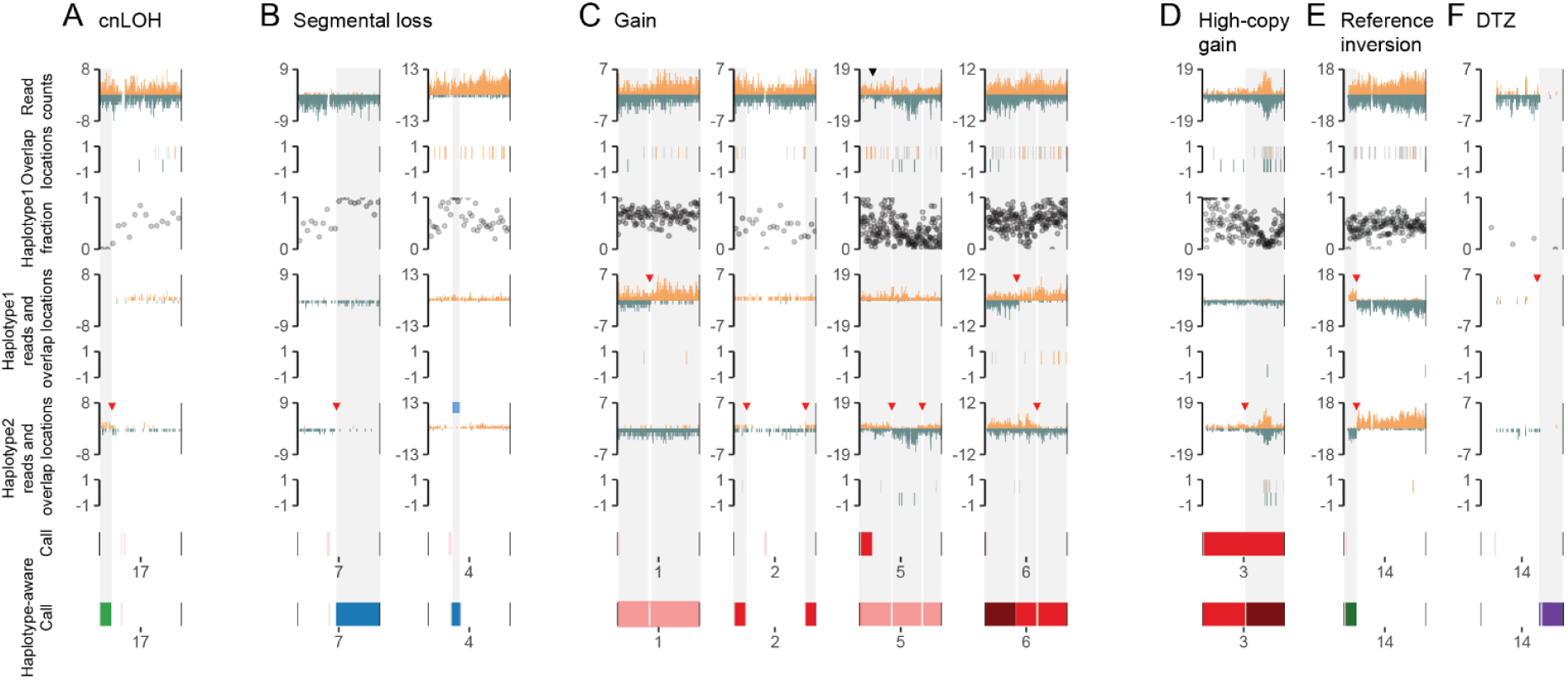
Haplotype-aware segmentation enables the identification and characterisation of additional events. Read counts show W and C reads. Haplotype specific reads are a subset of reads containing phased SNV(s) belonging to a single haplotype. Overlapping reads show the locations of overlaps. Each overlap location is either dark green, representing two overlapping C strand reads, dark yellow, representing two overlapping W strand reads, or gray representing two overlapping reads of opposite strands. Each point on the haplotype1 fraction track represents the phased allelic ratio of approximately 50 binned SNVs for Patski and 15 for BJ-5ta. Red vertical lines within the call tracks represent the centromere. Segments highlighted in red within the call track represent gain annotations. Strand switch-based segmentation is highlighted by black triangles in the read count tracks. Haplotype-aware segmentation is highlighted by red triangles. Gray highlights show new annotations as a result of the haplotype-aware segmentations and manual curation. Breakdown of original, haplotype-aware, and manually curated haplotype-aware annotations is provided in TabS11. **(A) Haplotype-aware segmentation helps to call cnLOH**. The uniform read counts on both strands of chr17 in the BJ-5ta cell resulted in no initial strand switch calls, despite a drop in the haplotype1 fraction. Breakpoint calling on each haplotype individually (haplotype-aware) produced segmentation for both haplotypes (red triangle), highlighting a region with no reads in haplotype1 and WC reads in haplotype2 (highlighted in gray, green highlight cnLOH on haplotype-aware call track). **(B) Haplotype-aware segmentation helps to call segmental loss.** Drop in read counts from a loss can be difficult to discern, especially for regions with WW or CC strand states lacking an identifiable strand switch. These losses can be clearly observed in haplotype specific read tracks (gray highlight, blue highlight on haplotype-aware call track). On chr7 (left panel) and chr4 (right panel) in BJ-5ta cells, the DTZ (absence of reads) is present within haplotype2 for both cells and enables segmentation of these regions (red triangle). The terminal loss on chr7 despite showing a haplotype1 fraction drop was just below the cutoff threshold due to segmentation by whole arms and the inclusion of a centromere proximal heterozygous region. The chr7 loss is not a clonal event and therefore is not rescued using segmentation from other cells. **(C) Segmental and whole chromosome gain.** ***New haplotype-aware segmentation helps to call trisomy with SCE.*** Similar to loss, subtle changes in read counts can be difficult to recognize. The first panel shows chr1 from a Patski cell with a haplotype1 trisomy containing an SCE (red triangle; light red highlight on haplotype-aware call track). The presence of an unidentified SCE masked the presence of the trisomy in our initial haplotype-unaware analysis.The first segment (left of the red triangle) contains both W and C strand reads for haplotype1 highlighting the presence of three copies. ***New haplotype-aware segmentation helps to call segmental gain.*** For the second segment (right of the red triangle), the gain can be identified based on the same strand overlapping locations, which get diluted without the correct segmentation. The second panel shows an example of terminal segmental gains on chr2 in a BJ-5ta cell (gray highlight, red highlight on haplotype-aware call track). Similar to the first panel, the regions with an additional copy for haplotype2 appear as regions with WC strand states, which once segmented also contain W same-strand overlapping reads. ***Haplotype-aware overlapping reads provide digital counts of haplotype specific copy numbers.*** The third panel shows a trisomy on chr5 in Patski cells containing two SCEs (red triangles). Initial analysis found a gain within the centromere proximal segment (red highlight on call track), but similar to the first panel example, the presence of the gain in the remaining segment failed to be annotated due to the masking effect of the unidentified SCEs (light red highlight on haplotype-aware call track). Haplotype2 specific read counts and overlapping locations reveal a CC state segment with two copies that is preceded and followed by WC state segments. ***New haplotype-aware segmentation helps distinguish 3- from 4-copy number gains.*** The fourth panel shows chr6 from a Patski cell containing a 4-copy gain followed by regions with 3-copies (gray highlights, dark red and light red highlights on the haplotype-aware call track). The initial 4-copies appear as WC regions in both haplotype1 and haplotype2. A switch to W only reads for haplotype1 with a lack of any overlapping reads is indicative of a loss, suggesting a haplotype2 biased (still WC) 3-copy number that is also reflected in the haplotype1 fraction track. The subsequent switch to C only reads for haplotype2 with multiple W same strand overlapping reads in haplotype2 indicates a haplotype1 biased 3-copy number that once again is reflected in the haplotype1 fraction track (second gray highlight). **(D) New haplotype-aware segmentation highlights high-copy number gain.** The high-copy number segmental gain on chr3 in a Patski cell resulted in a whole chromosome gain annotation in our initial analysis due to the information being averaged across an entire chromosome. Haplotype2 specific overlapping reads and haplotype-aware segmentation highlighted an amplification of only a small interstitial region (gray highlight, dark red highlight on the haplotype-aware call track). **(E) New haplotype-aware segmentation highlights mouse reference inversions.** Haplotype-aware analysis reveals inversion within the reference genome as reciprocal SCE (red triangle), while the haplotype-unaware read counts appear as a uniform whole chromosome WC strand state (dark green highlight on the haplotype-aware call track). **(F) New haplotype-aware segmentation helps to call segmental DTZ.** BreakpointR for calling strand-switches was not designed to identify regions with only background reads, meaning some DTZ regions without sufficient background reads will not be segmented. The lack of segmentation will prevent the annotation of DTZ. For haplotype-aware segmentation tried overcoming this limitation by adding background noise to each cell to force breakpointR into recognizing DTZ regions as regions with a WC strand state. Additionally, we performed a bin based SVM annotation of DTZ based on HAP1 cells training data. These two approaches enabled us to not only identify haplotype-specific loss, but also identify segmental DTZs that were previously missed (gray highlight, purple highlight on the haplotype-aware call track).

**Figure S20.**
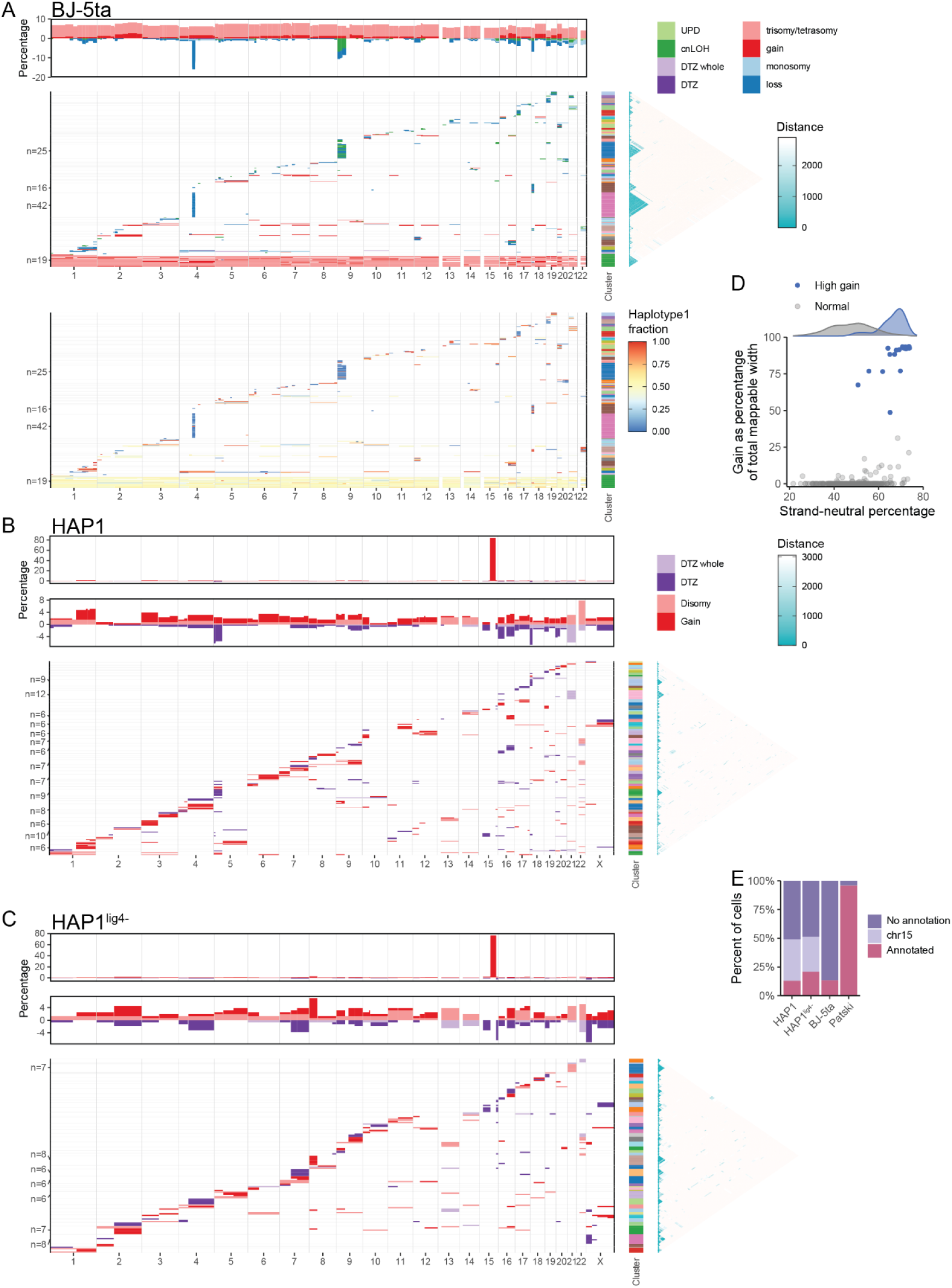
Heatmaps of mutational crossover outcomes. **(A-C) Heatmaps of mutational crossover outcomes in (A) BJ-5ta, (B) HAP1 and (C) HAP^lig4-^ cells.** The heatmaps only show cells with annotated events. The top panel of each plot shows a summary of all the annotated events as the percentages of cells with an annotation at a given genomic location. For HAP1 and HAP1^lig4-^ cells, an additional panel shows the percentage after removing annotation of the disomic region on chr15. Each row of the main plots is a single cell ordered into clusters using Hierarchical clustering. The different clusters are shown on the right side of each main plot, with (A) 58 clusters for BJ-5ta, (B) 60 clusters for HAP1, and (C) 48 clusters for HAP1^lig4-^. Number of clusters was chosen using the silhouette method. The Canberra distance matrix with 1Mb bins shows the pairwise similarity between cells and was used for hierarchical clustering (white-less similar; blue - more similar). Each cluster and the cells within were ordered by the start position of the most upstream annotated event. The plotted HAP1^lig4-^ annotation was obtained with an SVM model trained using the HAP1 manual annotation (TabS12). For BJ-5ta cells, an additional heatmap with the haplotype1 fraction overlaid for each annotated event is shown under the main event annotation heatmap. **(D) Distribution of strand-neutral percentages for normal and high gain BJ-5ta cells.** Regions that have a WC strand state and/or not biased enough to be assigned to WW or CC states are defined as “strand-neutral”. High gain BJ-5ta cells display slightly higher strand-neutral percentages but are not removed as unsuccessful Strand-seq libraries. **(E) Percent of cells with annotated events.** For HAP1 and HAP1^lig4-^ cells, the known disomic region on chr15 was excluded from the main set of events. Cells with only the chr15 disomic region annotation are plotted as a separate category (chr15).

**Figure S21:**
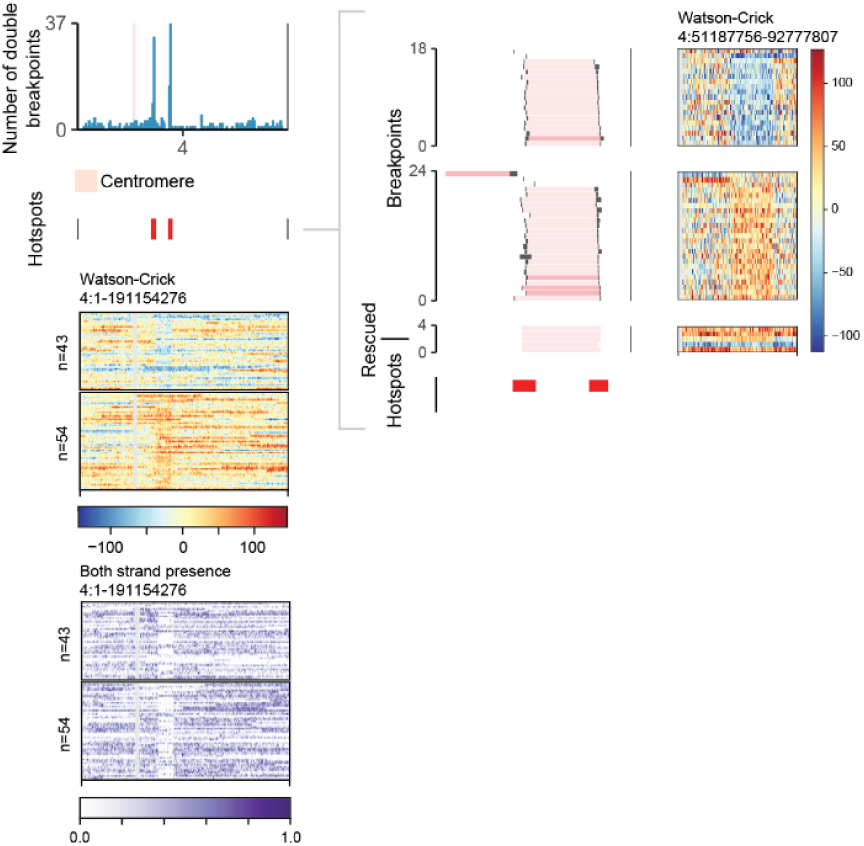
Identifying clonal LOH from double breakpoint hotspots. The sub-clonal interstitial loss measuring between 10.9-19.5Mb was observed on chr4 in BJ-5ta cells as a hotspot (red) of coincident breakpoints shown in the top left bar plot. Below, matching whole chromosome heatmaps show cells with two or more coincident breakpoints, with one cell plotted per row at an approximate resolution of 200Kb. Bins in the top heatmap show the subtraction of W from C read counts, with W reads plotted in red and C reads plotted in blue. Masked regions, namely the centromere, are plotted in gray. The bottom heatmap shows regions where both W and C reads are present in purple, highlighting the clonal deletion as a block of only W or only C reads. A zoom-in on the breakpoints overlapping the two hotspots is shown in the right panel along with a matching zoom-in heatmap showing the W minus C read counts. Breakpoints are shown in black, while SVM loss annotations (details in **Annotation of mutational mitotic crossover outcomes reveals clonal populations**) are plotted in light red, while manual annotations are in dark red. The SVM annotation identified the clonal loss segment for 89.7% (35/39) of cells with double strand switches (light red). The remaining 10.3% (4/39) of cells with double strand switches identified from the hotspots were manually inspected and annotated as loss (dark red). LOH rescue of cells without strand switches further identified 5 additional cells.

**Figure S22.**
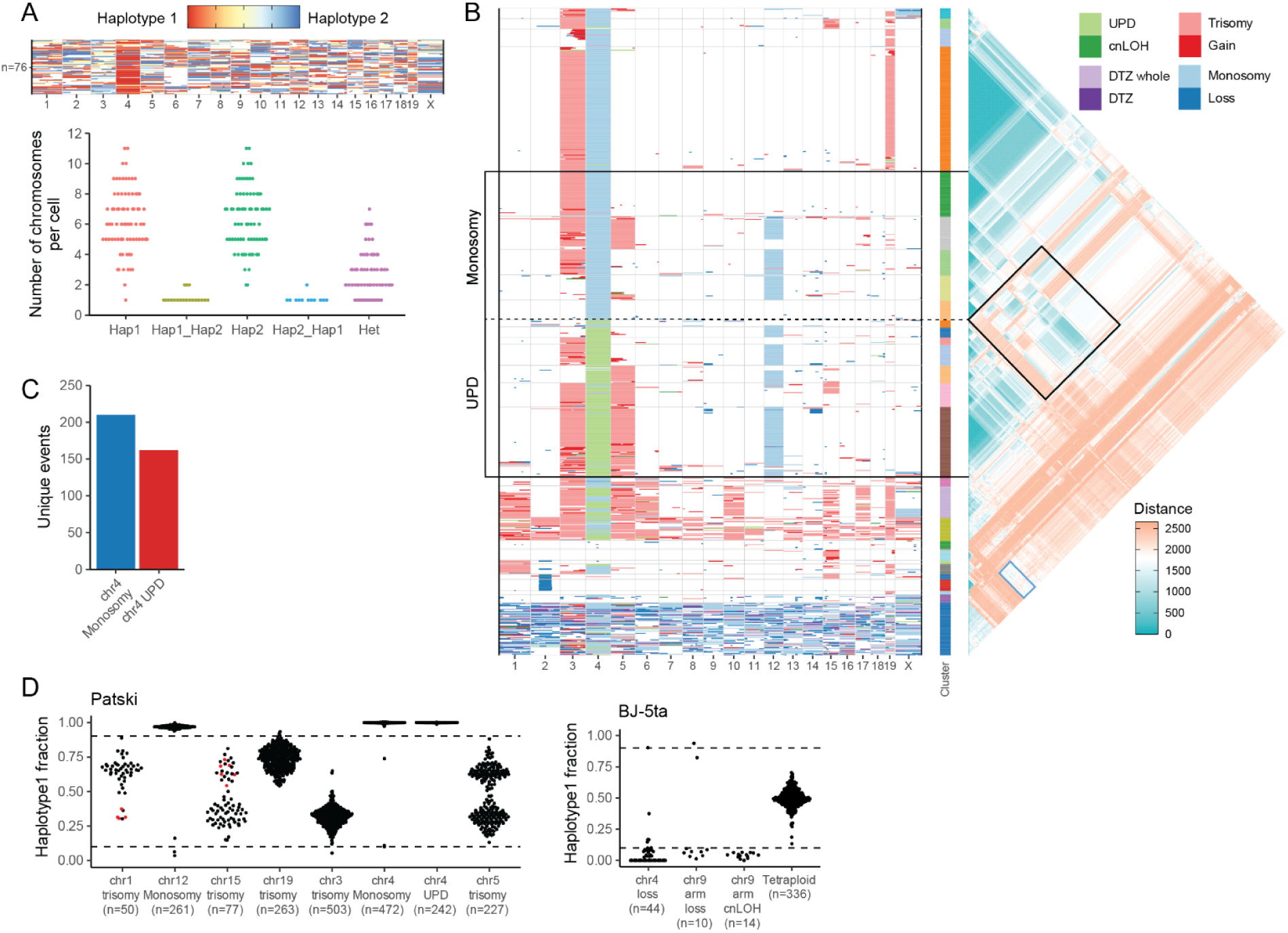
Examination of mutational events and haplotype selection. **(A) Low levels of haplotype switching within cells with a high number of loss events.** Top panel shows the cluster of high loss cells overlaid with the haplotype1 fraction, where haplotype1 is red and haplotype2 is blue. The bottom panel shows the number of chromosomes per cell with a single haplotype or heterozygous (Hap1, Hap2, or Het), or a switch in haplotype (Hap1 to Hap2, Hap2 to Hap1). A haplotype1 fraction of <= 0.2 and >= 0.8 was used to categorize Hap1 and Hap2 states respectively, while anything in between was categorized as Het. **(B) Heatmap of mutational crossover outcomes highlighting mirror clonal populations separated by chr4 monosomy and UPD.** Each row in the main panel shows the annotations of a single cell. The heatmap only shows cells with annotated events. The Canberra distance metric showing pairwise similarity of 1Mb bins between cells (red – less similar, blue – more similar) was used for hierarchical clustering and is presented on the right side of the main plot. Different category weights were chosen to group cells by chr4 Monosomy and UPD (highlighted region in black with dashed line). Mirror populations created by monosomy and UPD on chr4 are highlighted in black on the distance heatmap. Similarity between cells with many loss events and cells with many gain events is highlighted in blue on the distance heatmap. **(C) Chr4 monosomy has higher diversity of unique events compared to UPD.** Cells with identical events were deduplicated based on the 1 Mb weighted count matrix. Excluding high gain and high loss cells, the remaining cells were tallied for unique events based on chromosome 4 monosomy or UPD. **(D) Haplotype bias of events in Patski and BJ-5ta cells**. The distributions of haplotype1 fractions (calculated as haplotype 1/(haplotype 1+ haplotype 2)) for sets of whole chromosome and segmental annotated events. The horizontal dashed lines highlight the 0.1 and 0.9 thresholds used for LOH calling. For chr1 and chr15 trisomy, the red points highlight Patski cells with chr2 segmental loss, which predominantly has a different haplotype gain.

**Figure S23.**
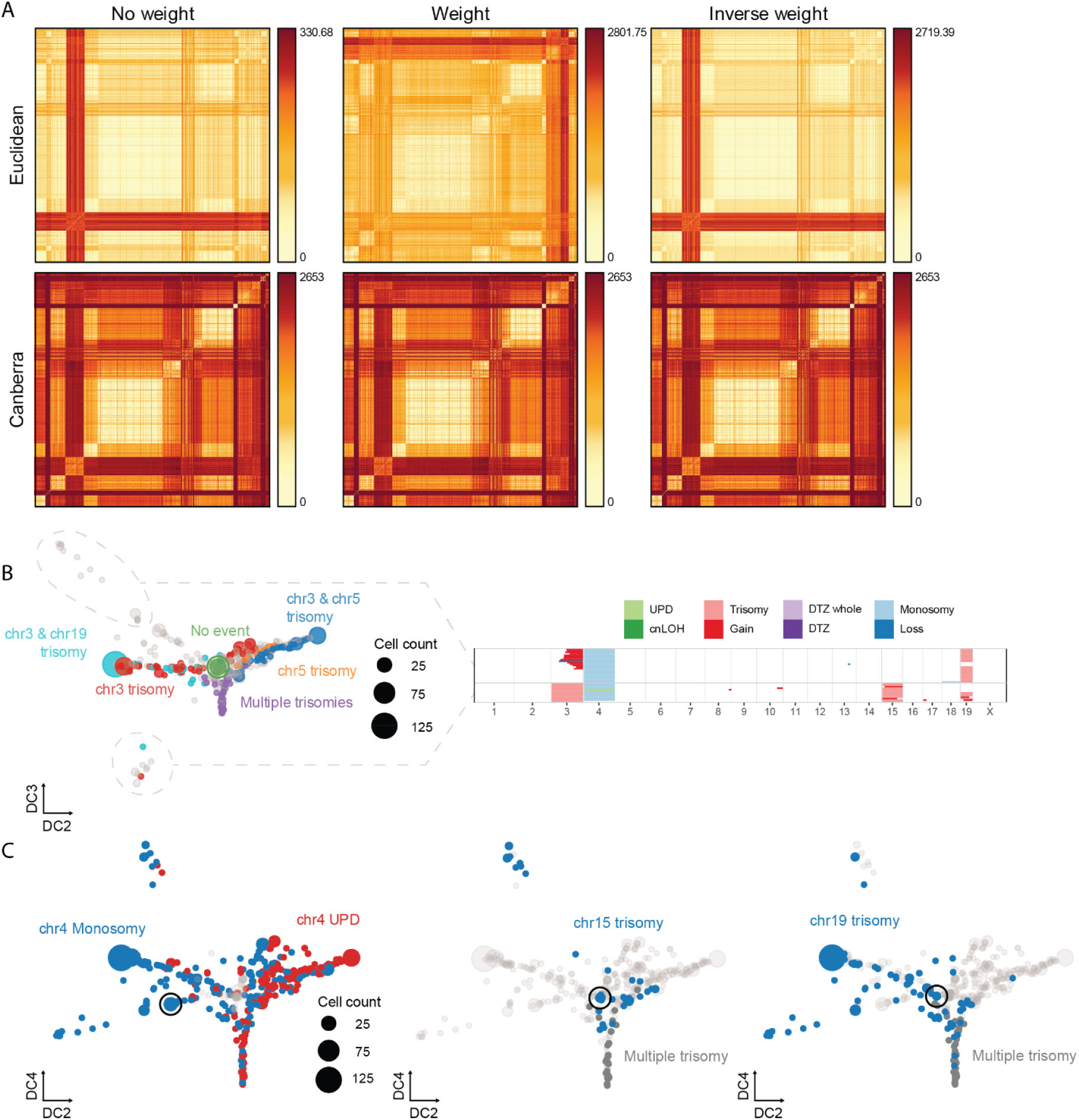
Diffusion map construction and annotation. **(A) Canberra distance is tolerant of different count weights.** Each heatmap shows the Euclidean (top row) or Canberra (bottom row) distance between single cells with a unique set of annotated events at a 1Mb resolution. Distances were calculated from count matrices without weights (left panel), with weights (middle panel) and with inverted weights (right panel). **(B-C)** High gain and high loss cells were excluded prior to making the diffusion maps. Multiple trisomy represents cells with trisomy on chromosome 1, 3, 5, 6, 10, 15, 17 and/or 19. Each point represents a unique set of annotated events, while the size of each point represents the number of cells that all share the same set of annotated events. **(B) Diffusion map with heatmap highlighting events that drive diffusion component 3 (DC3).** Despite somewhat separating cells with multiple trisomies, DC3 is mainly driven by two clusters. The first cluster consists of chr3 gain annotations, while the second is a combination of chr15 trisomy/gain and chr19 trisomy/gain. With DC3 being driven by a narrow set of cells, we chose to instead use DC4 that better separates out the diverse trisomy events. **(C) Diffusion maps annotated with select mutational crossover outcomes.** All panels show diffusion components 2 (DC2) and 4 (DC4). Circled points show cells with a single event that likely represents the founder population from which other clonal populations originated. For the first panel, the chr4 monosomy founder is highlighted. The second panel highlights the chr15 trisomy founder, while the third panel highlights the chr19 trisomy founder.

**Figure S24.**
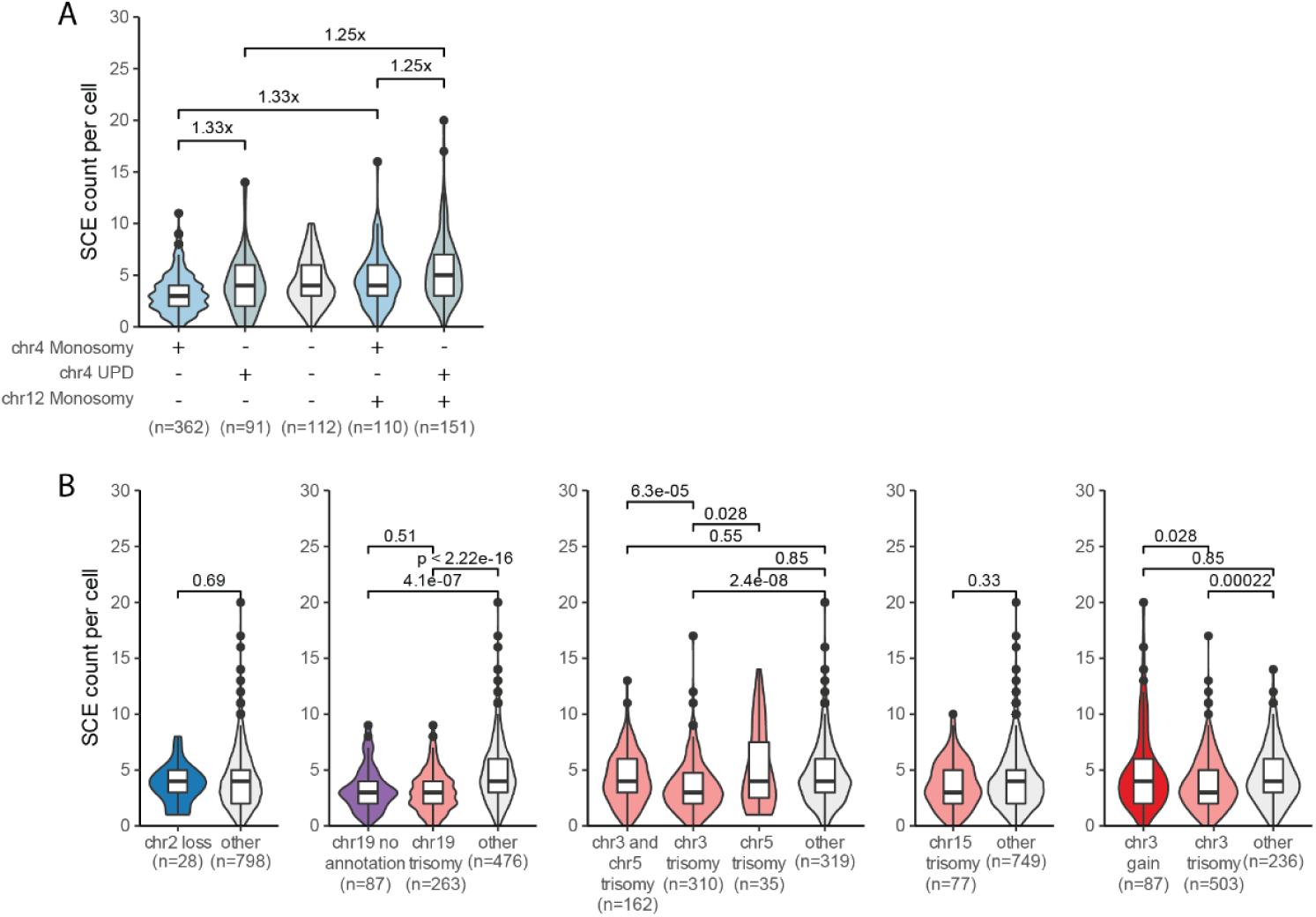
Influence of mutational crossover outcomes on the rate of SCE. **(A) Additive effect of chr4 UPD and chr12 monosomy on rate of SCE.** For pairs of annotated events the fold change in the median SCE counts per cell are shown above the relevant plots. Significant differences were calculated with Mann-Whitney U for chr4 monosomy without chr12 monosomy vs chr4 UPD without chr12 monosomy (p-value: 8e-03); chr4 monosomy without chr12 monosomy vs chr4 monosomy with chr12 monosomy (p-value: 6.45e-07); chr4 monosomy with chr12 monosomy vs chr4 UPD with chr12 monosomy (p-value: 6e-03); chr4 UPD without chr12 monosomy vs chr4 UPD with chr12 monosomy (p-value: 1.14e-04). Additional summary statistics such as median and MAD are provided in TabS13. **(B) Segmental loss, gain and trisomy does not influence the rate of SCE.** The significant differences calculated with Mann-Whitney U are shown above the relevant combination of plots. The “other” category contains all cells not present within the main categories. For the second panel, the “chr19 no annotation” category is limited to cells with chr3 trisomy and chr4 monosomy, such that they match the cells with chr19 trisomy in all aspects except chr19 itself. The “chr3 trisomy” category in the third panel excludes any chr3 trisomies with a coincident gain or trisomy annotation on chr5, while the “chr5 trisomy” category excludes any chr5 trisomies with a coincident gain or trisomy on chr3.

**Figure S25.**
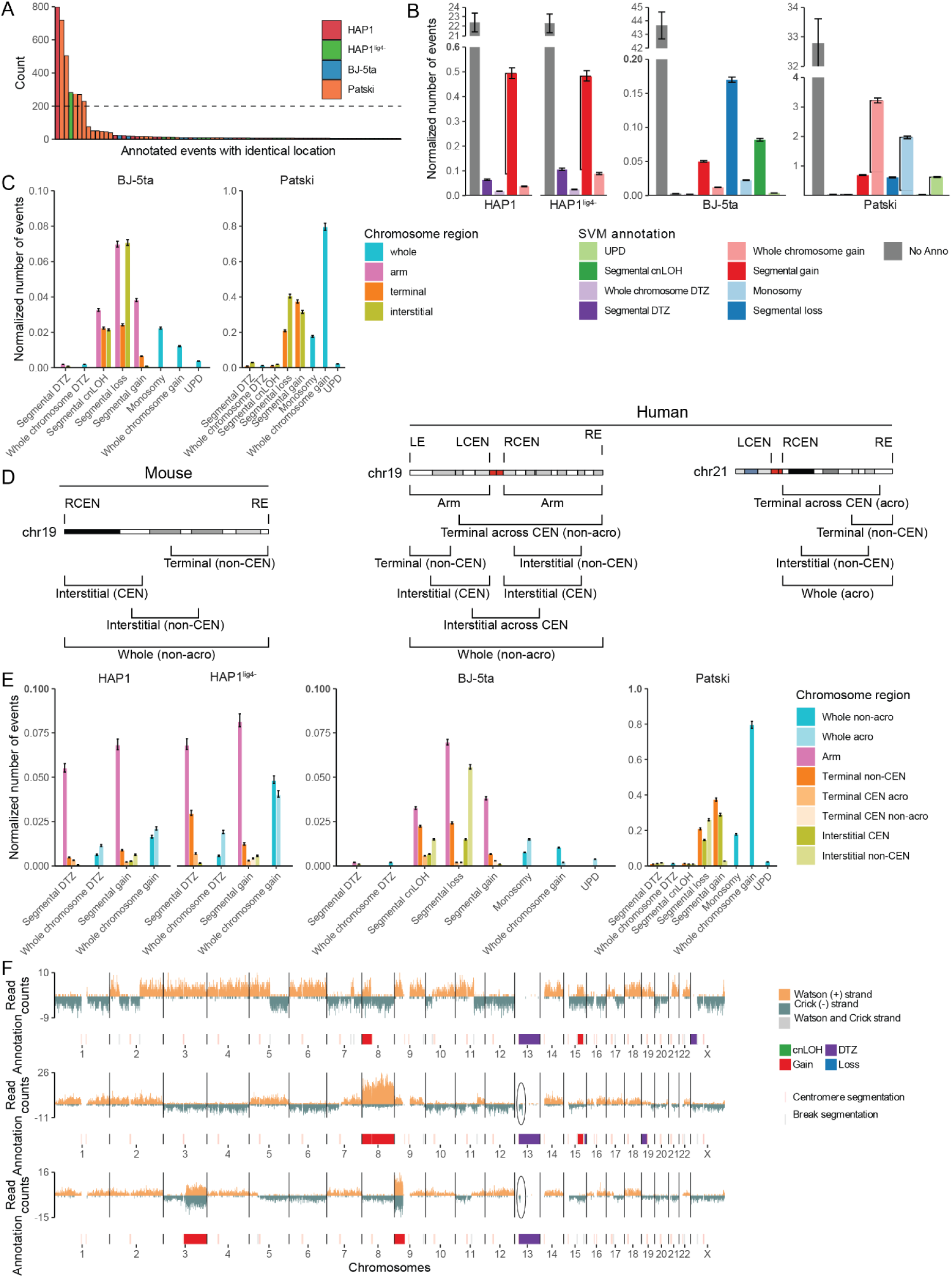
Classification and quantification of annotated events. **(B, C, E)** Event counts were normalized by cell line ploidy and total number of cells. Error bars show the 95% confidence interval. **(A) Identifying annotations arising from clonal expansions.** Annotated events within each cell line were grouped together by identical start and end positions and counted. A set of events with counts above 200 (dashed line) were labeled as pre-existing events. An example of pre-existing events are the disomic region on chr15 in HAP1 and HAP1^lig4-^ cells (red and green bars respectively). The threshold was chosen based on the sharp drop-off observed after the first 7 events and was meant to strike a balance between the removal of dominant pre-existing events while preserving recurring events. **(B) Normalized counts of all mutational mitotic crossover events by chromosome.** Black lines parallel to the event bars mark pre-existing events from (B, e.g. chr15 segmental gain). For each cell, every chromosome was counted as either having no annotation (No Anno) or having one of the eight annotation categories. For BJ-5ta cells the sex chromosomes are excluded. **(C) Normalized counts of mutational SVs in BJ-5ta and Patski cells split by chromosome location.** Contingency tables with normalized counts, fold change and chi-squared p-values are provided in TabS14. **(D) Location features of mutational events.** Diagram for a human metacentric (chr19), acrocentric (chr21) and a mouse telocentric (chr19) chromosome depicting the location features used for the classification of mutational mitotic crossover events. Events overlapping a combination of different location features such as the right side of the centromere (RCEN) and the right end of the chromosome (RE) would be labeled with one of the seven classifications. In the example of RCEN and RE, the annotation would be labeled as an arm event. **(E) Normalized counts of mutational SVs split by chromosome location, centromere and chromosome type.** Pre-existing events and chromosomes without any event annotations are not plotted. Whole chromosome events on acro-centric chromosomes were split into a separate category (whole acro). Interstitial events were split depending on whether they contact the centromere (interstitial CEN, almost exclusively in Patski) or do not (interstitial non-CEN). Terminal events not spanning the whole arm were additionally split by whether they go across the centromere. Terminal across CEN is hardly observed, regardless of whether they are on an acro-centric chromosome (terminal CEN acro or terminal CEN non-acro). Most terminal events involving CEN are in the arm category. **(F) Patterns of DTZ for the chr13-chrX translocation.** Top tracks show DTZ of the whole chr13 and the translocated portion of chrX. The middle and bottom tracks show DTZ of chr13 without DTZ of the translocated portion of chrX, explained by the retention of a short centromere proximal portion of chr13 (black oval). Note the strand state of the chrX translocation matches that of the retained portion of chr13. The HAP1^lig4-^ annotation was performed with an SVM model trained with manual annotation from HAP1 cells.

**Figure S26.**
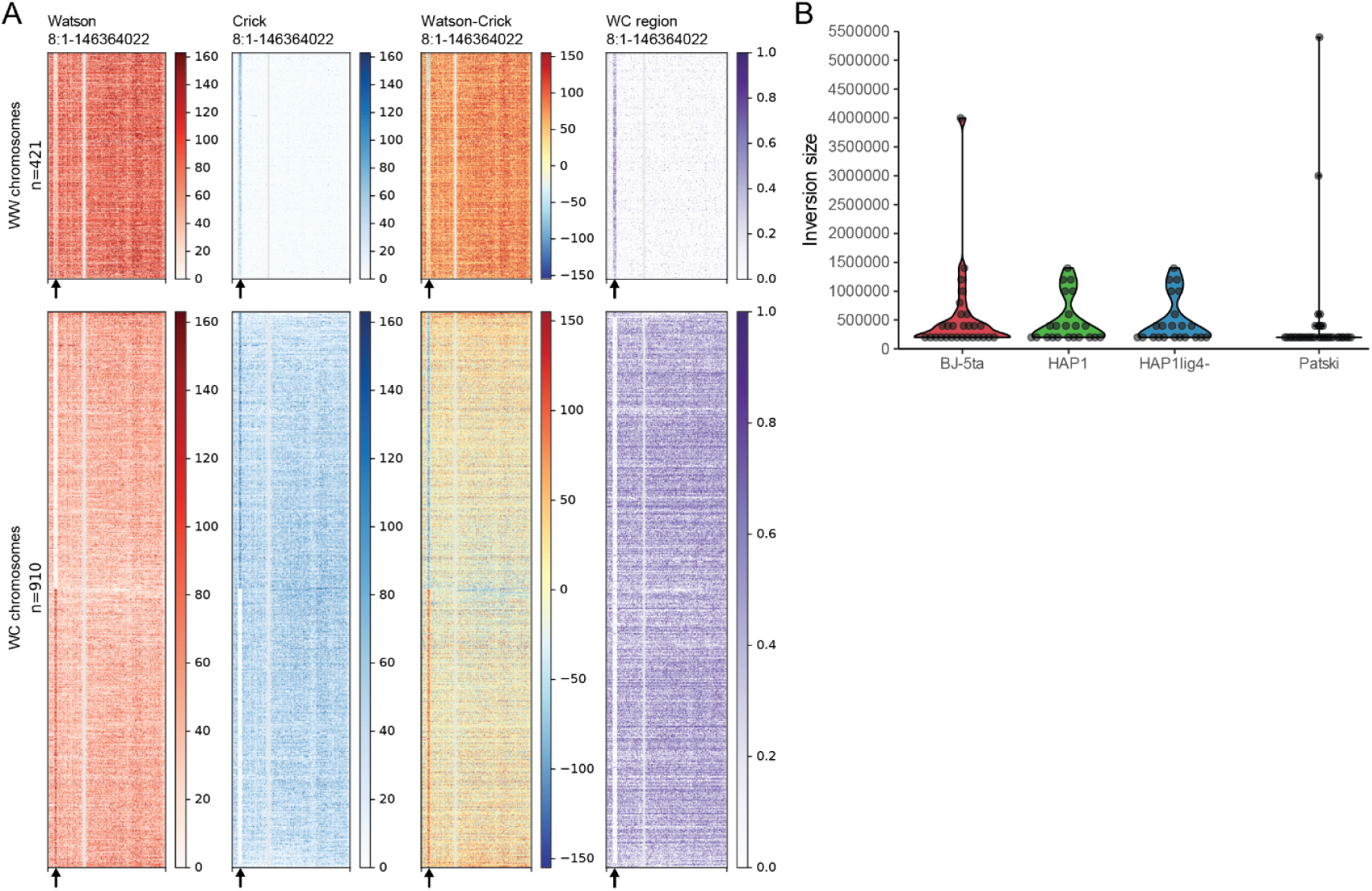
Inversion detection and size distribution. **(A) A universal inversion on chr8 in BJ-5ta shown for WW and WC chromosomes.** Each row of a heatmap represents a single cell plotted at an approximate resolution of 200Kb, sorted by the strand state of a large 4Mb inversion at the start of chr8 (arrow). W reads are plotted on the first column of heatmaps in red (inversion being half red). C reads in blue are plotted on the second column (inversion being half blue). The subtraction of W from C reads is shown on the third column, highlighting WC regions in yellow (W minus C being around 0). The fourth column shows regions where both W and C reads are present. Masked regions are in gray. **(B) Size distribution of inversions**. Inversions were called from data binned at an approximate resolution of 200kb, imposing a 200kb lower limit on inversion call sizes. BJ-5ta median: 299713.5 bp, MAD: 148263.71; HAP1 median: 299713.5 MAD: 148211.82; HAP1^lig4-^ median: 299713.5, MAD: 148211.82; Patski median: 199840, MAD: 85.99.

**Figure S27.**
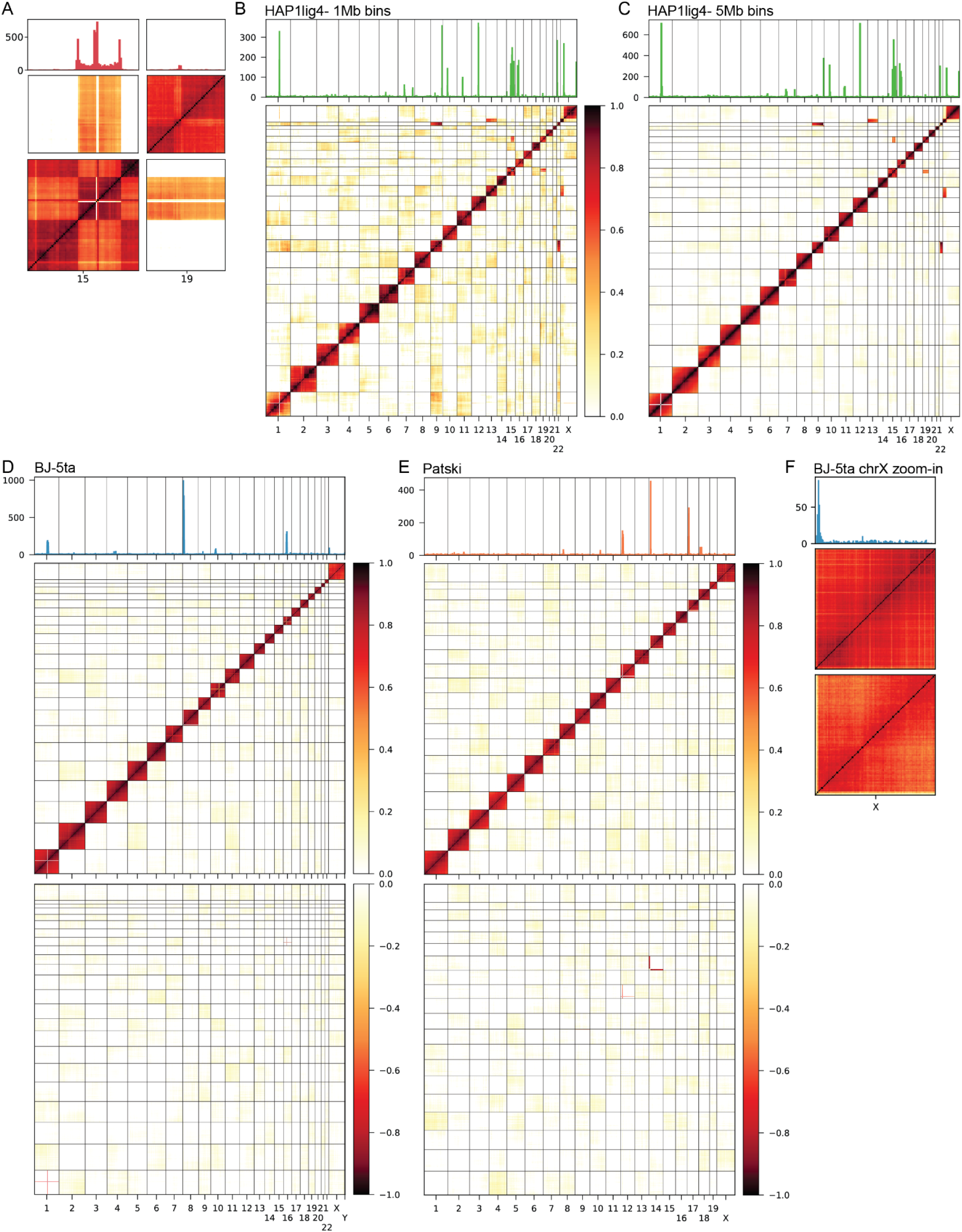
Genome-wide strand-state correlation heatmaps for translocation detection. **(A-F)** Pearson correlation of strand states for pairs of genome-wide bins across hundreds of single cells. Heatmaps were plotted at a 1Mb bin resolution unless stated otherwise. The panel above each heatmap shows breakpoints without the “region filter” that highlight enriched strand switches over translocations. Breakpoints were counted across the same bins as the heatmap. The list of translocations with exact coordinates is provided in TabS2. **(A) A zoom-in of chr15 and chr19 in HAP1 cells.** The strand state of interstitial duplicated region of chr15 correlates with the strand state of chr19, highlighting an insertional translocation. Scale is the same as in (B). **(B) Three highly correlated sites *in trans* were observed for HAP1^lig4-^ cells.** The correlation heatmap identified the two known translocations also present within HAP1 cells. An additional translocation between chr13 and chrX is also observed. High background observed in trans is due to the limited set of 62 cells used. **(C) A lower resolution correlation heatmap using shallow sequenced HAP1^lig4-^ cells.** A total of 689 cells with less than 20 reads per Mb were used to create a 5Mb sized bin correlation heatmap. The same three translocations are observed as in (B). Scale same as in (B). **(D) BJ-5ta cells have no detectable translocations.** The heatmaps were created from 582 cells. The top heatmap shows positive and the bottom shows negative Pearson correlation. Negative Pearson correlation or that close to zero *in cis* corresponds to inversions, e.g., chr1. **(E) Patski cells have no detectable translocations.** The heatmap was created from the strand state correlation across 426 cells (top: positive correlation; bottom: negative. Same as (D)). No translocations were detected and a reference inversion was captured on chr12 and chr14. **(F) A zoom-in of chrX in BJ-5ta cells.** With the full 582 cell complement, the terminal start of chrX shows a correlation decrease to close to zero. The region is a breakpoint hotspot due to a inversional duplication in a large proportion of cells. The bottom heatmap was created with a subset of 397 cells containing the inversional duplication.

**Figure S28.**
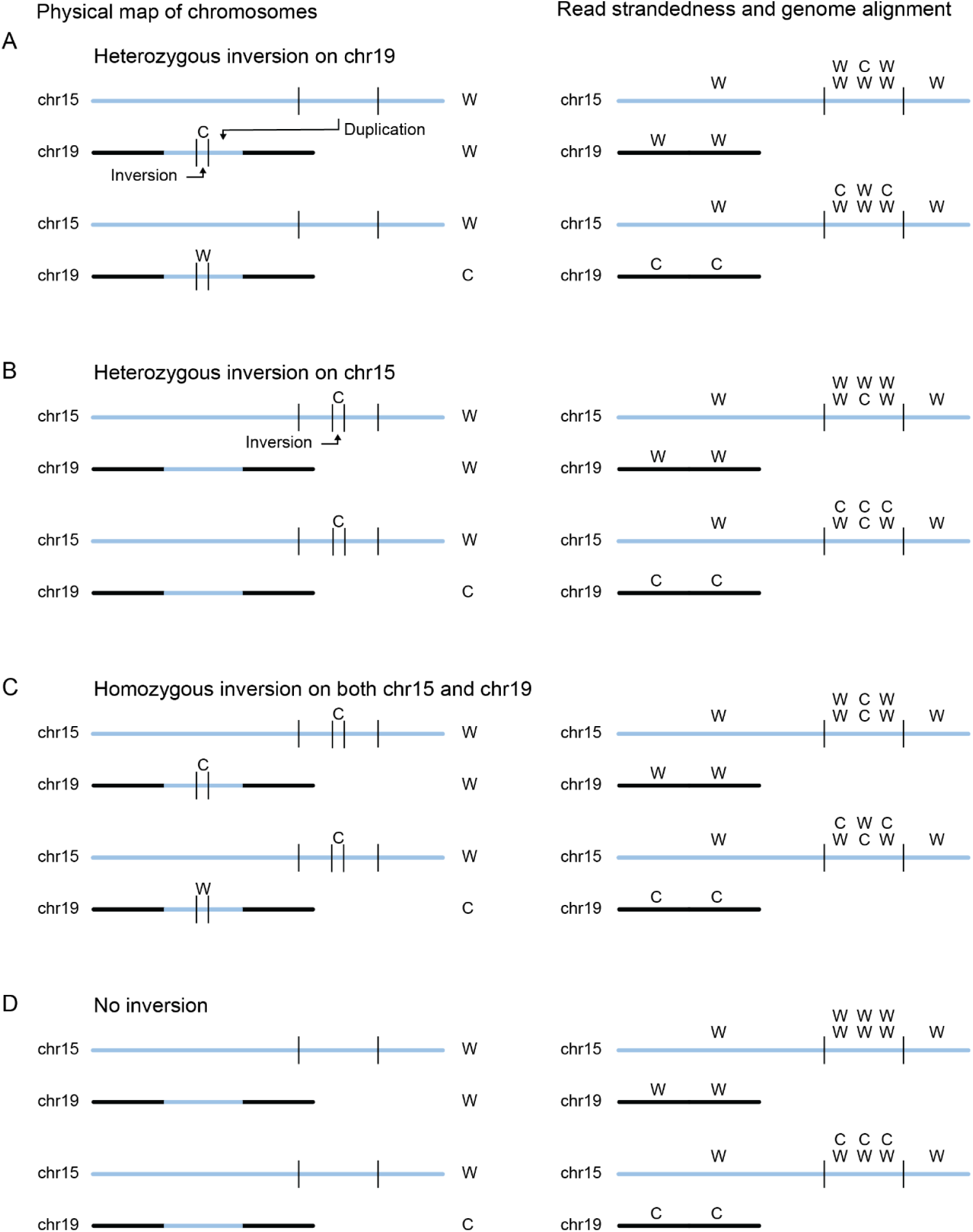
Schematic of all possible scenarios of the inversion within the chr15-chr19 insertional translocation region in the haploid HAP1 cells. **(B) Expected strand segregation for a heterozygous inversion embedded in the duplicated chr15 sequence on chr19.** An inversion within the insertional translocation of chr15 into chr19 results in a C in an otherwise W chromosome (top left) or vice versa (bottom left). In this instance the resulting read alignment would result in W-C-W strand states over the duplicated region for W chr15 and W chr19 (an overall W-WW-WC-WW-W strand state read-out for chr15, and W for chr19, matching the observed heatmap in the lower right quadrant in Fig7B for n=332 cells), or C-W-C for an otherwise W chr15 and C chr19 (an overall W-WC-WW-WC-W strand state read-out for chr15, and C for chr19, right panel, matching the observed heatmap in the upper left quadrant in Fig7B for n=323 cells). C chr15 and C chr19 (matching the other two observed heatmaps) are not shown as they are the opposite of the W chr15 and W chr19 example. **(C) Expected strand segregation for a heterozygous inversion embedded in the chr15 sequence on the original copy of chr15.** If *<u>after</u>* the insertional translocation, an inversion occurred within the region of the original chr15 and not on the duplicated copy (left panel), the resulting read alignment would result in distinct W-C-W strand states for W chr15 and W chr19 or W-C-W for W chr15 and C chr19 (right panel). Patterns in B-D are not observed. **(D) Expected strand segregation for a homozygous inversion on both chr15 and chr19.** Before the insertional translocation, an inversion could occur within chr15 leading to its later presence on both chr15 and chr19 (right panel). In this instance, both copies of chr15 would contain a strand switch. For a W chr15 and a W chr19, the resulting read alignment would produce a W-C-W strand states for both copies, while for a W chr15 and a C chr19 a combination of C-W-C and W-C-W would be observed. **(E) No inversion.** Without an inversion, the strand states of the duplicated region would match the resident chromosomes; the insertional duplication would match the strand state of chr19.

## Notes

### Competing Interest Statement

The authors have declared no competing interest.

### Summary of Updates

FigS1E, Fig2E and FigS10C were added to provide additional detail; Fig7B was revised for clarity.

